# Tumor-targeted delivery of childhood vaccine recall antigens by attenuated *Listeria* reduces pancreatic cancer

**DOI:** 10.1101/600106

**Authors:** Benson Chellakkan Selvanesan, Dinesh Chandra, Wilber Quispe-Tintaya, Arthee Jahangir, Ankur Patel, Kiran Meena, Rodrigo Alberto Alves Da Silva, Madeline Friedman, Steven K Libutti, Ziqiang Yuan, Jenny Li, Sarah Siddiqui, Amanda Beck, Lydia Tesfa, Wade Koba, Jennifer Chuy, John C. McAuliffe, Rojin Jafari, David Entenberg, Yarong Wang, John Condeelis, Vera DesMarais, Vinod Balachandran, Xusheng Zhang, Claudia Gravekamp

## Abstract

Pancreatic ductal adenocarcinoma is highly metastatic, poorly immunogenic, and immune suppression prevents T cell activation in the tumor microenvironment. We developed a microbial-based immunotherapeutic concept for selective delivery of a highly immunogenic tetanus toxoid protein (TT_856-1313_), into tumor cells by attenuated *Listeria monocytogenes*, and reactivation of pre-existing TT-specific memory T cells (generated during childhood) to kill infected tumor cells. Thus, TT here functions as an alternative for neoantigens. Treatment of KPC mice with *Listeria*-TT resulted in TT accumulation in tumors and inside tumor cells, and attraction of predominantly TT-specific memory CD4 T cells. Moreover, gemcitabine (GEM) combined with *Listeria-TT* significantly improved the migration of CD4 T cells into tumors and the production of perforin and granzyme B, turning cold tumors into immunological hot tumors. *In vivo* depletion of T cells in Listeria-TT+GEM-treated mice demonstrated CD4 T cell-mediated eradication of tumors and metastases (Mann-Whitney p<0.05). In addition, peritumoral lymph node like structures (LNS) were observed in close contact with the pancreatic tumors displaying CD4 T cells and CD8 T cells of KPC mice treated with *Listeria*-TT or *Listeria*-TT+GEM. The production of perforin and granzyme B was observed in LNS of KPC mice that received *Listeria*-TT+GEM. This combination not only reduced tumor burden (80%) and metastases (87%) significantly (p<0.05, Mann-Whitney), but also improved the survival time of KPC mice with advanced pancreatic cancer substantially (Mantel-Cox p<0.0001). Our results unveil new mechanisms of *Listeria* and GEM improving immunotherapy for PDAC.

## INTRODUCTION

Pancreatic ductal adenocarcinoma (PDAC) is extremely difficult to cure. Modern systemic therapies, such as gemcitabine, provide some survival benefits^1–4^, while gemcitabine and Abraxane or FOLFIRINOX further modestly improve survival^5,6^. This underscores the need for additional and innovative approaches. Cancer immunotherapy with checkpoint inhibitors has shown promising results, however, less so for PDAC^7^. PDAC is poorly immunogenic because of the low mutational load and the few effective neoantigens present^8,9^, and because immune suppression, particularly by myeloid-derived suppressor cells (MDSC) and tumor-associated macrophages (TAM), prevents T cell activation in the tumor microenvironment (TME)^10–12^. Moreover, evidence suggests that naïve T cells are more prone undergoing apoptosis or are less efficiently activated than memory T cells in the TME of cancer patients and tumor-bearing mice^13,14^.

To address these problems we have developed a novel treatment modality using an attenuated bacterium *Listeria monocytogenes*^15^ (*Listeria*) to deliver a highly immunogenic recall antigen such as tetanus toxoid (TT)(as an alternative to neoantigens) selectively into tumor cells, and to reactivate pre-existing memory T cells against TT (generated during childhood vaccinations) **(Fig. S1A)**. These TT-specific memory cells are in turn attracted to the *Listeria-*TT in the TME, where they destroy the infected now highly immunogenic tumor cells. These memory T cells circulate in the blood for life and can be reactivated at any age. This has been shown even in patients with cancer, including pancreatic cancer^16–18^. Such an approach overcomes the need for participation of naïve T cells during cancer treatment. To further improve T cell responses against TT, *Listeria*-TT was combined with low doses of gemcitabine (GEM) to reduce immune suppressive MDSC and TAM populations.

In previous studies we have shown that, most likely through C3bi and C1q receptors^19–21^, *Listeria* attracts and infects MDSC^10^. These MDSC are present in large numbers in patients and mice with cancer^11,22^. However, the primary tumor also selectively attracts MDSC through the production of cytokines and factors^11,23^, and thus MDSC deliver *Listeria* to the TME as a Trojan horse^10,22^. Once at the tumor site, *Listeria* spreads from MDSC into tumor cells through a cell-to-cell mechanism unique to *Listeria*^24^. *Listeria* can also infect tumor cells directly^25^. *Listeria* is rapidly killed in healthy tissues, but protected from immune clearance in the TME through strong immune suppression. Because of this combination of selective attraction of MDSC by bacteria and cancer with the strong immune suppression occurring in the TME but not in normal tissues, *Listeria* can selectively enter, multiply, and survive in the TME but not in normal tissues^10,22,26,27^. Based on these results we now use *Listeria* as a platform for the selective delivery of anticancer agents to the TME^22,26,28^. Of note is that this approach is completely different from previous clinical trials, in which *Listeria* has been used to infect antigen-presenting cells to stimulate T cells against naturally expressed tumor antigens^29^. In our approach *Listeria* instead colonizes and changes the tumor microenvironment, delivers childhood vaccine recall antigens into tumor cells, and reactivates the pre-existing memory T cells to TT, now killing the infected tumor cells.

In the current study, we tested the combination of *Listeria*-TT+GEM **(Fig. S1B)** in two mouse models of pancreatic cancer, a syngeneic Panc-02 model^30^ and a transgenic KPC model^31,32^. *Listeria*-TT was administered intraperitoneally, and the biodistribution of *Listeria*-TT, as well as the production of TT in tumor cells was monitored *in vivo*. We demonstrate that TT was delivered to the TME and secreted inside and outside tumor cells, attracting TT-specific CD4 T cells producing granzyme B and less abundantly perforin, resulting in a significant reduction of pancreatic cancer, and a significant improvement in survival. We demonstrated that these CD4 T cells and not the CD8 T cells were responsible for eradication of the pancreatic cancer *in vivo*. This study provides new insight into mechanisms of improving immunotherapy for pancreatic cancer.

## RESULTS

### Development and characterization of *Listeria*-TT

The *Listeria* construct used to develop *Listeria*-TT includes pGG34^33^, a truncated non-cytolytic Listeriolysin O (LLO) fused to a non-toxic TT_856-1313_ fragment, and a myc sequence for detection as outlined in **Fig. 1A.** Secretion of LLO-TT_856-1313_ protein into the culture medium by the bacteria was detected by western blotting **(Fig. 1B).** Infection of Panc-02 tumor cells with *Listeria*-TT_856-1313_ resulted in the expression of TT protein in the tumor cells **(Fig. 1C).**

**Fig. 1.**
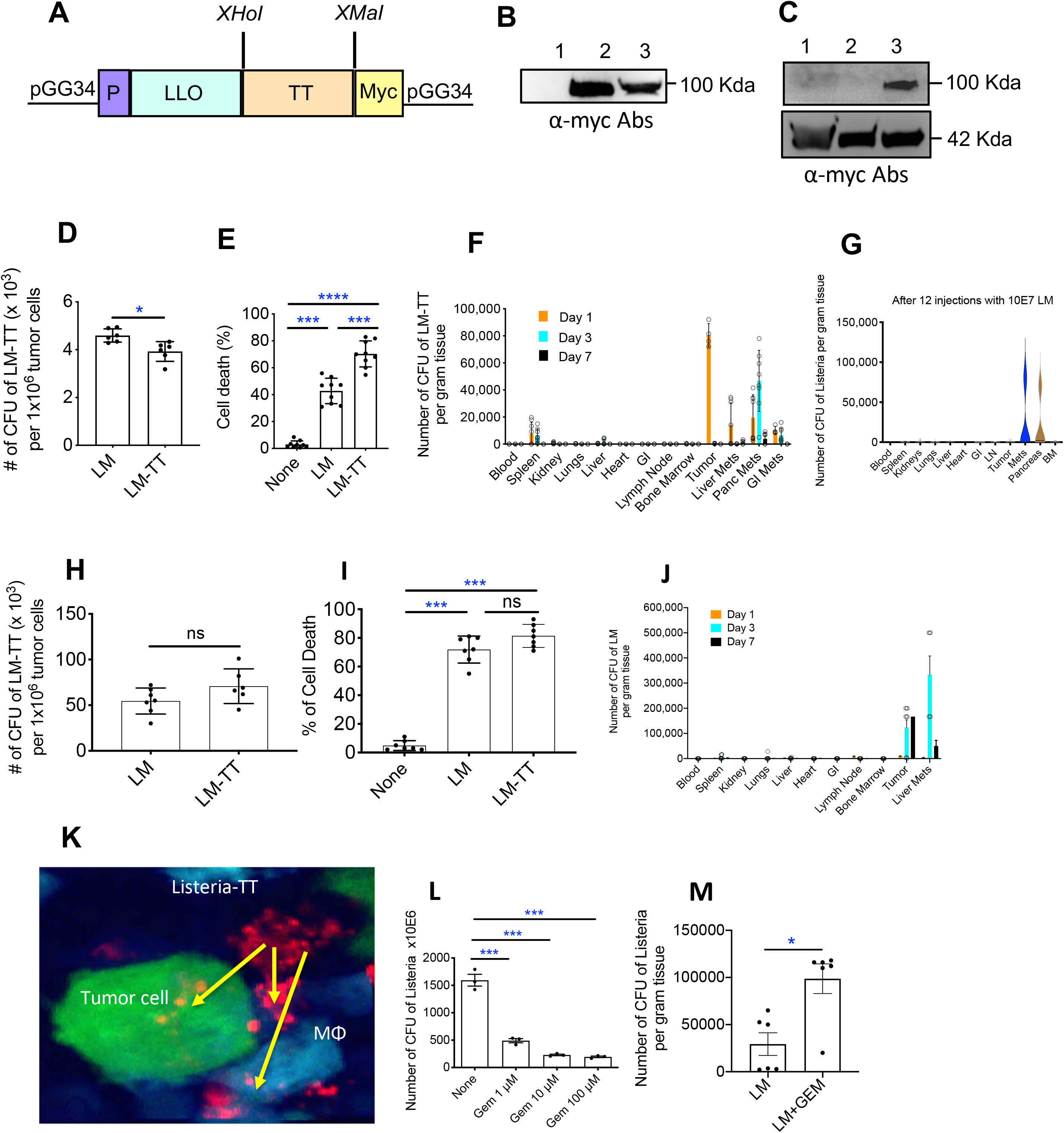
Development and characterization of the *Listeria*-TT vaccine. **(A)** *Listeria*-TT construct. TT_856-1313_, a non-toxic fragment of the C-terminus of TT cDNA (aa position 856-1313, 52 kDa) was cloned as a fusion protein with a truncated non-cytolytic Listeriolysin O (LLO, 48 kDa) into the *Listeria* plasmid pGG34, under the control of the LLO promoter (P). A myc tag was included for detection of the TT protein. **(B)** Western blot detection of TT protein secretion using anti-myc. **Lane 1**: negative control (growth medium);; **Lane 2**: supernatant of *Listeria*-TT culture;; **Lane 3**: pellet of *Listeria*-TT culture. **(C)** Detection of TT protein in tumor cells. **Lane 1**: Panc-02 tumor cells;; **Lane 2**: Panc-02 tumor cells infected with *Listeria* alone;; **Lane 3**: Panc-02 tumor cells infected with *Listeria*-TT. (**D)** Panc-02 tumor cells infected with *Listeria*-TT. Average of 2 independent experiments, 3 wells per group. (**E)** Panc-02 tumor cell killing by *Listeria*-TT. Averages of 3 independent experiments;; 3 wells per group. **(F)** *Listeria*-TT accumulation in tumors and metastases, but not in normal tissues. A single high dose of *Listeria*-TT, 10^7^ CFU/200 μL, was injected ip into Panc-02 mice, and *Listeria*-TT CFUs were measured in tumor, metastatic, and normal tissues at different time points. Averages of a single experiment;; n=3 mice per timepoint, n=3 tissues per organ/tumor/metastases. **(G)** *Listeria* still accumulates in pancreatic tumors and metastases after 12 high doses (10^7^ CFU). **(H)** *Listeria*-TT infects human Mia-PaCa2 pancreatic tumor cells *in vitro*. **(I)** *Listeria*-TT killing human MiaPaCa2 pancreatic tumor cells *in vitro*. **(J)** *Listeria* also accumulates in human pancreatic tumors and metastases *in vivo.* A single high dose of *Listeria*-TT, 5×10^7^ CFU/200 μL, was injected ip in nude mice with MiaPaCa2 tumors in the pancreas, and *Listeria*-TT CFUs were measured in tumor, metastatic, and normal tissues at different time points. Averages of a single experiment;; n=3 mice per timepoint, n=3 tissues per organ/tumor/metastases. **(K)** *Listeria*-TT infecting tumor cells and macrophages (MΦ) in Panc-02-dendra-2 tumors of live mice, by intravital imaging. Tumor cells are green, macrophages blue, and *Listeria*-TT red. **(L)** Effect of GEM on *Listeria in vitro*. Average of one experiment;; n=3 wells per concentration, 3 fields per well. **(M)** Effect of GEM on *Listeria in vivo*. Average of 2 independent experiments;; n=3 mice per group. Mann-Whitney *p <0.05, ***p<0.001, ****p<0.0001. LM-TT = *Listeria*-TT. The error bars represent SEM. In E and I, all groups were compared to untreated (none). In L, all groups were compared to untreated *Listeria* (none).

### *Listeria*-TT infection of tumor cells *in vitro* and *in vivo*

We previously showed that *Listeria* can be used as a delivery platform for anticancer agents^22,26^, however TT fusion to LLO could potentially alter LLO function (LLO is required to escape the vacuole after phagocytosis^34^). To test *Listeria*-TT delivery function, we determined the infection rate of pancreatic tumor cells cultured with *Listeria*-TT in comparison to *Listeria* alone. Panc-02 tumor cells were infected in both cases **(Fig. 1D).** We also determined that *Listeria*-TT effectively killed the Panc-02 cells **(Fig. 1E)**. Of note is that the infection time *in vitro* is 2hrs, while *in vivo* a complete treatment cycle takes 14 days. Thus, *Listeria* has significantly more time to infect tumor cells *in vivo* than *in vitro*. Indeed, confocal microscopy shows *Listeria* spread all over the pancreatic tumor **(Fig. S2).** We also have shown this in multiple other studies in KPC and Panc-02 mice^22,26^.

TT is highly immunogenic and might lead to faster elimination of *Listeria*-TT *in vivo* than *Listeria* alone, consequently preventing accumulation in the TME. To assess this, a single high dose of *Listeria*-TT was injected intraperitoneally (ip) into Panc-02 mice, and numbers of *Listeria*-TT were quantified in all tissues (tumors, metastases, and normal tissues) at different time points. As shown in **Fig. 1F and Table S1**, on day 1 after injection, the CFU of *Listeria*-TT was 10-fold higher in the primary tumor than in the spleen, while this was true for metastases in the pancreas (7.5-fold) on day 3, and on day 7 no *Listeria*-TT bacteria were found in the spleen or other normal tissues while *Listeria*-TT was still detectable in liver and pancreas metastases (1000 and 3875 CFU per gram tissue, respectively). In summary, *Listeria*-TT accumulated more abundantly in tumors and metastases than in spleen or other normal tissues on days 1 and 3 after injection of the *Listeria*-TT, while on day 7 *Listeria*-TT was completely absent in normal tissues but still detectable in tumors and metastases. Also important is that *Listeria* even after 12 high doses is still able to colonize the immune suppressive tumor microenvironment but is eliminated faster in normal tissues that lack immune suppression **(Fig. 1G).**

Interestingly, the same dose of *Listeria* injected intravenously lead to nearly undetectable numbers of CFU of *Listeria* in tumors and metastases **(Fig. S3A and B).** This was true not only for metastases in the liver and pancreas, but also for metastases in the lungs **(Fig. S3C).**

We also evaluated the infection and kill rate of human pancreatic tumor cells by *Listeria*. The infection rate of human pancreatic tumor cells Mia-PaCa2 **(Fig. 1H)** by *Listeria* and *Listeria*-TT was 5-10-fold higher than of the mouse pancreatic tumor cells **(Fig. 1D)**, while the kill rate overnight was similar **(Figs. 1E and I).** In a previous study we also found that also human breast tumor cells were more efficiently infected by *Listeria in vitro* than mouse breast tumor cells^25^. To evaluate the clinical application of *Listeria*, we determined the biodistribution of *Listeria* in nude mice with human pancreatic tumors. For this purpose, 10^6^ human pancreatic tumor cells (Mia-PaCa2) were injected orthotopically in the pancreas of nude mice, and 6 weeks later when tumors were palpable (tumors were 1 cm in diameter), a single high dose of *Listeria* was injected ip, and the numbers of *Listeria* were quantified in all tissues at different time points as described above. Of note is that when 1×0^7^ CFU of *Listeria* was injected, the bacteria were hardly detected in tumors, metastases and normal tissues (data not shown). Also, others reported that less CFU of *Listeria* was observed after injection in nude than in wildtype mice^35^. Therefore, we increased the dose to 5×10^7^ CFU of *Listeria*, which resulted in a 5-time higher number of CFU of *Listeria* in the human pancreatic tumor **(Fig. 1J)** compared to the mouse tumors **(Fig. 1F).** Although on day 1, the number of CFU of *Listeria* was very low in tumors and metastases (8.3×10^3^ and 3.3×10^3^ per gram tissue, respectively), on day 3 the number of CFU was high particularly in metastases (average 3.3×10^5^ per gram tissue), and even on day 7, there were still high numbers of *Listeria* in the primary tumor (1.6×10^5^ CFU of *Listeria* per gram tissue) and metastases (5×10^4^ CFU per gram tissue) **(Fig. 1J and Table S2).** Thus, the pattern of biodistribution in human tumors was somewhat different from mouse tumors. We believe that the immune system in nude mice play a role here. Nude mice do not have T cells, and allow the *Listeria* to survive longer, while the innate immune system in nude mice is more efficient in eliminating the bulk of *Listeria* in the early stage of infection. In summary, we have shown here that *Listeria* also efficiently accumulate in human tumors.

To confirm that *Listeria*-TT was taken up *in vivo* by tumor cells within the TME of live mice, the *Listeria*-TT was incubated with anti-*Listeria* polyclonal antiserum and anti-IgG-Alexa-680 antibodies. Subsequently, the Alexa-680-labeled *Listeria*-TT was then injected into the peritoneal cavity of transgenic mice whose macrophages expressed cyan fluorescent protein (CFP)^36^, and which had orthotopic, Dendra-2 labeled pancreatic tumors (Panc-02-Dendra). High-resolution intravital multiphoton microscopy (IVMI) of these tumors showed individual *Listeria*-TT bacteria inside tumor cells and macrophages (**Fig. 1K**). Outside the tumor cells aggregates of *Listeria* were observed. The polyclonal antiserum against *Listeria* contained agglutinating antibodies and were responsible for the aggregated *Listeria* observed outside the cells in **Fig. 1K.** We also analyzed *Listeria* in the TME through IHC by confocal microscopy using anti-*Listeria* antibodies and showed that many *Listeria* were present in tumors and even more in metastases **(Fig. S2)**.

One of the hallmarks of our *Listeria* is that they continuously spread from cell to cell and that TT protein is secreted in an outside the tumor cells. To determine how much of a tumor expressed TT, we performed IHC on tumors of KPC mice that received a complete treatment cycle. As shown in **Fig. 2A**, TT was secreted in about 80% of the primary tumor. This included TT inside tumor cells and most likely macrophages. Since *Listeria* continuously spreads from one cell to another, and simultaneously secretes TT in and outside cells, detection of TT itself instead of the number of *Listeria* per tumor cell, is more reliable to measure the success of delivery. Therefore, we now use TT as measurement of success of delivery. We also analyzed all 4 treatment groups by TT staining, and as expected TT was only detected in tissues of KPC mice treated with *Listeria*-TT or *Listeria*-TT+GEM **(Fig. 2B).**

**Fig. 2:**
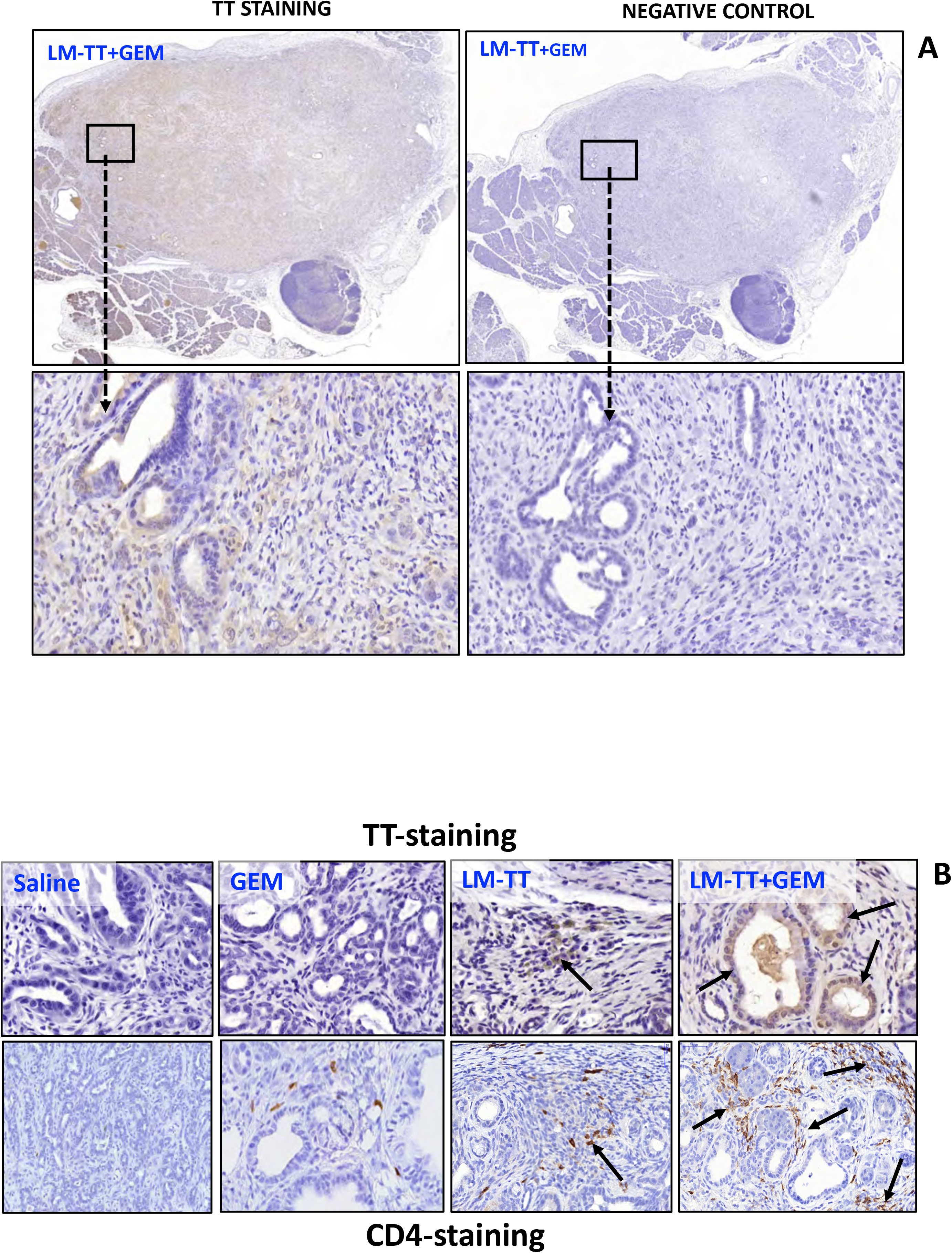
Delivery of TT by *Listeria* in pancreatic tumors. **(A)** KPC mice received a complete treatment cycle with *Listeria*-TT+GEM as outlined in Fig S1B. Two days after the last treatment tumors were dissected and processed for IHC staining using anti-TT antibodies. As negative control the secondary antibody (anti-IgG-HRP conjugated) only was used. TT expression was observed in about 80% of the KPC tumor, while no TT expression was observed in the negative control. **(B)** TT expression and CD4 T cells were analyzed in KPC tumors by IHC of *Listeria*-TT+GEM-treated and control mice (saline, GEM, LM-TT). TT expression was observed in KPC tumors of mice treated with *Listeria*-TT or *Listeria*-TT+GEM only. TT was observed in and outside tumor cells. CD4 T cells were present in the areas where TT was highly expressed, i.e. around the neoplastic pancreatic ducts. LM-TT = *Listeria*-TT, GEM = gemcitabine.

Notably, detection of TT was stronger in the tumors of *Listeria*-TT+GEM-treated mice than of *Listeria*-TT-treated mice. While GEM kills *Listeria in vitro*, more *Listeria* bacteria were detected in the tumors of *Listeria*+GEM-treated mice compared to *Listeria* alone **(Figs. 1L and M).** Since GEM reduces immune suppression, the extracellular *Listeria* will be more exposed to immune selective pressure which may drive *Listeria* to an area with better survival, i.e. to the TME and inside tumor cells. However, it is also possible that GEM+*Listeria* induced more necrosis than *Listeria* alone. *Listeria* survives better in necrotic areas, which are more hypoxic^37^.

Based on TT expression in the tumors **(Fig. 2AB)**, it seems that *Listeria* more abundantly accumulated around the pancreatic ducts in the KPC tumors. It has been reported that phospholipase A2 and B are produced in the pancreatic duct in bile products, normally to digest food (https://www.studyread.com/pancreatic-enzymes/). However, *Listeria* survives better in an environment with phospholipase A2 or B, because its critical for the spread of *Listeria*^38,39^.

### *Listeria*-TT reactivates memory T cells to TT in tumor-bearing mice

The elderly lack naïve T cells^40^ and most cancer patients are old. Therefore, we expect that cancer immunotherapy might be more effective if the need for naïve T cells could be avoided during cancer treatment^41^. To address this point, we generated TT-specific memory CD4 and CD8 T cells in mice through two injections of Tetanus vaccine (TTvac) prior to tumor development **(Fig. 3A)**, similar to the TT vaccinations that have been used in humans to generate memory T cells and B cells to TT. Once the tumors and metastases were detected by PETscan, we started with a complete treatment cycle with *Listeria*-TT+GEM as outlined in **Fig. S1B.** After the complete treatment cycle, spleen cells were isolated from the KPC tumor-bearing mice, restimulated with TT protein *in vitro*, and analyzed for memory T cell responses producing IFNγ **(Fig. 3B)**. We found that CD4 and CD8 T cells of mice that received *Listeria* -TT or *Listeria* -TT+GEM were strongly activated by TT, but not of mice that received saline or GEM alone (Mann-Whitney p<0.05 and p<0.001), indicating the high specificity of CD4 and CD8 memory T cells responsive to the TT protein. Similar CD4 and CD8 T cell responses were observed in the spleen of Panc-02 mice **(Fig. S3D).**

**Fig. 3.**
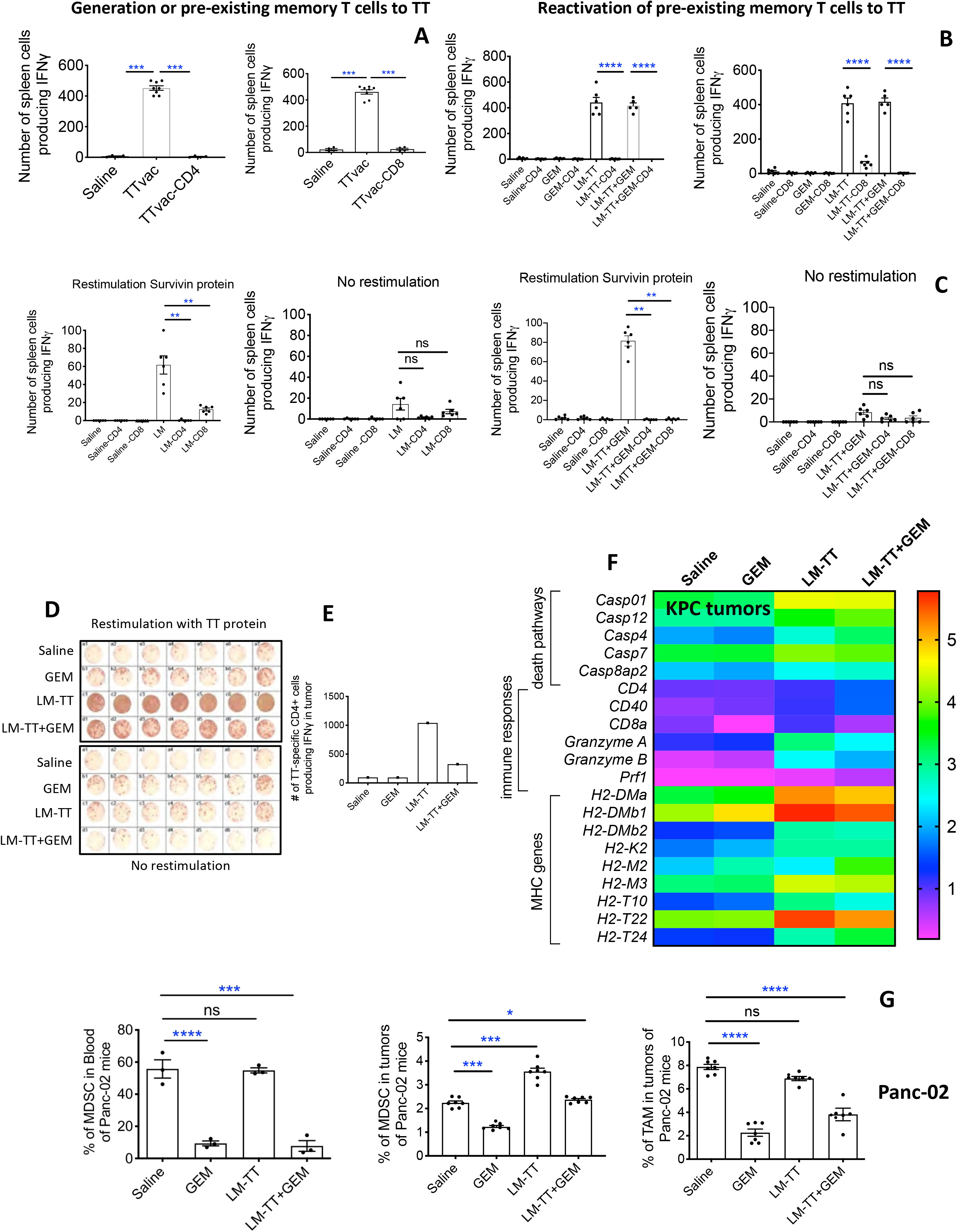
Generation and recall of memory CD4 and CD8 T cell responses to TT. **(A)** Generation of CD4 and CD8 memory T cells to TT by immunization of mice with Tetanus vaccine (TTvac), prior to tumor development. **(B)** Reactivation of TT-specific memory CD4 and CD8 T cells in KPC mice post tumor development through the *Listeria*-TT+GEM treatment cycle described in Fig S1B. **(C)** *Listeria* induces immunogenic tumor cell death. C57Bl/6 mice with Panc-02 tumors received one high and multiple low doses of *Listeria* alone as described in fig S1B. Spleen cells were isolated and analyzed for T cell responses to *Survivin* protein by ELISPOT. A similar experiment was performed in KPC mice with pancreatic tumors, but now received *Listeria*-TT+GEM. Representative results of 2 experiments are shown;; n=3 mice per group (pooled), 6 wells per group. Statistical significance by Mann-Whitney test: *p<0.05, **p<0.01 and ****p<0.0001. **(D)** ELISPOT analysis of CD45^+^ IFNγ-producing immune cells isolated from orthotopic Panc-02 tumors in mice treated with *Listeria*-TT+GEM. **(E)** Flow cytometry of IFNγ-producing CD4^+^ T cells in the CD45^+^ fraction, isolated from orthotopic Panc-02 tumors. In each group tumors of 5 mice were pooled. Mann-Whitney *p <0.05, **p<0.01, ***p<0.001, ****p<0.0001. **(F)** RNAseq analysis of immune responses and tumor cell death pathways in pancreatic tumors of KPC mice in response to *Listeria*-TT+GEM treatment and control groups. The results of 2 mice in each group was averaged. The heatmap was generated by unbiased hieratical clustering at the following statistical parameters. Statistical parameters for the paired-wise comparison: p<0.05. The signature contains 20 important genes. **(G)** Flow cytometric analysis of changes in MDSC (CD11b^+^Gr1^+^) in blood and tumors, and TAM (CD11b^+^F4/80^+^) in tumors of Panc-02 mice in response to the *Listeria*-TT+GEM treatment cycle. Average of a single experiment of MDSC in blood with n=3 mice per group, and the average of 2 independent experiments of MDSC and TAM in tumors with n=3 mice per group. LM-TT = *Listeria*-TT, GEM = gemcitabine. The error bars represent SEM. In A, all groups were compared to the TTvac group. In A, all groups were compared to TTvac. In B and C, T cell-depleted groups were compared to non-depleted groups. In G, all groups were compared to the saline group.

Other mechanism(s) of tumor cell destruction has been analyzed. In a previous study on breast cancer we have shown that *Listeria*-generated reactive oxygen species (ROS) induces immunogenic tumor cell death resulting in T cell responses to tumor-associated antigens (TAA)^14^. In the current study we demonstrate by ELISPOT that *Listeria* generates CD4 and CD8 T cell responses to *Survivin* **(Fig. 3C)**, a TAA that is expressed by Panc-02 and KPC mice^42,43^. Therefore, non-infected tumor cells can be destroyed by TAA-specific T cells. TAA are less immunogenic than TT and their T cell responses are less vigorous compared to TT-specific T cells **(Fig. 3BC).**

To evaluate whether *Listeria*-TT+GEM could overcome immune suppression in the tumor-bearing KPC mice, we also analyzed “*in vivo*” T cell responses, i.e., without *in vitro* TT-restimulation. At the end of the treatment cycle, we found that CD4 and CD8 T cell responses (CD69, granzyme B, and perforin) were strongly improved in spleen cell populations (pooled) of mice treated with *Listeria*-TT+GEM compared to all control groups **(Table 1A).** Both CD4 and CD8 T cells produced high levels of granzyme B and perforin, while the production of IFNγ was variable. *Listeria* or GEM alone also increased the production of perforin in CD4 and CD8 T cells compared to the saline treatments. Apparently, this has also been reported by others^44^. Similar results were found in Panc-02 mice, i.e., *Listeria*-TT+GEM improved T cell responses compared to all other groups **(Table 1B and Fig. S4AB).** In summary, the T cell responses in **Table 1** are the combination of T cell responses to TT, *Listeria* and GEM-activated T cell responses *in vivo*. This is different from the restimulation assay with TT *in vitro* **(Fig. S3D)**, which shows the production of IFNγ of TT-specific memory T cells by ELISPOT from mice that received TTvac and/or Listeria-TT treatments, underlining the specificity of the TT restimulation assay.

**Table 1A:**
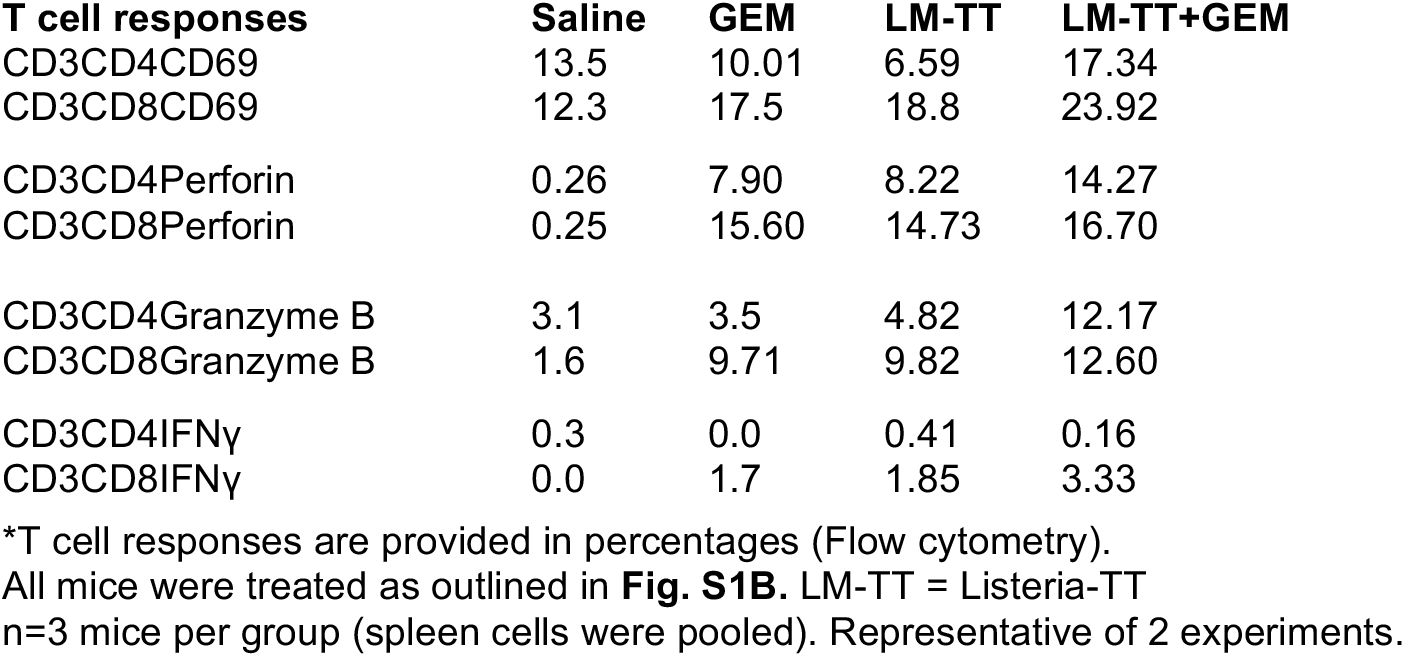
Listeria-TT and GEM increase CD4 and CD8 T cell responses *in vivo* in Panc-02 mice

**Table 1B:**
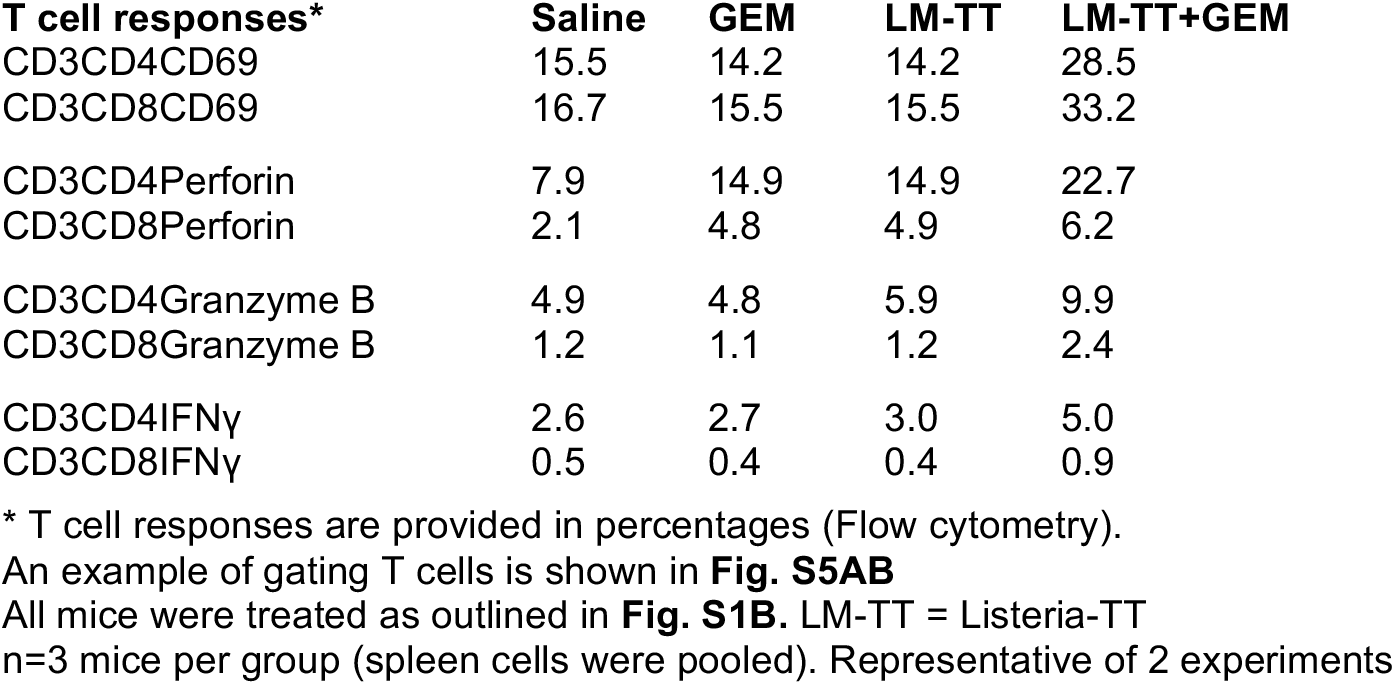
Listeria-TT and GEM increase CD4 and CD8 T cell responses *in vivo* in KPC mice

| T cell responses* | Saline | GEM | LM-TT | LM-TT+GEM |
| --- | --- | --- | --- | --- |
| CD3CD4CD69 | 15.5 | 14.2 | 14.2 | 28.5 |
| CD3CD8CD69 | 16.7 | 15.5 | 15.5 | 33.2 |
| CD3CD4Perforin | 7.9 | 14.9 | 14.9 | 22.7 |
| CD3CD8Perforin | 2.1 | 4.8 | 4.9 | 6.2 |
| CD3CD4Granzyme B | 4.9 | 4.8 | 5.9 | 9.9 |
| CD3CD8Granzyme B | 1.2 | 1.1 | 1.2 | 2.4 |
| CD3CD4IFNγ | 2.6 | 2.7 | 3.0 | 5.0 |
| CD3CD8IFNγ | 0.5 | 0.4 | 0.4 | 0.9 |

\* T cell responses are provided in percentages (Flow cytometry).
An example of gating T cells is shown in **Fig. S5AB**All mice were treated as outlined in **Fig. S1B**. LM-TT = Listeria-TT
n=3 mice per group (spleen cells were pooled). Representative of 2 experiments

### *Listeria*-TT+GEM turns “cold” tumors into immunological “hot” tumors

Based on the fact that *Listeria* causes TT accumulation in the TME **(Fig. 2AB)** and TT-specific memory T cells have been generated **(Fig. 3B)**, we expect these memory T cells to be attracted to the TME. To visualize the TME and T cells in more detail, the KPC tumors were analyzed by immunohistochemistry (IHC). As shown in **Fig. 4A and E**, CD4 T cells were present in the tumors of KPC mice treated with LM-TT, but hardly at all in the tumors of saline- or GEM-treated mice. More detail about the CD4 T cells in the KPC tumors of all four treatment groups is shown in **Fig. S5A-D.** Most remarkable was that addition of GEM to the *Listeria*-TT treatment significantly enhanced the migration of CD4 T cells to the tumors of KPC mice compared to all other groups, even compared to the *Listeria*-TT group. These CD4 T cells were found in the tumor areas that expressed TT protein and CD31-positive vessels **(Fig. 2B and Fig. S6A-C).** Of note is, that CD8 T cells were sparsely present in the tumor areas **(Fig. S5E-H).** Of clinical relevance is to show that these T cells are functional, i.e. that perforin and granzyme B, both involved in T cell-mediated tumor cell cytolysis, are present. Here we demonstrate that both granzyme B and perforin, were detected in the pancreatic tumors of *Listeria*-TT- or *Listeria*-TT+GEM-treated mice by IHC, but not in the tumors of saline or GEM-treated mice **(Fig. 4CE)**. Also, L-selectin as well as Cxc3 and chemokines Cxcl9 and Cxcl10 were highly upregulated in KPC tumors of mice treated with *Listeria*-TT+GEM compared to untreated mice **(Fig. S7A)**. These genes are known to be involved in T cell trafficking^45,46^. Also, CD40 and CD40L was highly upregulated in KPC tumors by *Listeria*-TT+GEM **(Fig. S7A).** It has been reported that CD40 engagement activates T cells and correlates with antitumor activity^47^.

**Fig. 4.**
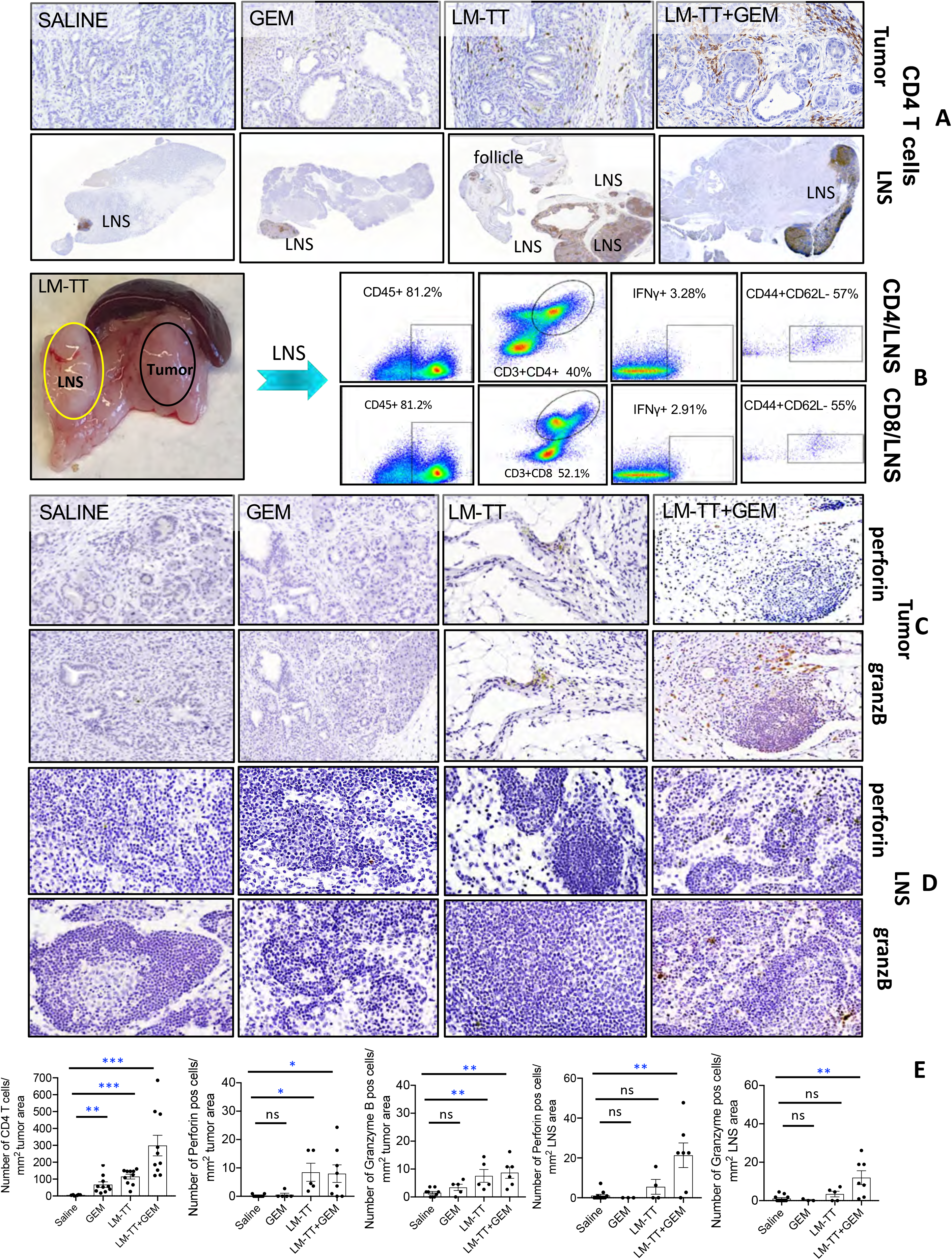
Tetanus protein attracts CD4 memory T cells to the TME. KPC mice were treated with saline, GEM, *Listeria*-TT, or *Listeria*-TT+GEM as outlined in Fig. S1B. Three days after the last treatment, the pancreas with tumors including lymph node like structures (LNS) were excised and processed for CD4 and CD8 staining by IHC. **(A)** CD4 T cells in the KPC tumors and the presence of LNS were analyzed in all treatment groups. **(B)** Isolation and flow cytometry analysis of CD4 and CD8 T cells in a LNS of a mouse that received *Listeria*-TT. The CD45+ cells were gated, followed by CD3CD4 or CD3CD8 gating, followed by gating of CD3CD4 and CD3CD8 T cells producing intracellular IFNγ, followed by staining to identify T cells with memory phenotype (CD44^+^CD62L^-^). **(C)** Production of perforin and granzyme B in KPC tumors, by IHC. **(D)** Production of perforin and granzyme B in KPC LNS, by IHC. **(E)** Quantification of CD4 T cells in KPC tumors and the production of perforin and granzyme B in the KPC tumors and LNS. CD4 T cell numbers were counted in the tumor areas of all treatment groups. Also, the number of perforin and granzyme B-producing cells were counted in the tumor areas and LNS of all treatment groups. In each group, four-ten fields were measured, and the number of cells per 1mm^2^ was calculated. Results of three mice per group were averaged. Mann-Whitney *p<0.05, ** p<0.01, ***p<0.001. LM-TT = *Listeria*-TT, GEM = gemcitabine. Scale bars=100μm. In E, all groups were compared to the saline group.

We also analyzed whether T cells in pancreatic tumors of mice treated with *Listeria*-TT or *Listeria*-TT+GEM were specific for TT. For this purpose, CD45^+^ cells were isolated from the tumors of all four treatment groups, restimulated with TT and then analyzed for the production of IFNγ by ELISPOT **(Fig. 3D)** and for the number of CD4 T cells producing IFNγ by flow cytometry **(Fig. 3E).** The production of IFNγ was 10-fold higher in the tumors of *Listeria*-TT-treated mice compared to the saline or GEM control groups, and 3-fold higher than *Listeria*-TT+GEM **(Fig. 3DE).** This latter was kind of a surprise since the effect of *Listeria*-TT+GEM on tumors and metastases was stronger in KPC and Panc-02 mice than of *Listeria-TT* alone. In a previous study we found that Listeria induces IFNγ in macrophages^17^. Eliminating macrophages by GEM may be an explanation for this lower production of IFNγ production in tumors of LM-TT+GEM-treated compared to LM-TT-treated mice. Also, IFNγ production *in vitro* may not be the best parameter for success of tumor eradication *in vivo*. In contrast, perforin and granzyme B are known to be responsible for tumor cell killing. We observed that GEM improves the production of perforin and granzyme B in the LNS (as shown by IHC) of LM-TT-treated KPC mice **(Fig. 4DE)**, and improved the migration of CD4 T cells into the KPC tumors **(Figs. 4A and E).** This correlated with improved efficacy and survival of the KPC mice.

To confirm the IHC, we performed RNAseq analyses of pancreatic tumors of KPC mice of Listeria-TT+GEM treated and control groups. As shown in **Fig. 3F**, the expression of CD4 but not CD8 genes were upregulated in the tumors of *Listeria*-TT+GEM-treated KPC mice. Moreover, multiple granzymes were expressed (A and B), while perforin was less abundantly expressed, in the tumors of *Listeria*-TT+GEM- and *Listeria*-TT-treated mice. Also, genes involved in tumor cell apoptotic pathways, including multiple caspases, TNF and Fas were upregulated, most likely activated by perforin and granzyme B. Of note also is that MHC class II genes were strongly upregulated in the tumors of KPC mice treated with *Listeria*-TT+GEM or *Listeria*-TT alone **(Fig. 3F).** This was confirmed by IHC **(Fig. S8).**

Strikingly, lymph node-like structures (LNS) were frequently observed in close contact with the KPC tumors of mice treated with *Listeria*-TT or *Listeria*-TT+GEM, and less frequently and in a much smaller format in KPC mice treated with GEM, and hardly in saline-treated mice **(Fig. 4A)**. Higher magnification images of a LNS are shown in **Fig. S6D.** A significant increase in the production of perforin and granzyme B was observed in the LNS of *Listeria*-TT+GEM-treated mice compared to the saline control group **(Fig. 4BE and Fig. S6E-H).** We also analyzed the T cells in the LNS for the production of IFNγ and memory phenotype by flow cytometry. Briefly, the CD45^+^ lymphocytes (81.2%) from the LNS of *Listeria*-TT-treated mice were gated, followed by gating of CD4 (40.4%) or CD8 (52.0%) T cells, followed by gating of the IFNγ+ CD4 (3.28%) or CD8 (2.91%) T cells **(Fig. 4B).** Within these IFNγ+ CD4 and IFNγ+ CD8 T cell populations, 54% and 55%, respectively, exhibited a memory phenotype (CD44+CD62L-). This was supported by RNAscope showing both CD4 and CD8 T cells expressing IFNγ **(Fig. S9)** in the LNS of *Listeria*-TT- and *Listeria*-TT+GEM-treated KPC mice.

In addition, we compared the RNAseq data of KPC tumors with and without LNS for immune function. The difference in expression levels of immune-related genes between the saline and *Listeria*-TT+GEM group was much larger when LNS were present **(Fig. S10)**, compared to the saline and *Listeria*-TT+GEM group of tumors without LNS **(Fig. 3F).** This suggests the importance of LNS for immune function in the TME of KPC tumors. IHC showed that the production of perforin and granzyme B was significantly higher in the LNS of *Listeria*-TT+GEM than in the LNS of saline **(Fig. 4DE)**, supporting the RNAseq data of tumors including the LNS **(Fig. S10).**

Together, the results indicated that *Listeria*-TT+GEM treatment altered “cold” tumors to “hot” tumors, promoting a strong influx of CD4 T cells into the KPC tumors, as well as production of perforin and granzyme B, and that LNS may have played a role here.

### *Listeria*-TT and GEM alter the TME

In Panc-02 tumor bearing mice, GEM reduced the blood MDSC population by 80-90%, and in primary tumors by 50%, compared to the saline group **(Fig. 3G)**, and the TAM population by 67% in primary tumors compared to the saline group **(Fig. 3G).** Moreover, for the residual MDSC and TAM cells, the production in pancreatic metastases of factors involved in immune suppression, including IL-10, IL-6 and the transcription factor MARCO, was reduced in mice treated with *Listeria*-TT+GEM **(Table 1CD, Fig. S11AB)**, while there was increased production by these cells of TNFα, involved in tumor cell killing, and expression of CD80, involved in T cell activation. These findings indicated that *Listeria* TT+GEM altered the TME.

**Table 1c:**
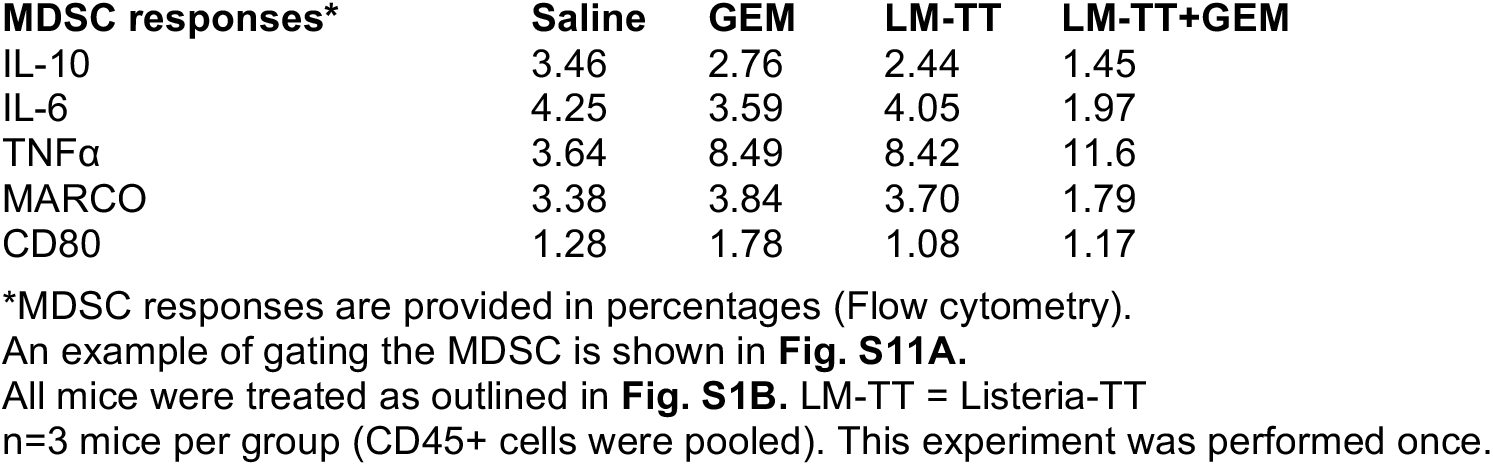
Listeria-TT and GEM reduce immune suppressive function of MDSC (CD11b+Gr1+) in metastases of Panc-02 mice

| MDSC responses* | Saline | GEM | LM-TT | LM-TT+GEM |
| --- | --- | --- | --- | --- |
| IL-10 | 3.46 | 2.76 | 2.44 | 1.45 |
| IL-6 | 4.25 | 3.59 | 4.05 | 1.97 |
| TNFα | 3.64 | 8.49 | 8.42 | 11.6 |
| MARCO | 3.38 | 3.84 | 3.70 | 1.79 |
| CD80 | 1.28 | 1.78 | 1.08 | 1.17 |

\*MDSC responses are provided in percentages (Flow cytometry).
An example of gating the MDSC is shown in **Fig. S11A**.All mice were treated as outlined in **Fig. S1B**. LM-TT = Listeria-TT
n=3 mice per group (CD45+ cells were pooled). This experiment was performed once.

**Table 1D:** Listeria-TT and GEM reduce immune suppressive function of TAM (CD11b+F4/80+) in metastases of Panc-02 mice

| TAM responses* | Saline | GEM | LM-TT | LM-TT+GEM |
| --- | --- | --- | --- | --- |
| IL-10 | 1.45 | 1.47 | 0.65 | 0.72 |
| IL-6 | 1.09 | 1.25 | 0.76 | 0.42 |
| TNFα | 4.78 | 8.85 | 8.73 | 8.04 |
| MARCO | 1.96 | 1.94 | 2.16 | 1.33 |
| CD80 | 1.93 | 4.28 | 3.07 | 5.26 |

\*TAM responses are provided in percentages (Flow cytometry).
An example of gating TAM is shown in **Fig. S11B**.All mice were treated as outlined in **Fig. S1B**. LM-TT = Listeria-TT
n=3 mice per group (CD45+ cells were pooled). This experiment was performed once.

### *Listeria*-TT+GEM reduces advanced and early pancreatic cancer in KPC and Panc-02 mice and improves survival

We evaluated *Listeria*-TT+GEM treatment of 3.5-5 months old KPC mice with pancreatic cancer by PET scan analysis. SUVmax, a measure of the metabolic activity of tumors and metastases, provided an indicator of tumor growth. Significant reductions in SUVmax for PDAC tumors from 1.9 to 0.4, and for liver metastases from 2.1 to 0.3, were observed in the mice receiving *Listeria* TT+GEM treatment **(Fig. 5A)**. The PDAC tumors were reduced by 80% (p<0.01) and the liver metastases by 87% (p<0.01). Small changes in the SUVmax of tumors and metastases in the saline and GEM control groups were not statistically significant, nor was the 50% decrease in the LM-TT group **(Fig. 5A).** A representative example of a pancreatic KPC tumor in each group is shown in **Fig 5B.**

**Fig. 5.**
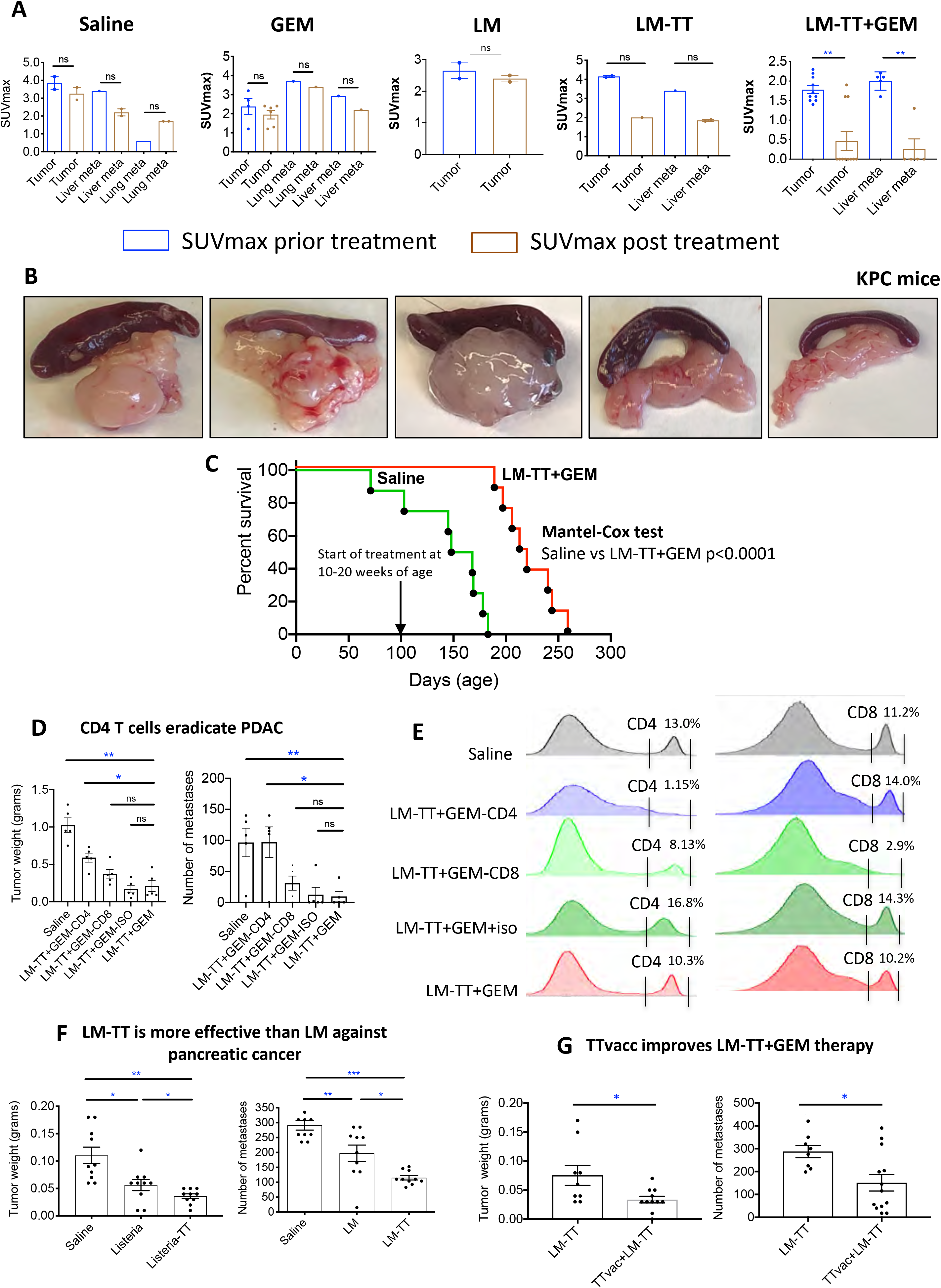
*Listeria*-TT+GEM robustly reduces advanced pancreatic cancer in KPC mice. KPC mice received the combined treatment as outlined in Fig. S1B. **(A)** Effect of treatment on advanced pancreatic cancer in KPC mice. Averages are shown for a single experiment with 5 mice in the *Listeria*-TT+GEM group, and 2 mice in each other treatment group. *Listeria*-TT+GEM was started at age 3–5.5 months, after tumors and metastases were verified through PET scan. SUVmax of tumors and metastases were measured prior (blue bar) to and post-treatment (brown bar). **(B)** Examples for each treatment group of pancreatic tumors in KPC mice**. (C)** *Listeria*-TT+GEM significantly improves survival of KPC mice. KPC mice of 10-20 weeks of age were treated with *Listeria*-TT+GEM or saline as described in Fig. S1B, but now with two instead of one high dose of *Listeria*-TT, one week apart. At the end of treatments mice were monitored until death, and survival was assessed, when defined clinical endpoints were reached. This was one experiment with 8 mice per group. Significant differences were determined by the Mantel-Cox test. **(D)** CD4 T cells eradicates pancreatic cancer *in vivo.* Mice with Panc-02 tumors and metastases were treated with anti-CD4 and anti-CD8 antibodies during *Listeria*-TT+GEM treatment. The antibodies (300μg/200μL) were administered ip every 3^rd^ day (5 doses total). Appropriate isotype antibodies were used as negative control. At the conclusion of the experiment the tumor weight and number of metastases was determined. n=7 mice per group. Mann-Whitney. *p<0.05, **p<0.01. **(E)** Depletion of CD4 and CD8 T cells was confirmed by flow cytometry of the spleen. **(F)** *Listeria*-TT is significantly more effective against advanced pancreatic cancer than LM alone. Panc-02 mice with advanced pancreatic cancer received the Listeria-TT ± TTvac as outlined in Fig 1B. **(G)** Effect of TTvac on pancreatic cancer. The results of 2 experiments were averaged with n=8-9 mice per group, and analyzed by Mann-Whitney test *p<0.05, **p<0.01, ***p<0.001. Error bars represent SEM. LM-TT = *Listeria*-TT, GEM = gemcitabine. In D, all groups were compared to the saline group. In F, all groups were compared to each other.

We also tested *Listeria*-TT+GEM therapeutically in Panc-02 mice (peritoneal model) with early and advanced pancreatic cancer. We found nearly complete elimination of early stage pancreatic cancer in mice treated with *Listeria*-TT+GEM **(Figs. S7B and E)**, and a significant reduction in primary (85%;; p<0.001) and metastatic tumor burdens (89%;; p<0.0001) in Panc-02 mice with advanced pancreatic cancer and treated with *Listeria*-TT+GEM or *Listeria*-TT alone **(Fig. S7C)** as outlined in **Fig. S1B**, compared to the saline group **(Figs. S7C and F).**

Finally, we tested *Listeria*-TT+GEM in mice with advanced orthotopically generated Panc-02 tumors and metastases. Fourteen days after injection of Panc-02 tumor cells into the pancreas, when tumors and metastases were developed (0.5-1 cm), treatment with *Listeria*-TT+GEM and control groups were started as in **Fig. S1B** and continued for two weeks. Two days after the last treatment, mice were euthanized and analyzed for tumor weight and number of metastases. A significant reduction was observed in the weight of the primary tumors (68%;; p<0.05) and in the number of liver metastases (95%;; p<0.01) of *Listeria*-TT+GEM-treated compared to the saline group **(Figs. S7D and G).**

To validate the clinical value of these efficacy studies, we analyzed the survival of KPC mice treated with *Listeria*-TT+GEM or saline. Mice were immunized with TTvacc and *Listeria*-TT+GEM as outlined in **Fig. S1B.** However, two instead of one high dose of *Listeria*-TT was administered one week apart, to deliver more TT to the TME. The *Listeria*-TT treatments were started when the mice were 10-20 weeks of age (tumors were palpable). As shown in **Fig. 5C**, KPC mice treated with *Listeria*-TT+GEM lived 2 months longer than the untreated mice, which is highly significant (Mantel-Cox p<0.0001).

### *In vivo* depletion of T cells demonstrates CD4 T cell-mediated eradication of tumors and metastases through *Listeria*-TT+GEM treatment

Since the number and the activity of CD4 T cells were significantly increased in the KPC tumors of *Listeria*-TT+GEM-treated mice in correlation with a significant reduction of tumor burden and metastases, we analyzed in further detail whether this *Listeria*-TT+GEM effect could be reduced by CD4 (or CD8 T) cells depletion *in vivo*. Antibodies were administered every third day (300 μg/dose, 5 doses total). As shown in **Fig. 5D**, the average tumor weight in mice treated with *Listeria*-TT+GEM plus CD4 antibodies was significantly increased by 64% compared to the *Listeria*-TT+GEM alone group (p=0.0159, Mann-Whitney), while the tumors in mice depleted for CD8 T cells were increased by 43% compared to the *Listeria*-TT+GEM alone group but this was not significant (p=0.2063, Mann-Whitney). As expected, the tumors in mice treated with *Listeria*-TT+GEM plus the isotype control (negative control) were not significantly different from the tumor in mice treated with *Listeria*-TT+GEM only (p>0.9999, Mann-Whitney). Similarly, the number of metastases in mice treated with *Listeria*-TT+GEM plus CD4 antibodies significantly increased by 92% compared to mice treated with *Listeria*-TT+GEM alone (p=0.0435, Mann-Whitney), while the number of metastases in mice treated with *Listeria*-TT+GEM plus antibodies to CD8 T cells was increased by 74% but this was statistically not significant (p=0.1746, Mann-Whitney) **(Fig. 5D).** Also, here the number of metastases in mice treated with *Listeria*-TT+GEM plus the isotype control was not significantly different compared to mice treated with *Listeria-TT*+GEM only (p>0.999, Mann-Whitney). Flow cytometry analysis of the spleen confirmed that CD4 and CD8 T cells were efficiently depleted *in vivo* **(Fig. 5E).** In summary, these results indicate that CD4 T cells mediated the eradication of tumors and metastases *in vivo*.

While we demonstrate above that CD4 T cells *in vivo* eradicates tumors and metastases **(Fig. 5D)**, we have not identified yet through which mechanism(s) the tumor cells were destroyed. CD4 T cells exhibit various mechanisms of tumor cell killing. This includes direct tumor cell killing through the production of perforin and granzyme B (which we think is most likely based on the production of perforin and granzyme B), or tumor cell killing through antibody-dependent cellular cytotoxicity (ADCC) (by NK cells and TT-specific antibodies generated with help of CD4 T cells), or tumor cell killing through “killer” macrophages (producing IFNγ and TNF generated with help of CD4 T cells)^48–50^. This needs to be further analyzed in detail.

### TT contributes to eradication of pancreatic cancer

An important question is to what extent TT contributes to eradication of the pancreatic cancer. This question has been analyzed through different angles. First, we compared the effect of *Listeria*-TT with *Listeria* alone on advanced pancreatic cancer in different mouse tumor models (Panc-02 and KPC). Those mice that received *Listeria*-TT also received the TT vaccine prior tumor development (**Fig. 1B**). As shown in **Fig. 5F**, the tumor weight and number of metastases was significantly lower in the mice treated with *Listeria*-TT compared to *Listeria* alone (Mann-Whitney p<0.05). A similar trend was observed in the transgenic KPC mice. *Listeria*-TT reduced the SUVmax of tumors and metastases by 50% while hardly an effect was observed by *Listeria* alone **(Fig. 5A).**

Second, we evaluated the effect of the TTvac on efficacy of the Listeria-TT treatment. For this purpose, *Listeria*-TT was administered with and without prior TT vaccinations in the Panc-02 model. As shown in **Fig. 5G**, the number of metastases and tumor weight was significantly less in the TTvac+*Listeria*-TT-treated mice compared to the mice treated with *Listeria*-TT alone (Mann-Whitney p<0.05). In summary, TT significantly contributes to the eradication of the pancreatic cancer.

### Safety of *Listeria*-TT

Considering clinical trials, the safety aspects are important. While this *Listeria* has been tested in cancer patients over 15 years (FDA approved)^27^, and appeared to be safe when injected iv, we have not tested yet if this is true for ip injections in patients. However, we have tested the safety of ip-injected *Listeria* in multiple mouse tumor models. A dose-limiting toxicity study (DLT) with *Listeria*-TT showed that the LD_50_ of *Listeria*-TT was 2×10^8^ CFU **(Fig. S7H)**. Our treatment studies used a dosage of 10^7^ CFU of *Listeria*-TT, which is well below the LD_50_.

We also analyzed potential toxicity by pathological examination, 2 days after treatment of C57Bl6 mice with *Listeria*-TT+GEM. Though a mild increase in leukocytes was observed in the spleen, liver, and lungs, no significant pathological effects of *Listeria*-TT+GEM were observed. **(Table S3).** In addition, we compared liver functions of the treated C57Bl/6 mice. A small increase in the aminotransferase (AST) level was observed in plasma from *Listeria*-TT+GEM-treated mice compared to the saline group, while alanine aminotransferase (ALT) levels were similar between the two groups **(Table S4).** Similar studies have been performed earlier in mice with breast cancer which also appeared to be safe^25^.

This is also true for GEM. In most clinical trials GEM has been administered iv. However, in some clinical trials with pancreatic and ovarian cancer GEM was injected ip. Results suggest that ip injection was even more effective than iv injection without more toxicity^51^.

## DISCUSSION

The success of cancer immunotherapies such as checkpoint inhibitors and CAR T cells in PDAC has been underwhelming, due to low immunogenicity of the tumor, strong immune suppression at multiple levels, and inefficient activation of naïve T cells in the TME^8,9, 11–14^. Moreover, low neoantigen expression levels result in poor attraction of neoantigen-specific T cells^52,53^. We addressed these problems by using TT as an alternative for neoantigens, and GEM was used to reduce immune suppression in the TME. In this study, we demonstrated the effectiveness of a tumor-targeting attenuated *Listeria*, in combination with GEM, in treating animal models of early and advanced pancreatic cancer. Our treatment strategy offers the advantages of selective delivery of immunogenic TT protein, at relatively high levels, to the TME and inside tumor cells, and altering the function of macrophages and MDSC in the TME in favor of immune stimulation. TT, now used as tumor antigen, could be an alternative for those patients that lack neoantigen expression. Moreover, we showed the contribution of TT to eradicating pancreatic cancer from different angles, i.e. *Listeria*-TT was significantly more effective than *Listeria* alone in KPC and Panc-02 models, and omitting the TT vaccine prior *Listeria*-TT treatments resulted in a significant weaker effect on metastases and tumors compared to mice that received TTvacc prior the *Listeria*-TT treatments.

We found novel effects of GEM, i.e. GEM significantly improved the migration of CD4 T cells to the KPC tumor, when added to the *Listeria*-TT treatment, which resulted in more *Listeria*-TT bacteria in the tumors and thus more TT protein in the tumors of mice treated with *Listeria*-TT+GEM compared to *Listeria*-TT alone. One potential reason for more *Listeria* in the tumors when *Listeria*-TT is combined with GEM compared to *Listeria*-TT alone is that GEM reduces immune suppression resulting in more immune selective pressure that restricts bacterial survival to the TME and inside tumor cells, to escape the immune system. Alternatively, *Listeria*-TT+GEM may have induced more necrosis (which is more hypoxic) than *Listeria*-TT alone, and Listeria survives better in hypoxic areas^37^.

Also, GEM reduced the MDSC population which down regulates L-selectin on T cells, a gene involved in T cell trafficking. L-selectin was strongly upregulated in tumors of KPC mice treated with *Listeria*-TT+GEM.

While ELISPOT data showed that both CD4 and CD8 T cells were activated by TT *in vitro* in the spleen of KPC mice treated with *Listeria*-TT+GEM, we observed predominantly CD4 T cells by IHC (CD8 T cells were present in much smaller numbers in the tumors). Also, the LNS contained both CD4 and CD8 T cells but apparently only the CD4 T cells migrated efficiently to the tumor areas. Also, in humans the TT vaccine predominantly activated CD4 T cells^54^. Moreover, *in vivo* depletion of T cells in mice treated with *Listeria*-TT+GEM clearly demonstrate that CD4 T cells contributes to the eradication of tumors and metastases *in vivo.* However, we have not determined yet through which mechanism(s) these tumor cells are destroyed by CD4 T cells. Most likely the CD4 T cells destroy tumor cells by perforin and granzyme B, but this needs to be further analyzed. In previous studies we have shown that *Listeria* also kills tumor cells through ROS^25^ and immunogenic tumor cell death^14^.

A notable finding was the appearance of well-developed LNS in close contact with the tumors in KPC mice treated with *Listeria*-TT or *Listeria*-TT+GEM. Chronic infections with bacteria lead to trafficking of T cells into the circulation and the formation of LNS in different tissues where the infection develops^46^. The formation of LNS and T cell trafficking is generated through the production of chemokines and activation of their ligands and selectins on T cells^55^. While it is unclear whether or not these may have been the source of T cells trafficking to the tumors, the Cxcr3 receptor and chemokines Cxcl9 and Cxcl10 (interacts with Cxcr3) were highly upregulated by *Listeria*-TT+GEM in KPC tumors, possibly participating in T cell trafficking to the KPC tumors. Both Cxcl9 and Cxcl10 are induced by IFNγ^56,57^, which is highly upregulated by the *Listeria*-based immunizations. Most important, perforin and granzyme B were detected in LNS of *Listeria*-TT+GEM-treated mice. Detailed studies will be required to identify the pathways of T cell migration into these tumors.

Tertiary lymph node structures (TLS) in proximity to pancreatic tumors have been reported and correlated with increased patient survival in PDAC^58^. Conversely, tumor draining lymph nodes were found to correlate with immune suppression, a more metastatic character, and poor outcome in PDAC patients^59^. These lymph nodes are different from the *Listeria*-induced LNS. Our study showed that *Listeria*-TT+GEM overcame immune suppression in the TME, as demonstrated by the RNAseq data and IHC of the KPC tumors and the LNS showing the production of perforin and granzyme, in correlation with a strong decrease in the number of metastases and tumor growth in mice. It has been suggested by others that these peritumoral LNS protect the T cells from immune suppression^60^.

*Listeria*-based immunotherapies for pancreatic cancer have also been reported by others. KPC mice in the early stage of PDAC that were immunized with *Listeria* ΔActA ΔInlB (LADD) engineered to express the Kras12GD mutation, combined with anti-CD25 Abs and cyclophosphamide, demonstrated delayed PanIN progression and moderately but significantly improved survival of KPC mice when treatments were started at 6-8 weeks of age^61^. However, no improvement of survival was observed when the treatments were started at 8-12 weeks of age. In contrast, *Listeria*-TT+GEM improved the survival of KPC mice with 2 months, when treatments were started at 10-20 weeks of age (Mantel-Cox test p<0.0001), underlining the significance of our results. In another interesting study using the flank model, 4T1 and CT26 mice were immunized with LADD expressing AH1, also expressed by the tumors. This stimulated migration of tumor-specific CD8 T cells into tumors, reducing Tregs and converting M2 macrophages to M1^62^. In addition, mouse models of cervical, breast, and skin cancer have been successfully treated with *Listeria* expressing E7, Her2/neu, Flk-1, CD105, Mage-b, ISG15, and HMW-MAA^63^.

*Listeria* has also been tested in clinical trials in patients with various cancers, including pancreatic cancer, showing safety and tumor-specific T cell responses, however only modestly improved survival (for a review see Forbes et al)^29^. As mentioned earlier, the concept of *Listeria* previously tested in clinical trials is completely different from our concept. The *Listeria* used in the clinical trials is based on a “classical concept”, i.e. the delivery of tumor-specific antigens into dendritic cells to stimulate naïve and memory T cells to these antigens which are naturally expressed by the tumors, while our concept is based on colonizing and changing the TME by *Listeria*, and delivery of a highly immunogenic childhood vaccine antigen TT into tumor cells by Listeria, and reactivating pre-existing memory T cells to TT. This “classical concept” in clinical trials is partly the result of altering the *Listeria* (deletion of *ActA* and *InlB*) to improve safety and consequently removing its ability to spread and/or multiply *in vivo*, and partly the result of administering *Listeria* intravenously (iv). We administer the *Listeria* in our mouse models intraperitoneally (ip). As demonstrated in this study, *Listeria* administered iv hardly reach tumors and metastases (pancreatic and breast cancer), but when administered ip *Listeria* colonizes the TME abundantly, delivers TT to the TME and inside tumor cells attracting CD4 T cells, and infects/alters MDSC and TAM. In multiple biodistribution studies in mice with pancreatic and breast cancer we found that *Listeria* was rapidly eliminated most likely by the immune system in blood^10,22,26^, because the blood lacks immune suppression, while the TME is heavily immune suppressed. Of note is that MDSC in the areas of tumors are more immune suppressive than in the blood circulation (personal communication Dmitry Gabrilovich, AstraZeneca, Gaithersburg, MD). Thus, when *Listeria* bacteria are injected ip they directly infect the immune suppressive MDSC, which then migrate to the TME resulting in spread of the *Listeria* through the TME. As mentioned earlier, we have shown in various mouse tumor models that ip injection of *Listeria* is completely safe but this needs to be studied in patients with PDAC.

Last but not least, elderly cancer patients react less efficiently to vaccines than young adults^64,65^, which is also true for cancer immunotherapy^41,66^, mainly due to lack of naïve T cells later in life^40^. We believe that our approach of reactivating memory T cells generated during childhood overcomes the need for naïve T cells at older age, potentially making this therapeutic strategy more effective for older patients. This may lead to new treatment modalities against pancreatic cancer where other types of therapies fail.

## MATERIALS AND METHODS

### Animal care

C57BL/6 male and female mice aged 3 months were obtained from Charles River. KPC mice^31^ were generated in the laboratories of Drs. Gravekamp and Libutti, as described previously^26^. C57Bl/6N-Tg(Cfms-gal4-vp16)-(UAS-eCFP)^36^ were maintained in the laboratory of Dr. Condeelis. All mice were housed in the animal husbandry facility at Albert Einstein College of Medicine according to the Association and Accreditation of Laboratory Animal Care guidelines, and kept under BSL-2 conditions as required for *Listeria* treatments. All KPC mice were genotyped by Transnetyx (Cordova, TN).

### Cell lines

The Panc-02 cell line was derived from a methylcholanthrene-induced ductal adenocarcinoma growing in a C57BL/6 female mouse^67^. Panc-02 cells were cultured in McCoy’s5A medium supplemented with 10% fetal bovine serum, 2 mM glutamine, non-essential amino acids, 1 mM sodium pyruvate, and 100 U/ml Pen/Strep. The Panc-02 cell line expressing Dendra-2 was developed in the laboratory of Dr. Gravekamp. Briefly, Panc-02 cells were transfected with pDendra2C (ClonTech), and positive cells were selected using neomycin and FACS sorting. The KPC tumor cell line (kindly provided by Dr. Vinod Balachandran, MSKCC, NY), was derived from a pancreatic tumor of a transgenic KPC mouse (*Kras^G12D/+;;^LSL-Trp53^R172H/+^*)^68^. The 4T1 cell line was derived from a spontaneous mammary carcinoma in a BALB/c mouse^69^. Various 4T1 sub lines have been generated with different patterns of metastases^70^, which is kindly provided to our lab by Dr. Fred Miller, Karmanos Institute, Wayne State University, Michigan, MI). The 4T1 cell line used in this study is highly aggressive, metastasizing predominantly to the mesenteric lymph nodes (MLN), and less frequently to the diaphragm, portal liver, spleen and kidneys^71^.

### Listeria and Listeria-TT

In this study, an attenuated *Listeria monocytogenes* (*Listeria*) was used as the vehicle for delivery of TT_856-1313_^72^ to the TME and inside tumor cells. The *Listeria* plasmid pGG-34, expressing the non-cytolytic truncated form of Listeriolysin O (LLO) under control of the hemolysin promoter *(Phly)* and the positive regulatory factor A (*prfA*), with mutations to further reduce pathogenicity, have been described elsewhere^33^. *Listeria*-TT_856-1313_ was developed in our laboratory **(Fig. 1A)**. Briefly, TT_856-1313_^72^ was cloned by PCR into pCR2.1 using primers containing *XHo*I and *XMa*I restriction sites and a myc Tag for detection of TT. (**PrF5’:** CTC GAG TCA ACA CCA ATT CCA TTT;; **PrR5’:** CCC GGG TTA TAG ATC TTC TTC TGA AAT TAG TTT TTG TTC TGT CCA TCC TTC ATC TGT) Subsequently the pCR2.1-*XHo*I-TT_856-1313_-Myc-*Xma*I and pGG34 plasmids were digested with *XHo*I and *XMa*I, and the *XHo*I-TT_856-1313_-myc-XMaI DNA fragment was cloned into the pGG34 vector, and then electroporated into the *Listeria* background strain *XFL7*^33^. The *Listeria*-TT_856-1313_ was characterized by DNA sequencing and by evaluating the secretion of TT_856-1313_ into the culture medium **(Fig. 1A).** The TT_856-1313_ fragment contains both mouse and human immunodominant T cell epitopes^54,73^.

### Infection of tumor cells *in vitro*

The *in vitro* infectivity of Panc-02 tumor cells was assessed as described previously^25^. Briefly, 10^6^ cells/ml were infected with 10^7^ CFU of *Listeria* or *Listeria* -TT for 1 hr at 37°C in culture medium, then incubated with gentamicin (50 μg/ml) for 1 hr to kill extracellular *Listeria*. Finally, cells were washed with phosphate-buffered saline (PBS), lysed in water, and serial dilutions were plated onto LB agar to quantify the infection rate the next day.

### Evaluation of cell death

As described previously^25^, tumor cell killing by *Listeria* or *Listeria*-TT was determined *in vitro* as follows. Panc-02 cells (3×10^3^) plated in 96 well plates were infected with 10^7^ CFU/well of *Listeria* or *Listeria*-TT for 3 hrs at 37°C, then gentamicin (50 μg/ml) was added. Live and dead cells were counted the next day using Trypan blue staining. A similar experiment was performed with human Mia-PaCa2 pancreatic tumor cells as outlined in the text.

### Biodistribution of *Listeria*-TT

C57Bl/6 mice were injected with 10^6^ Panc-02 mouse pancreatic tumor cells as described above, and 14 days later injected with a single large dose 10^7^ CFU of *Listeria*-TT. Mice were euthanized at various time points after the injection, and metastases, tumors and normal tissues were dissected, weighed, and analyzed for CFU of *Listeria*-TT, as described previously^26^. In a separate study, Panc-02 tumors were generated, then treated with 12 high doses of *Listeria* (10^7^ CFU), and two days after the last treatment analyzed for CFU of *Listeria* in tumor and normal tissues.

Nude mice were orthotopically injected with 10^6^ Mia-Paca2 human pancreatic tumor cells in the pancreas, and 6 weeks later with a single large dose of 5×10^7^ CFU of *Listeria*. Mice were euthanized at various time points after the injection, and metastases, tumors and normal tissues were dissected, weighed, and analyzed for CFU of *Listeria*, as described previously^26^.

### Dose-limiting toxicity (DLT)

C57Bl6 mice were injected ip with various doses of *Listeria*-TT, and survival was monitored over the next 20 days, as described previously^26^.

### Tumor development

Panc-02 tumor cells were injected into the mammary fat pad (peritoneal cavity model) (10^5^) or orthotopically into the pancreas (10^6^) of C57Bl/6 mice. When injected into the mammary fad pat, a relatively small primary tumor develops at the place of injection (peritoneal membrane) and tumor cells metastasize via the peritoneal cavity to other organs, predominantly to the pancreas and liver, and less abundantly to mesenchymal lymph nodes along the gastrointestinal tract and the diaphragm^22^, but when injected into the pancreas tumor cells a primary tumor develops in the pancreas and tumor cells metastasize to the liver only^74^. In KPC mice, multiple primary tumors develop spontaneously in the pancreas, metastasizing to the liver and lungs^26,31^.

### Orthotopic Panc-02 or KPC model

Orthotopic Panc-02 or KPC tumors were generated in C57Bl/6 mice as described previously^75^. Briefly, mice were anesthetized with Ketamine/Xylazine (respectively 100 mg and 10 mg/Kg, ip), the hair was removed at the location of the spleen, and the skin was sterilized with betadine followed by 70% alcohol. The animal was covered with gauze sponge surrounding the incision site. A 1 cm incision was made in the abdominal skin and muscle just lateral to the midline and directly above the spleen/pancreas to allow visualization. The spleen/pancreas was gently retracted and positioned to allow injection of 10^6^ Panc-02 or KPC tumor cells directly into the pancreas, from the tail all the way to the head of the pancreas. To prevent leakage of injected cell suspension, the injection site was tied off after tumor cell injections with dissolvable suture. The spleen/pancreas were then replaced within the abdominal cavity, and both muscle and skin layers closed with sutures. Following recovery from surgery, mice were monitored and weighed daily. A palpable tumor appeared within 2 weeks in the pancreas, and metastases in the liver began to develop within 2-4 weeks.

### Protocol for treatment of Panc-02 and KPC mice with *Listeria*-TT+GEM

A detailed rationale for this treatment protocol and schematic view of all treatments are shown in **Fig. S1B.** <u>Panc-02 mice.</u> To generate memory T cells to TT, C57Bl/6 mice were immunized twice intramuscularly with the Tetanus vaccine (the same used for childhood vaccinations, 0.3 μg/50 μL) 1 week apart. Subsequently, Panc-02 tumor cells (10^5^/100 μL) were injected into mammary fat pad. A single high dose of *Listeria*-TT (10^7^ CFU) was injected ip, either 3-5 days after tumor cell injection when tumors were 1-3 mm (early pancreatic cancer) or 10-14 days after tumor cell injection when tumors were 5-10 mm (advanced pancreatic cancer). In addition, Panc-02 tumor cells (10^6^/50 μL) were injected into pancreas, and 10-14 days later when tumors were 5-10 mm, a single high dose *Listeria*-TT (10^7^ CFU) was injected ip. Three days later, GEM (Gemizan: Zuventus, Mombay, India) treatment was started (1.2 mg/mouse, every 3 days) and continued for 14 days (six doses in total). Concomitantly, low doses of *Listeria*-TT were administered daily for 2 weeks (14 doses in total). All mice were euthanized 2 days after the last treatment, and visually analyzed for tumor weight and the number of metastases, as described previously^22^. <u>KPC mice</u>. KPC mice, age 3.5-5 months, received the same treatment described above, after verification of the presence of tumors and metastases by PET scan.

### Survival

KPC mice were treated with TTvacc and *Listeria*-TT+GEM as outlined in **Fig. S1B.** At the end of treatments mice were monitored without any further treatment until they succumbed spontaneously, or were terminated upon appearance of severe pre-morbid symptoms requiring euthanasia as specified by our approved animal use protocol.

### *In vivo* depletion of T cells

CD4 and CD8 T cells were depleted in C57Bl/6 mice with orthotopic Panc-02 tumors during *Listeria*-TT+GEM treatment. Briefly, 10-14 days after tumor cell injection (when tumors were approximately 5mm^2^), mice were treated with *Listeria*-TT+GEM and with 300 μg of anti-CD4 (Clone GK1.5;; Cat#BE0003-1: BioXCell) or anti-CD8 (Clone YTS169.4;; Cat#BE0117: BioXCell) antibodies (five injections every third day). All mice were euthanized 2 days after the last anti-CD4 or -CD8 treatment and analyzed for tumor weight and number of metastases. As control, isotype-matched rat antibodies against HRPN were used (Clone LTF-2;; Cat#BE0090: BioXCell).

### Intravital multiphoton imaging (IVMI)

Panc-02 tumor cells (10^6^) expressing Dendra-2 were injected into the pancreas of transgenic mice [C57Bl/6N-Tg(Cfms-gal4-vp16)-(UAS-eCFP)]^36^ in which the macrophages were labeled with cyan fluorescent protein (CFP). Three weeks later, when a pancreatic tumor developed and on the same day as imaging, *Listeria*-TT was labeled *ex vivo* with Alexa-680 by incubation with rabbit anti-*Listeria* polyclonal antiserum (dilution 1:200) and anti-rabbit IgG-Alexa-680 (dilution 1:250). *Listeria*-TT-Alexa-680 was injected ip, and four hours later, when *Listeria*-TT had accumulated in the TME and infected tumor cells, mice were anesthetized using isoflurane. Pancreatic tumors were externalized through a small incision (∼0.7 cm) through the skin and peritoneal wall, and stabilized for imaging using previously published protocols^76^. Multiphoton intravital microscopy was performed using a custom-built two-laser multiphoton microscope^77^. Tumors were imaged with a femtosecond laser set to 880 nm for excitation of CFP or the green form of Dendra-2, and an optical parametric oscillator set to 1240 nm for excitation of Alexa 680. Z-stacks to a depth of 45 µm were acquired with a 1 µm slice interval. During the course of imaging, the animal was placed in a heated chamber maintained at physiological temperature. Mice were monitored for heart rate, breathing rate, pulse distension, breath distension and oxygen saturation using a pulse oximeter (MouseSTAT, Kent Scientific).

### PETscan

Positron emission tomography (PET) was performed as described previously^78^. Briefly, the effect of the *Listeria*-TT+GEM treatment on the tumor and metastases was monitored by microPET (Siemens Imaging System) prior to and post-treatment. For this purpose, mice were injected with the positron emitter [^18^F]-labeled deoxyglucose. The uptake of ^18^F-FDG by tumors and metastases was quantified by microPET as described previously^22^. Increased FDG uptake reflects increased metabolic activity of cancer cells, expressed as SUVmax.

### Flow cytometry

Immune cells from spleens, blood, or metastases of mice were isolated as described previously^66^. Anti-CD45 antibodies was used to identify the leukocyte population in tumors and metastases. Anti-CD3 and anti-CD8 antibodies were used to identify CD8 T cells, anti-CD3 and anti-CD4 to identify CD4 T cells, anti-CD11b and anti-Gr1 to identify MDSC, and anti-CD11b and anti-F4/80 to identify TAM. To detect intracellular cytokine production, the cells were incubated with GolgiPlug (1 μg/ml) for 6 hrs, then treated with Cytofix/Cytoperm according to the manufacturer’s instructions, before staining with antibodies to intracellular cytokines, IFNγ, granzyme B, perforin, IL-6, IL-10, TNFα, and other markers such as the T cell response inhibitor MARCO, the T cell activation marker CD69, or the costimulator CD80. Appropriate isotype controls were included for each sample. 50,000-100,000 cells were acquired by scanning using a special order LSR-II Fluorescence Activated Cell Sorter system (Beckton and Dickinson), and analyzed using FlowJo 7.6 software. Cell debris and dead cells were excluded from the analysis based on scatter signals and use of Fixable Blue or Green Live/Dead Cell Stain Kit.

### ELISPOT

Spleen cells were isolated from treated and control Panc-02 and KPC mice for ELISPOT (BD Biosciences, cat# 551083) analysis, as described previously^71^. To detect T cell responses to TT, 10^5^ spleen cells were incubated with purified TT protein (5 μg/mL). The frequency of IFNγ-producing spleen cells was measured 72 hrs later using an ELISPOT reader (CTL Immunospot S4 analyzer). To determine the frequency of IFNγ-producing CD4 and CD8 T cells, spleen cells were depleted for CD4 and CD8 T cells using magnetic bead depletion techniques according to the manufacturer’s instructions. A similar protocol was used for T cells isolated from Panc-02 orthotopic tumors.

In addition, T cell responses to TAA *Survivin* was analyzed by ELISPOT as described previously^14^. Briefly, spleen cells were isolated from *Listeria*-TT+GEM-treated and control KPC mice for ELISPOT. To detect T cell responses to *Survivin*, 2×10^5^ spleen cells were incubated with 50 μg/mL of *Survivin* protein (MGAPALPQIWQLYLKNYRIATF KNWPFLEDCACTPERMAEAGFIHCPTENEPDLAQCFFCFKELEGWEPDDNPIEEHRKHSPGCAFLTVKKQMEELTVSEFLKLDRQRAKNKIAKETNNKQKEFEETAKTTRQSIEQLA A)(LSBio, Seattle, WA: Cat no LS-G22824/147580) for 72 hrs. After the 72hr incubation, the frequency of IFNγ-producing spleen cells was measured using an ELISPOT reader as described above. To determine the frequency of IFNγ-producing CD4 and CD8 T cells, spleen cells were depleted for CD4 and CD8 T cells using magnetic bead depletion techniques as described above.

### Western blotting

Expression of TT protein in tumor cells was analyzed by western blotting as described previously^79^. Briefly, cells were lysed in RIPA buffer containing protease inhibitors, and proteins were separated on 4-12% gradient SDS-polyacrylamide gels, then electro-transferred to PVDF membrane. Membranes were incubated with rabbit anti-myc IgG, followed by horseradish peroxidase-conjugated goat anti-rabbit IgG. Detection was with a chemiluminescence detection kit. Antibody to β-actin was used to ensure equal loading.

### Immunohistochemistry (IHC) and Immunofluorescence

Tumors were dissected from pancreas and immediately fixed with buffered formalin, and the tissue was embedded in paraffin. Sections (5 µm) were sliced and placed on slides, then deparaffinized at 60°C for 1 hr, followed by xylene, an ethanol gradient (100-70%), water, and PBS. Slides were then incubated for 30 min in 3% hydrogen peroxide followed by boiling in citrate buffer for 20 min. Once the slides were cooled, washed, and blocked with 5% goat serum, the sections were incubated with primary antibodies such as anti-CD4 (1:100 dilution), anti-CD8α (1:400 dilution), anti-Perforin (1:300 dilution), anti-Granzyme (1:200 dilution), anti-CD31 (1:100 dilution), anti-TT (1:50 dilution), or anti-MHC class II (Invitrogen cat#14-5321-82)(1:100 dilution), followed by incubation with secondary antibody (mouse anti-goat IgG-HRP), and SignalStain® Boost IHC Detection Reagent (Cell Signaling Technology, cat# 8114S). Subsequently, the slides were incubated with 3,3’-diaminobenzidine (DAB)(Vector Laboratories, cat# SK-4100), counterstained with hematoxylin, dehydrated through an ethanol gradient (70-100%) and xylene, and mounted with Permount. The slides were scanned with a 3D Histech P250 High Capacity Slide Scanner to acquire images and quantification data. Secondary antibodies without primary antibodies were used as negative control.

KPC tumors were digested using Gentlemacs and Collagenase and Dispase as described above, and tumor cells were isolated by magnetic beads (CD45-negative fraction). The CD45-negative fraction was stained with anti-MHC class II-FITC antibodies (Invitrogen Cat#11-5321-82)(dilution 1:100), and analyzed by a ZOE Fluorescent Cell Imager (Biorad).

### RNAseq

#### Sample preparation for RNA seq

Total RNA was isolated from the metastases and tumor tissues, treated with DNase, and evaluated using the Agilent Bioanalyzer. The purified high quality total RNA was submitted to the Epigenetics Shared Facility (ESF) for assay and analysis. An RNAseq assay using 25 ng of RNA was performed and validated by the ESF.

#### Whole-transcriptome library preparation and high-throughput RNA sequencing

ESF personnel performed rRNA-depletion using a Ribo-Zero rRNA Removal Kit, and reverse transcription using random hexamers and oligo(dT) to synthesize ds-cDNA. Libraries were prepared with adapters and barcodes for multiplex sequencing, and sequenced using the Illumina HiSeq 2500. Raw FASTQ files were trimmed for adapter sequences using quart. FASTQ file sequence quality was evaluated using FastQC^80^, and read coverage determined using the RSeQC package^81^.

#### RNAseq data analysis

We generated a total of 150 RNAseq runs. For each run we obtained at least 20 million reads, with >70% of them aligned to the mouse genome. We used the Bowtie/Tophat/Cufflinks/Cuffdiff software suite^82–84^, applying the Tuxedo protocol. The RNAseq analysis was performed on tumors from transgenic KPC mice treated with *Listeria*-TT+GEM or saline (control group), and on orthotopic KPC tumors of mice treated with *Listeria*-TT+GEM, *Listeria*-TT, GEM or saline.

### RNAscope

RNA *in situ* hybridization (RNAscope, ACD Bio-Techne, Minneapolis, MN, USA) was performed on formalin fixed paraffin-embedded 5µm sections using the RNAScope® Multiplex Fluorescent V2 Assay according to manufacturer’s instructions. Probes used were directed against mouse CD4 (cat # 406841), mouse CD8 (cat # 401681-C2), and anti-mouse IFNγ (cat # 311391-C3). The Fluorophore Opal™ 520 (Akoya Biosciences, Cat # FP1494001KT) is C1, Opal™570 (Akoya Biosciences, Cat # FP1488001KT) C2 and Opal™ 690 C3 (Akoya Biosciences, Cat # FP1497001KT) were used for detection and DAPI was used as counter stain and ProLong Gold Antifade Mountant (Thermo Fisher Scientific, cat # P36930) was used as a mounting media. The slides were then imaged at Leica SP5 Confocal Microscope. Images were acquired using a 40x Oil objective (NA=1.25) with a 4x zoom.

### Statistics

To statistically compare the effects of *Listeria*-TT+GEM on the growth of metastases and tumors or on immune responses in the mouse models, the Mann-Whitney was applied using Prism. *p<0.05, **<0.01, ***<0.001, ****<0.0001. Values of p<0.05 were considered statistically significant. To statistically compare the effects of *Listeria*-TT+GEM in a survival study a Kaplan-Meier curve was generated using Mantel-Cox. ****p<0.0001.

## Acknowledgements

We greatly thank Ms. Hong Zhang, Department of Pathology, Einstein for providing outstanding training and support regarding the IHC and image analysis.

## Funding

This work was supported by the Pancreatic Cancer Action Network (PCAN) 422247, a private donation of Janet and Marty Spatz, another private donation Steve Bigelsen, NCI Administrative Supplement 3P30CA013330-44S3, and NCI cancer center support P30CA013330 (Flow Cytometry Core, Pathology Core, MicroPET Core, Genetics Core, Imaging Core).

## Authors contribution

Conception, study design, and study supervision;; development of methodology;; administrative, technical, or material support (i.e., reporting or organizing data, constructing data bases);; writing, review, and/or revision of the manuscript: B.C. Selvanesan, D. Chandra, W. Quispe-Tintaya, Z. Yuan, S.K. Libutti, C. Gravekamp. Acquisition of data (provided animals, cell lines, developed *Listeria*-TT construct, acquired immunological and efficacy data, provided facilities, etc.): B.C. Selvanesan, D. Chandra, W. Quispe-Tintaya, A. Jahangir, A. Patel, K. Meena, R. Alves Da Silva, J. Li, S. Siddiqui, DeMarais V, Friedman M, A. Beck, L. Tesfa, W. Koba, Z. Yuan, S.K. Libutti, Balachandran V, C. Gravekamp. Analysis and interpretation of data (statistical analysis, biostatistics, computational analysis): B.C. Selvanesan, D. Chandra, W. Quispe-Tintaya, A. Jahangir, A. Patel, K. Meena, R. Alves Da Silva, A. Beck, L. Tesfa, W. Koba, J. Chuy, Z. Yuan, S.K. Libutti, X. Zhang, C. Gravekamp. **Competing interests:** Dr. Gravekamp has ownership interest in a patent application filed for the *Listeria*-recall antigen concept (96700/2230). No other competing interests were disclosed by the authors.

## Data and materials availability

All data associated with this study are available in the paper or the Supplementary Materials.

## SUPPLEMENTAL MATERIALS

**Fig S1A:** *Listeria*-Recall Antigen Model

**Fig S1B:** Treatment protocol *Listeria*-TT+GEM

**Fig S2:** *Listeria* in KPC tumors

**Fig S3ABC:** Intraperitoneally but not intravenously injected *Listeria* colonizes tumors and metastases, and reactivation of pre-existing memory T cells to TT in Panc-02 mice

**Fig S4AB:** Gating of CD4 and CD8 T cells in spleens of tumor-bearing Panc-02 mice treated with *Listeria*-TT and GEM, in combination or separately

**Fig S5A-H:** Detail of CD4 and CD8 T cells in tumors of *Listeria*-TT+GEM-treated and control mice IHC

**Fig S6ABC:** IHC of KPC tumors stained with antibodies to TT, CD4, and CD31

**Fig S6D:** Detail of lymph node-like structures associated with pancreatic tumors in mice

**Fig 6E:** Perf/GranzB in LNS

**Fig S7A-H:** RNAseq KPC tumors (genes involved in T cell migration), treatment of early and late pancreatic cancer, DLT

**Fig S8:** MHC class II expression in KPC tumors

**Fig S9:** RNAscope

**Fig S10:** RNAseq on tumors including LNS of Saline and *Listeria*-TT+GEM treated KPC mice only

**Fig S11:** Gating of MDSC and TAM in metastases of tumor-bearing Panc-02 mice

**Table S1:** Number of CFU of *Listeria* in mouse Panc-02 tumors and normal tissues of C57Bl/6 mice

**Table S2:** Number of CFU of *Listeria* in MiaPaCa2 tumors and normal tissues of nude mice

**Table S3:** Pathological examination of tissues in C57Bl/6 mice 2 days after the last treatment with *Listeria*-TT and Gemcitabine

**Table S4:** The effect of *Listeria*-TT and Gemcitabine on liver functions in C57Bl/6 mice Additional Supplementary Information

## SUPPLEMENTARY INFORMATION

**Fig. S1.**
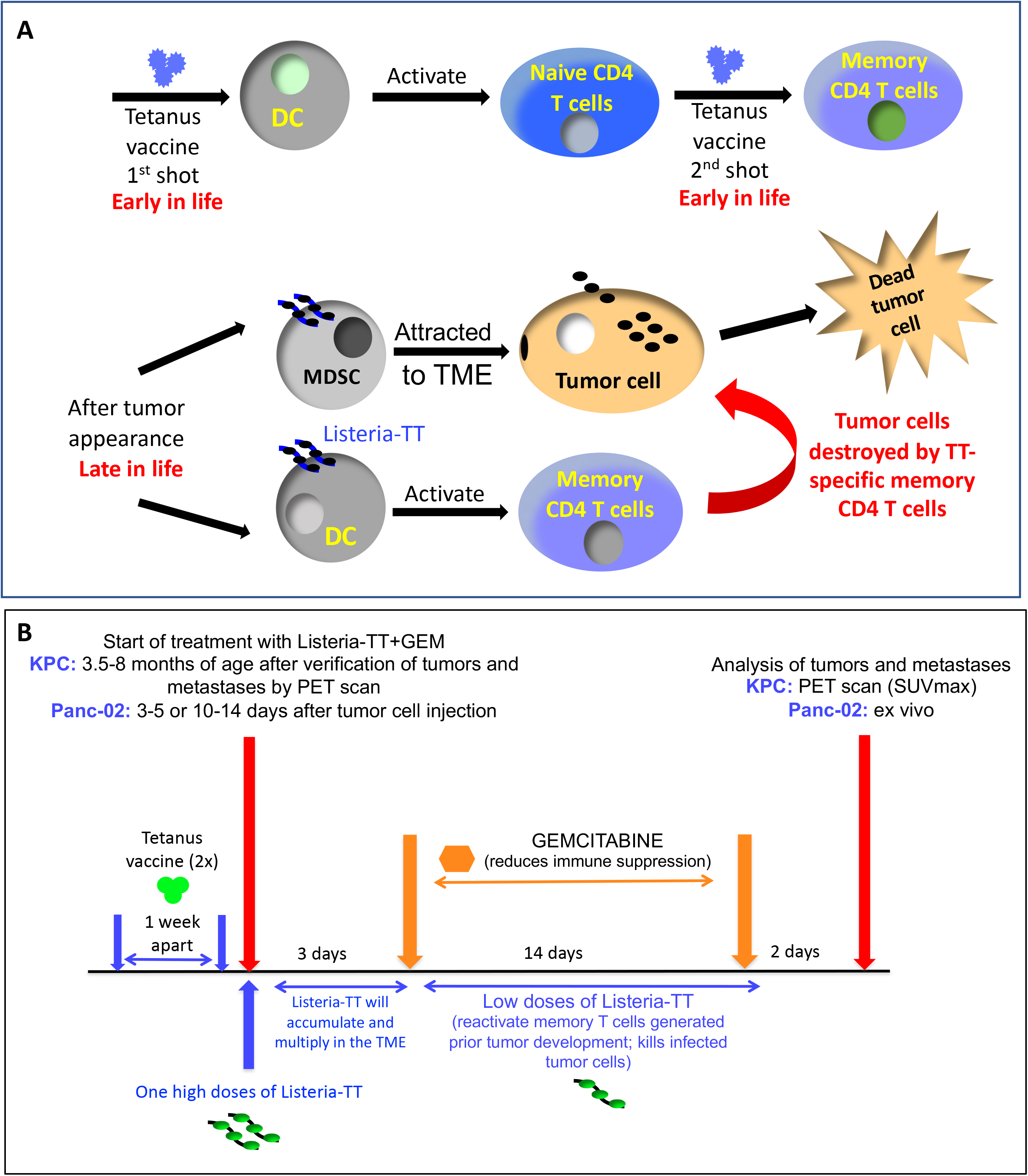
**(A) Mechanistic view of the *Listeria*-recall antigen concept.** Tetanus vaccine will be used to generate memory T cells to recall antigen TT in mice prior to tumor development. Antigen-presenting cells (DC) take up and present TT protein to naïve T cells. Upon repeated exposures to TT, naïve T cells differentiate into memory T cells, which circulating in the blood for life. These memory cells are to be reactivated after appearance of the primary tumor and metastases, as follows. To deliver TT into tumor cells, mice receive a high dose of *Listeria*-TT bacteria, whose replication will lead to more TT protein in the tumor cells. Subsequently, frequent low dose immunizations with *Listeria*-TT reactivate TT-specific memory T cells. Finally, TT-specific memory T cells migrate to the tumor and kill tumor cells presenting TT epitopes. Since attenuated *Listeria* bacteria colonize the TME, they will also infect MDSC and macrophages (not depicted). **(B) Protocol for treatment of Panc-02 and KPC mice with *Listeria*-TT+GEM.** In the Panc-02 model, mice were immunized twice with Tetanus vaccine (TTvac) to generate memory T cells. Subsequently, Panc-02 tumor cells were injected and tumors allowed to develop to the early stage of pancreatic cancer (1-3 mm) or advanced stage (5-10 mm). After an initial high ip dose of *Listeria*-TT (10^7^ CFU), it accumulated and multiplied in the TME for 3 days. Low doses of GEM (1.2 mg/dose) were then given every 3 days for 14 days in order to eliminate MDSC and TAM. At this point, MDSC are no longer required to bring *Listeria*-TT to the TME. Instead, reducing MDSC and TAM populations by GEM serves to improve T cell responses. Concomitantly, daily low doses of *Listeria*-TT (10^4^ CFU) reactivate TT-specific memory T cells, improved by GEM. Mice were euthanized two days after the last treatment. Panc-02 mice were analyzed for tumor weight and the number of metastases. In the KPC model, the presence of tumors and/or metastases were first verified by PET scan before mice received the same treatments described above. The metabolic activities (SUVmax) of tumors and metastases in KPC mice were measured by PET scan. Comparisons of SUVmax pre- and post-treatment provide a measure of the growth of tumors and metastases. In the survival study, a similar treatment protocol as in the efficacy study was used with one exception: two instead of one high dose of Listeria-TT (10^7^ CFU) was administered one week apart, to deliver more TT in the tumors and metastases. LM-TT = *Listeria*-TT.

**Fig. S2:**
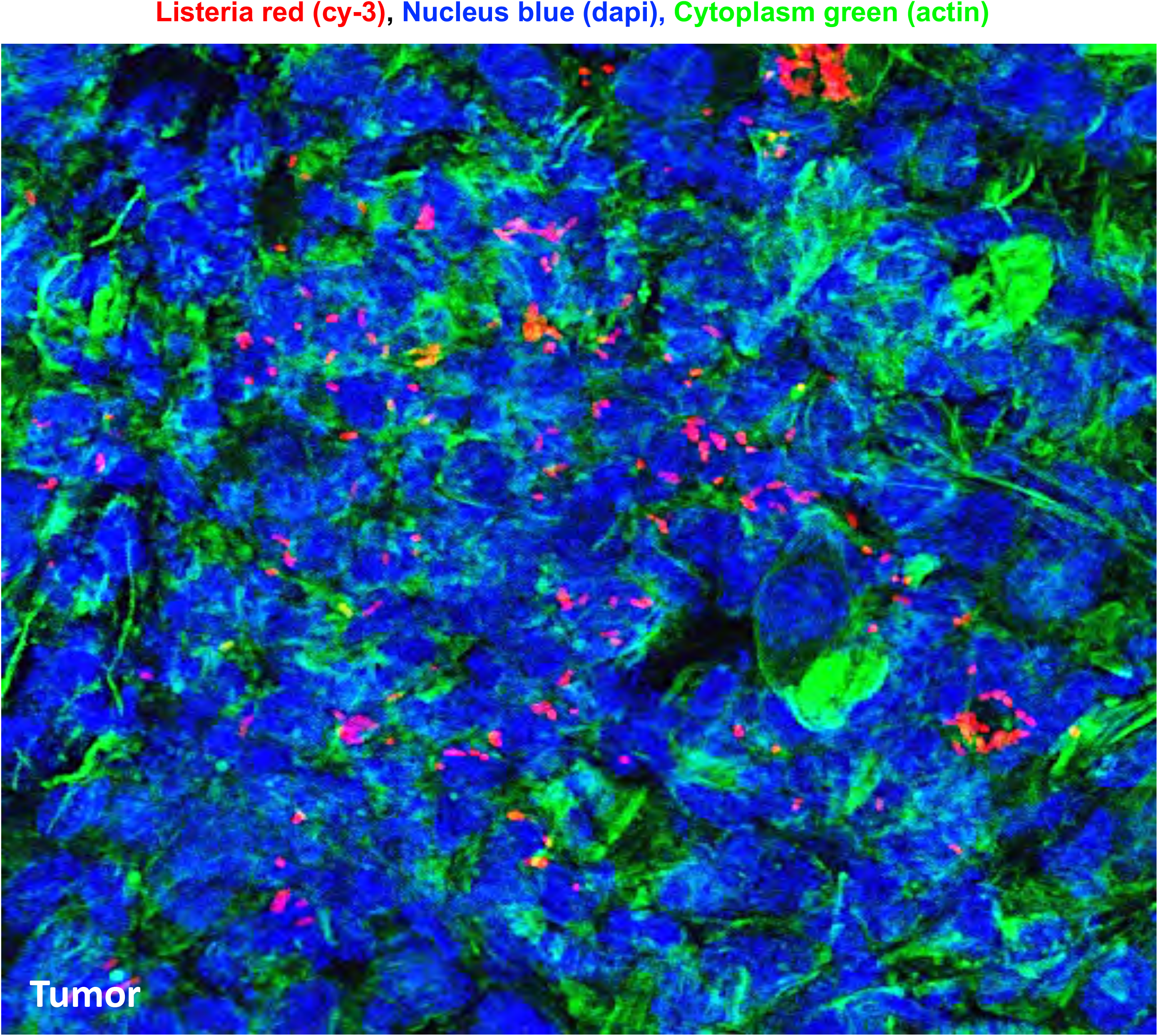
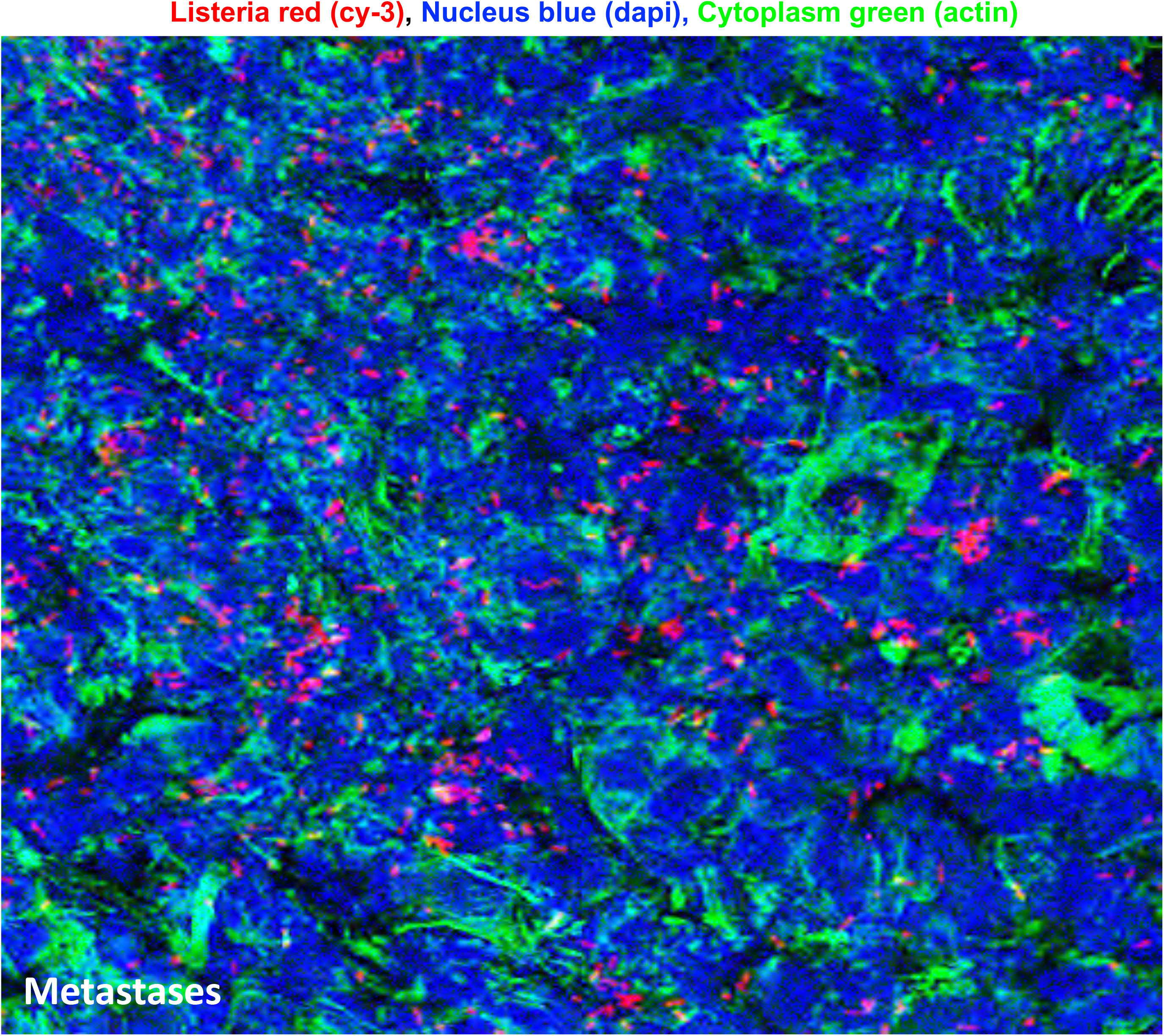
*Listeria* in pancreatic tumor. The presence of *Listeria* in pancreatic tumors **(A)** and metastases **(B)** was confirmed by confocal microscopy of Panc-02 mice. *Listeria* are red (Cy-3), nuclei blue (dapi), and cytoplasm green (actin staining).

**Fig. S3:**
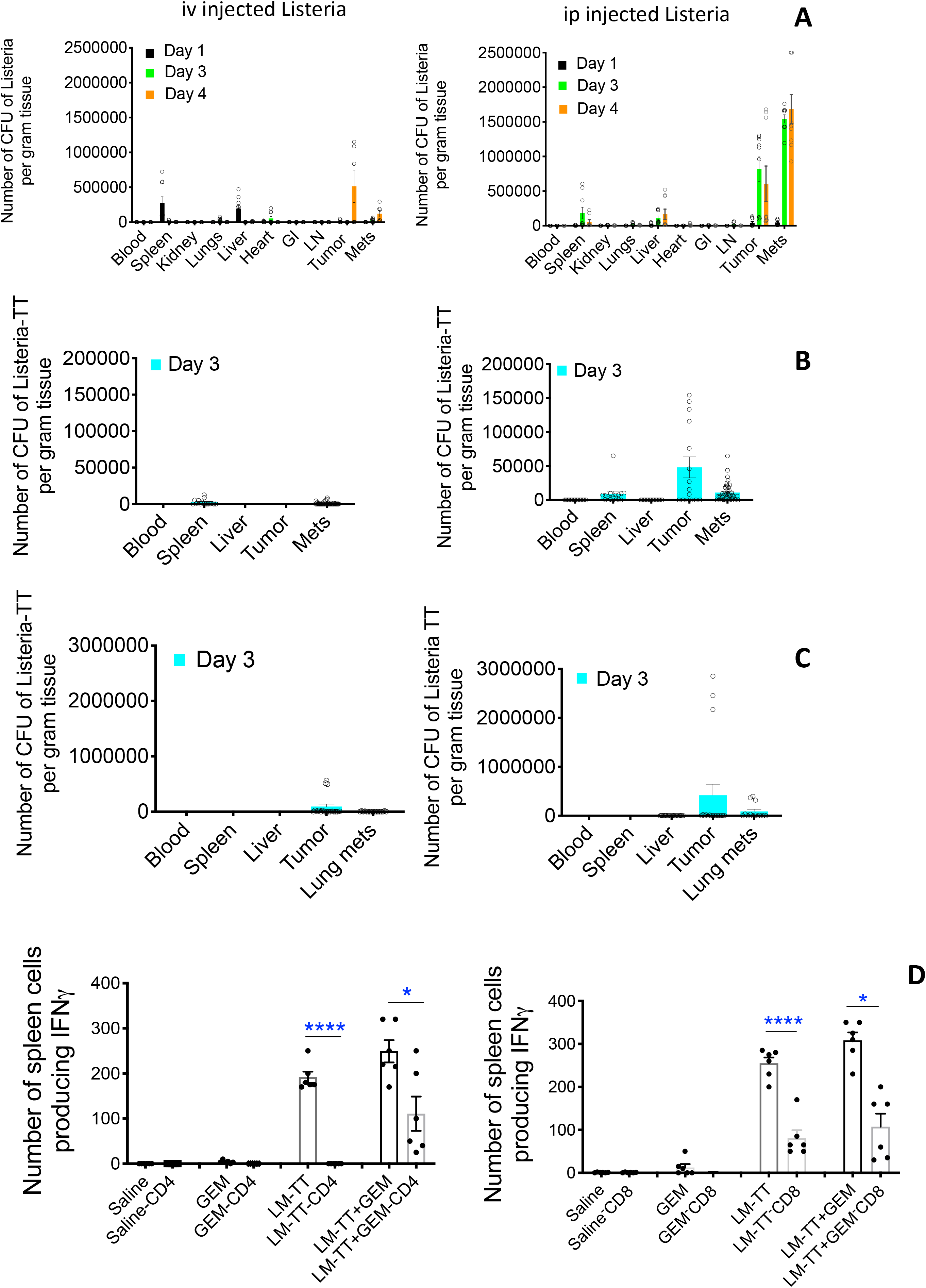
Intraperitoneally but not intravenously injected *Listeria* colonizes tumors and metastases. Experiments conducted in three different mouse models showed that *Listeria* colonized the TME efficiently if injected intraperitoneally (ip), but very poorly or not at all when injected intravenously (iv). **(A)** Balb/C mice were injected into the mammary fat pad with 4T1 tumor cells, resulting in primary tumors in the peritoneal membrane and multiple metastases in the liver, pancreas and along the GI, as described previously^1^. After tumors and metastases had developed, *Listeria* (10^7^ CFU) was injected ip or iv, and the number of *Listeria* bacteria was determined 1, 3, or 4 days later in blood and all organs, including tumors and metastases, as done previously^2^. **(B)** A similar experiment in the Panc-02 mouse peritoneal cavity model. Panc-02 tumor cells were injected into the mammary fat pad of Panc-02 mice, resulting in a primary tumor in the peritoneal membrane and multiple metastases in the pancreas, liver and gastrointestinal tract, as described previously^3^. Ten days later, when primary tumor was palpable at the place of injection and metastases had developed, *Listeria*-TT was administered as above. CFU in the blood, tumors, metastases and normal tissue (spleen) were determined 3 days later as described previously^3^. **(C)** A similar experiment in the 4T1 model. Injection of 4T1 tumor cells into the mammary fat pad resulted in a primary mammary tumor and metastases in the lungs within 4 weeks, *Listeria*-TT was administered, and CFU measured 3 days later, as described previously^4^. In **A-C**, n=3 mice per timepoint and multiple tissues per organ/tumor/metastases have been analyzed. **(D)** Reactivation of TT-specific memory CD4 and CD8 T cells in KPC mice post tumor development through the *Listeria*-TT+GEM treatment cycle described in Fig S1B. In D, T cell-depleted groups were compared to non-depleted groups.

**Fig. S4:**
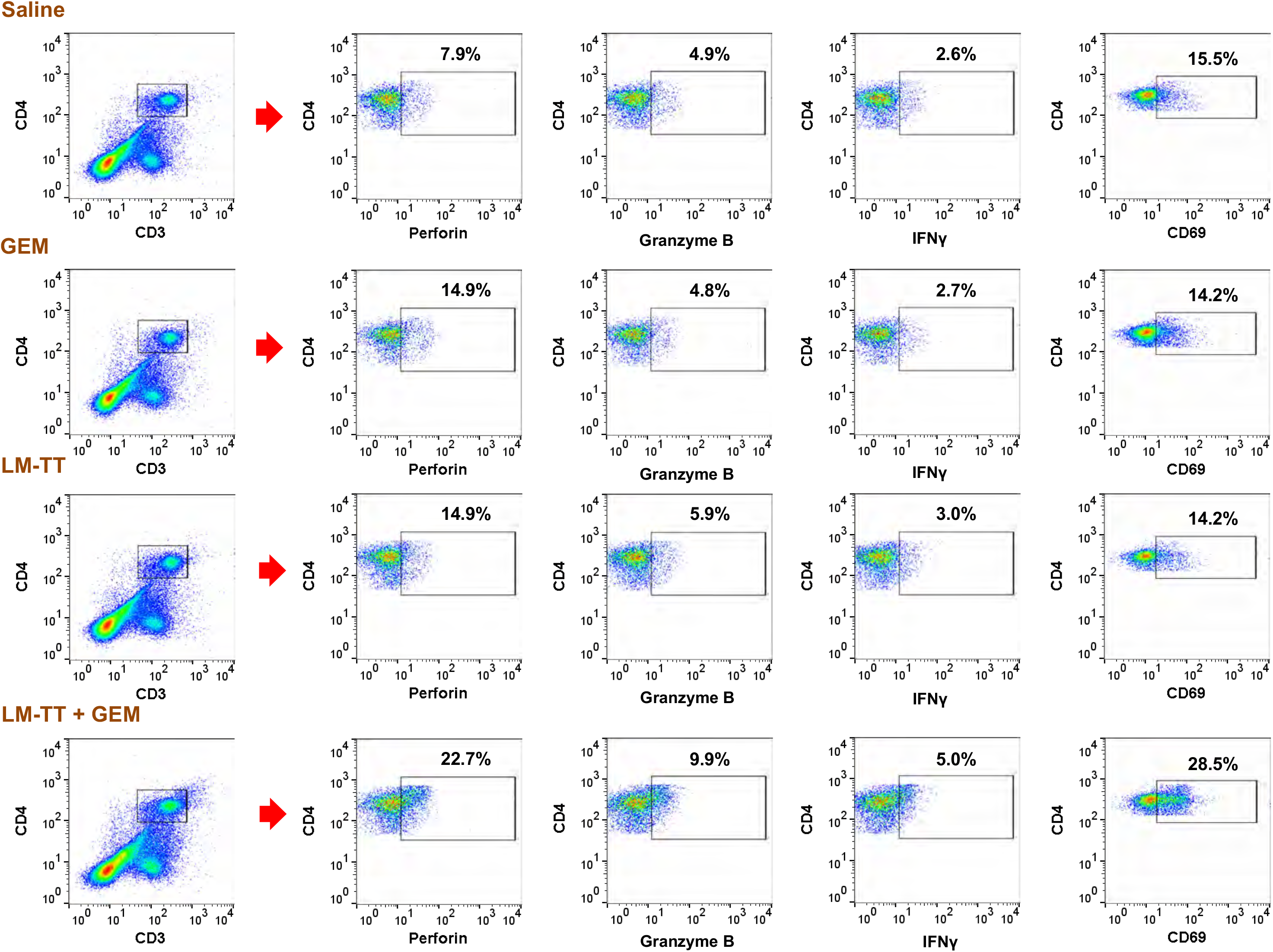
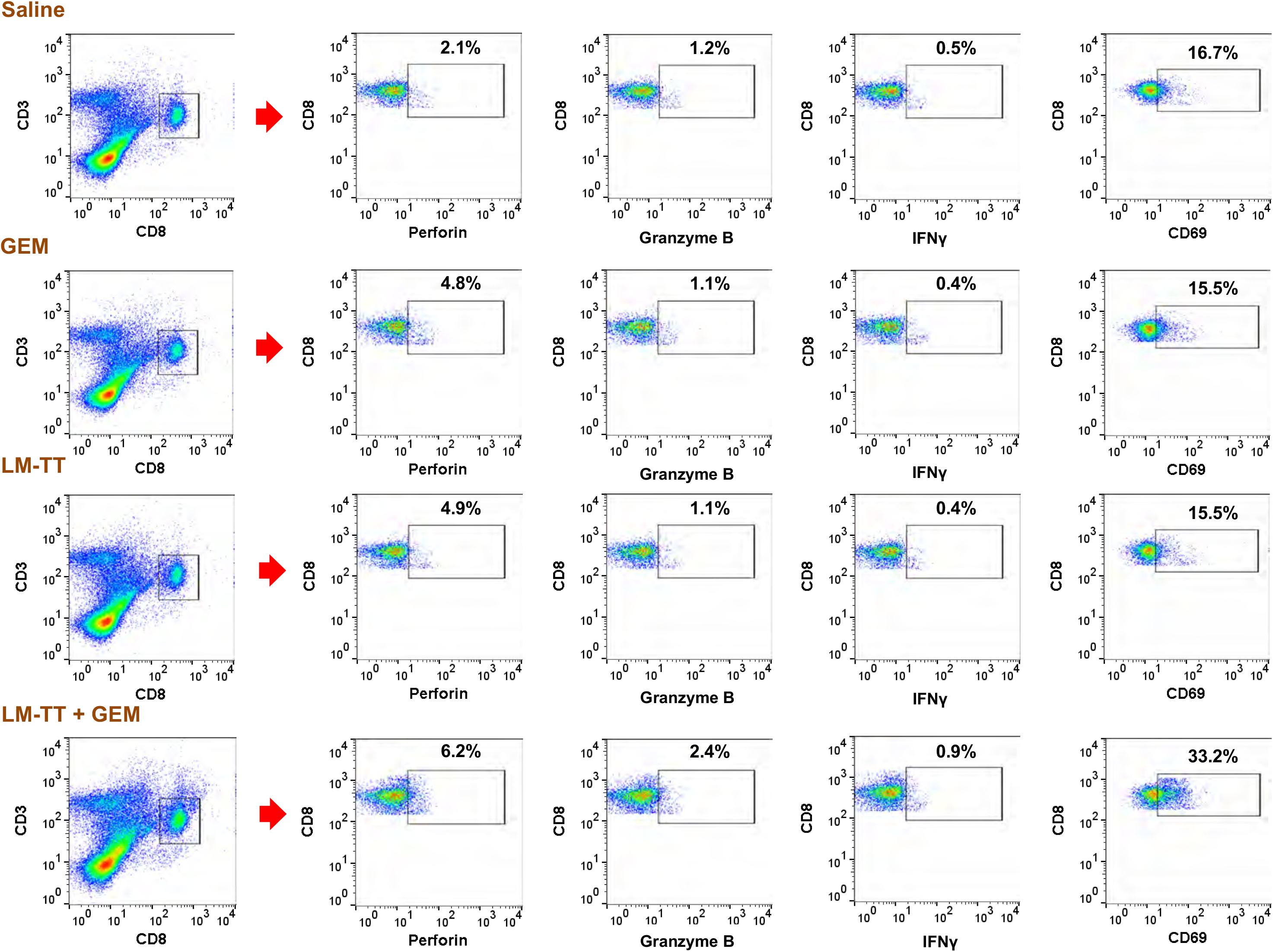
Gating of CD4 and CD8 T cells by flow cytometry in spleens of tumor-bearing Panc-02 mice treated with *Listeria*-TT and GEM, in combination or separately. Examples of gating **(A)** the CD4 T cells and **(B)** CD8 T cells isolated from spleens of treated and control mice. CD3CD4-pos cells and CD3CD8-pos cells were gated, followed by gating of CD4 or CD8 T cells producing IFNγ, Perforin, Granzyme B, or expressing CD69. A similar gating protocol was used to analyze CD4 and CD8 T cells in spleens of KPC mice. LM-TT = *Listeria*-TT.

**Fig. S5:**
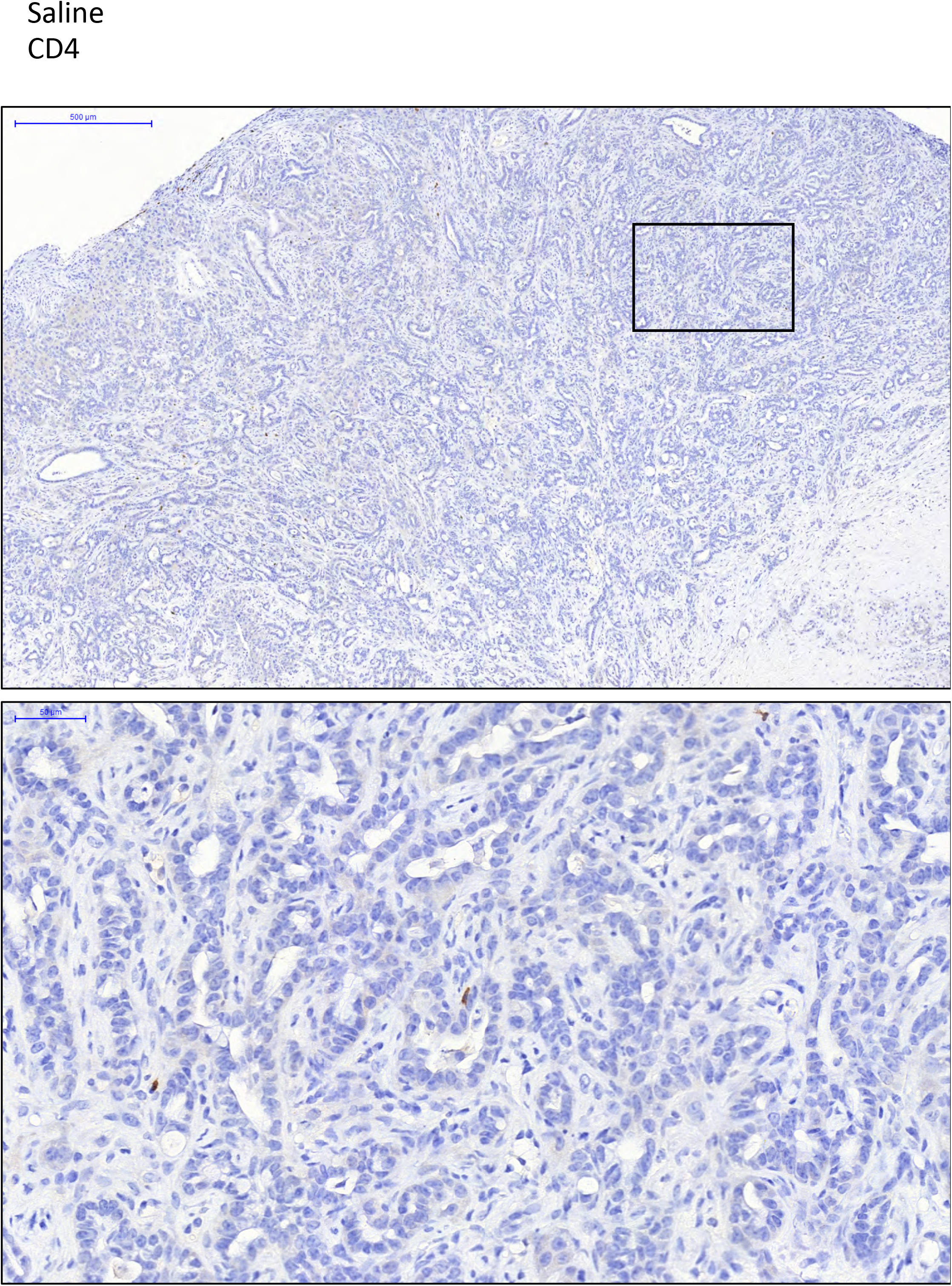
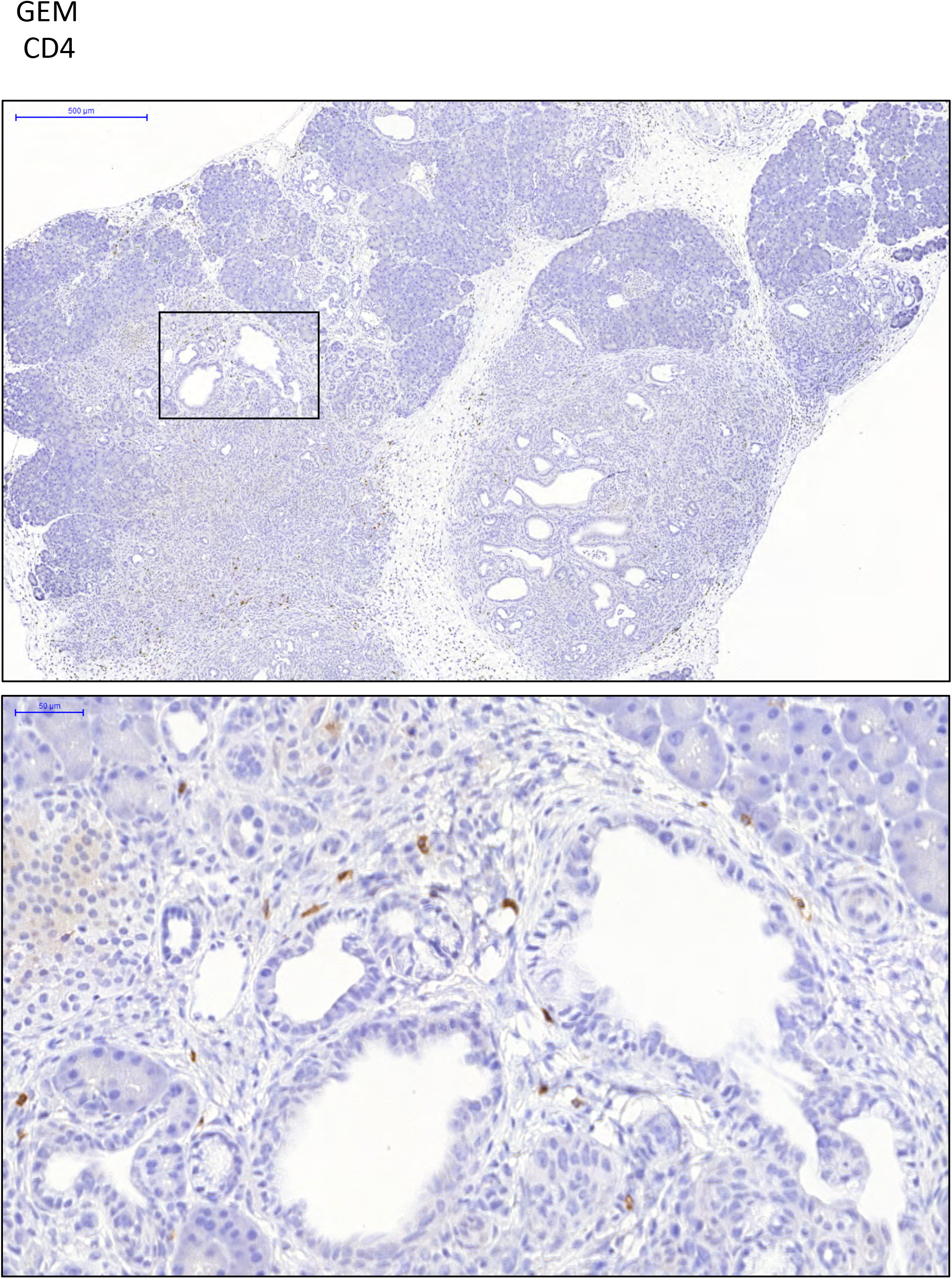
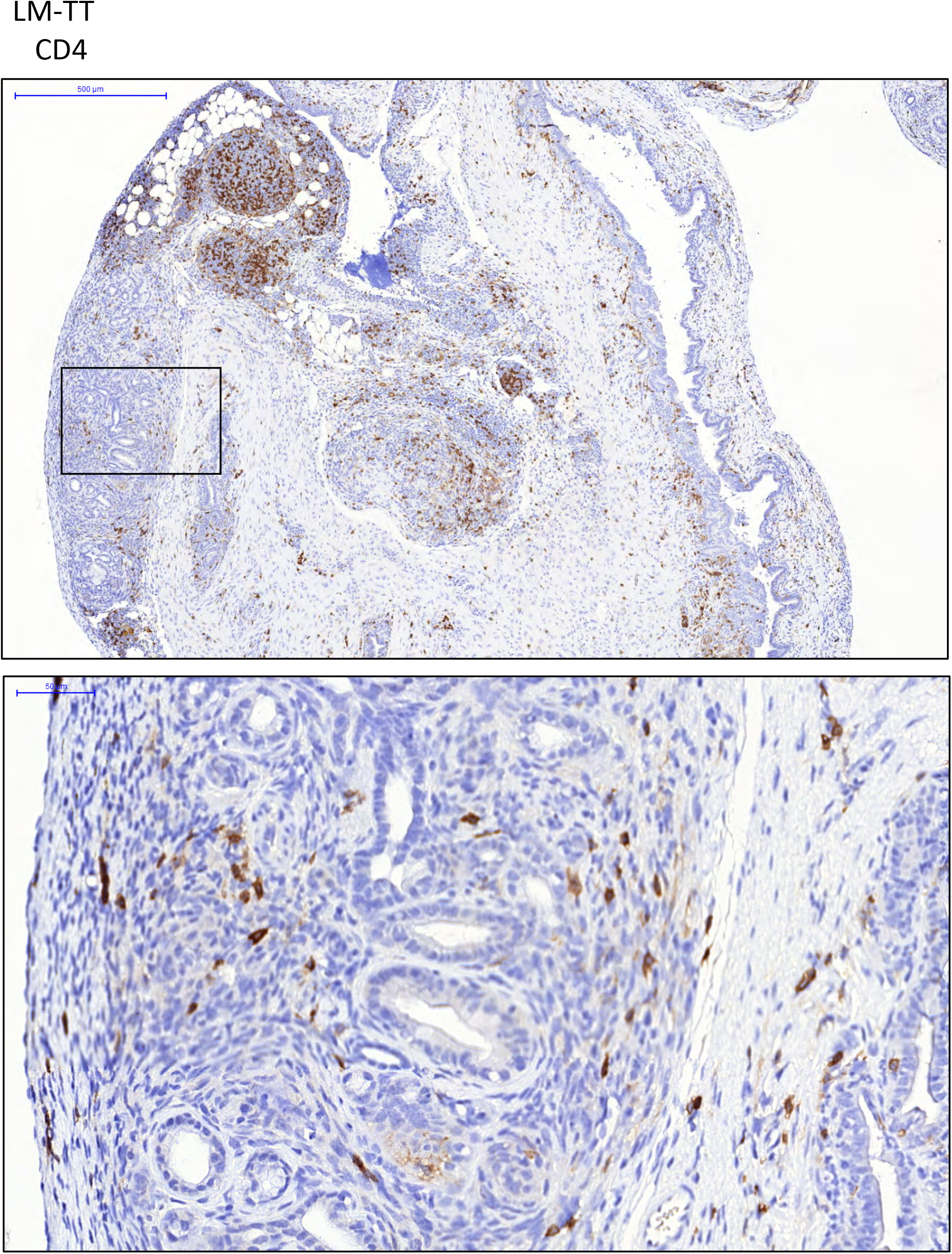
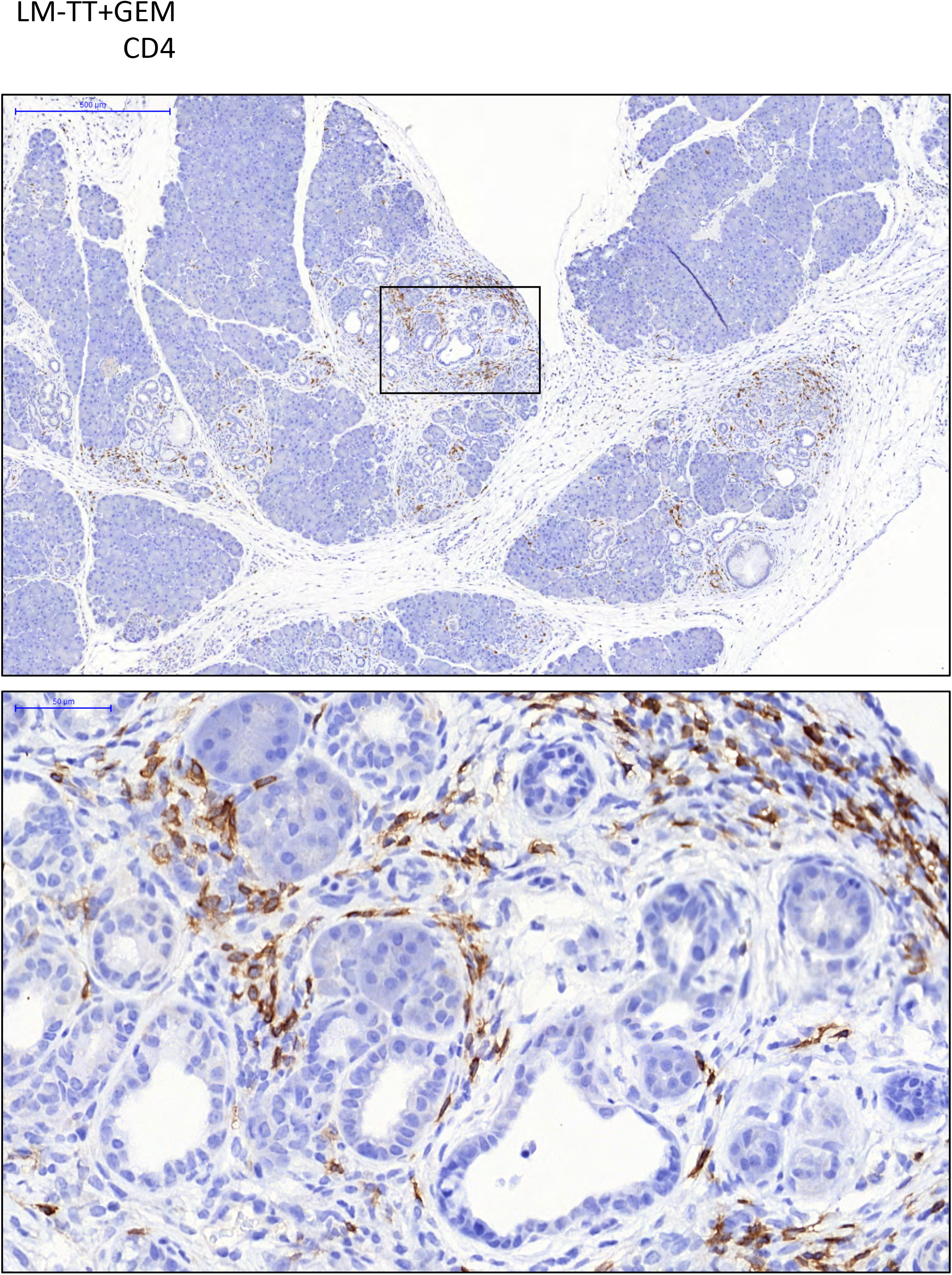
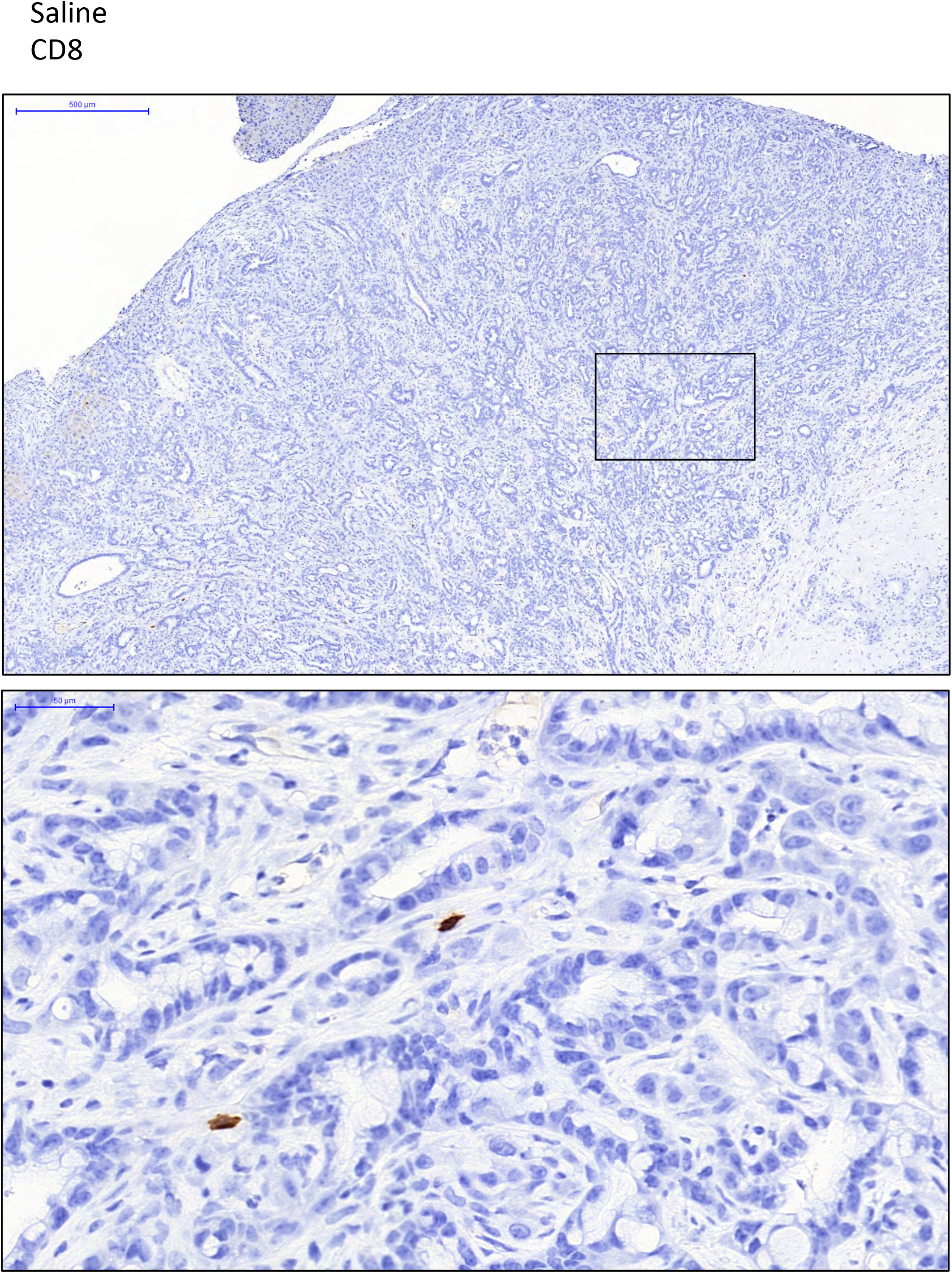
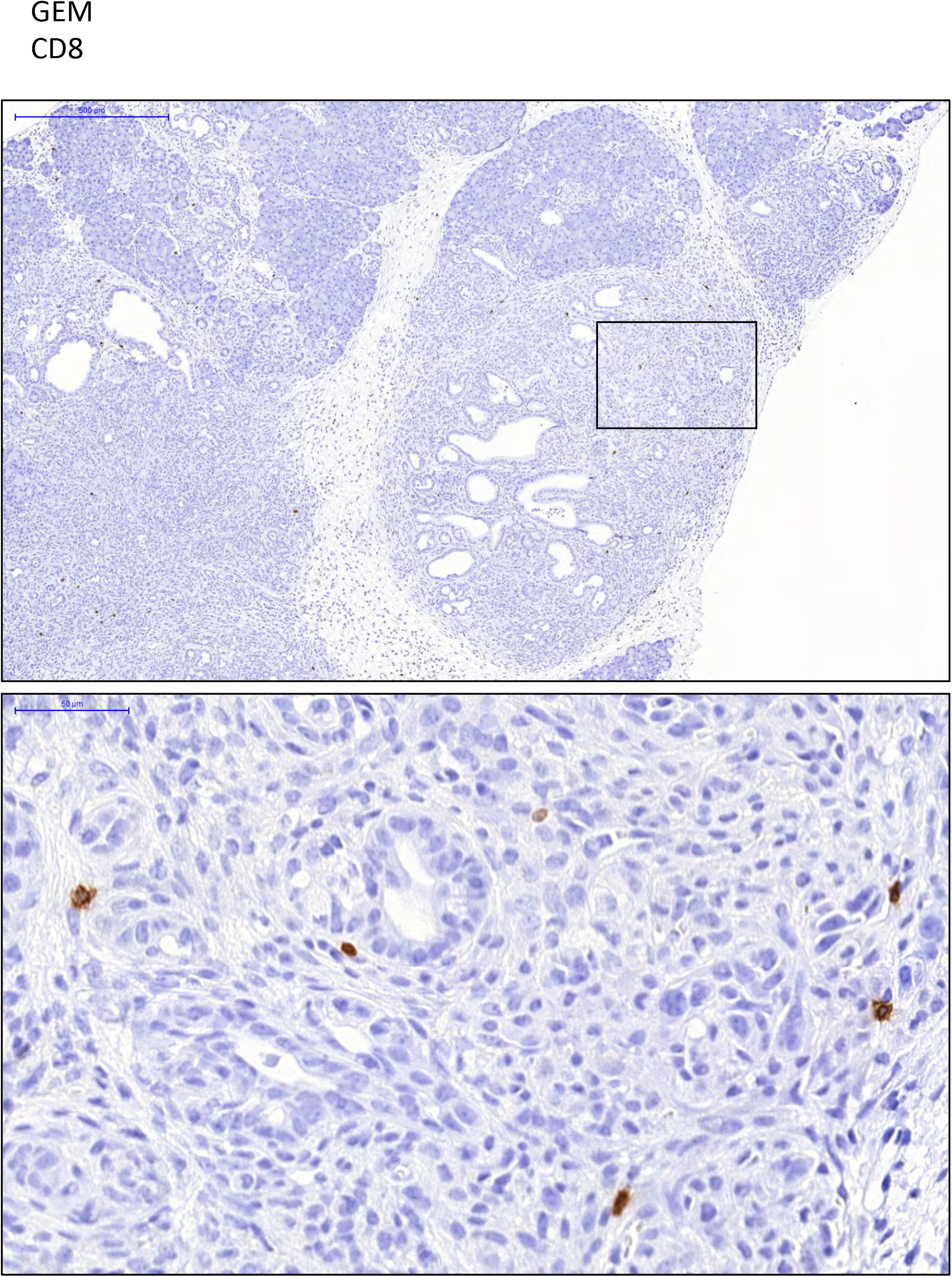
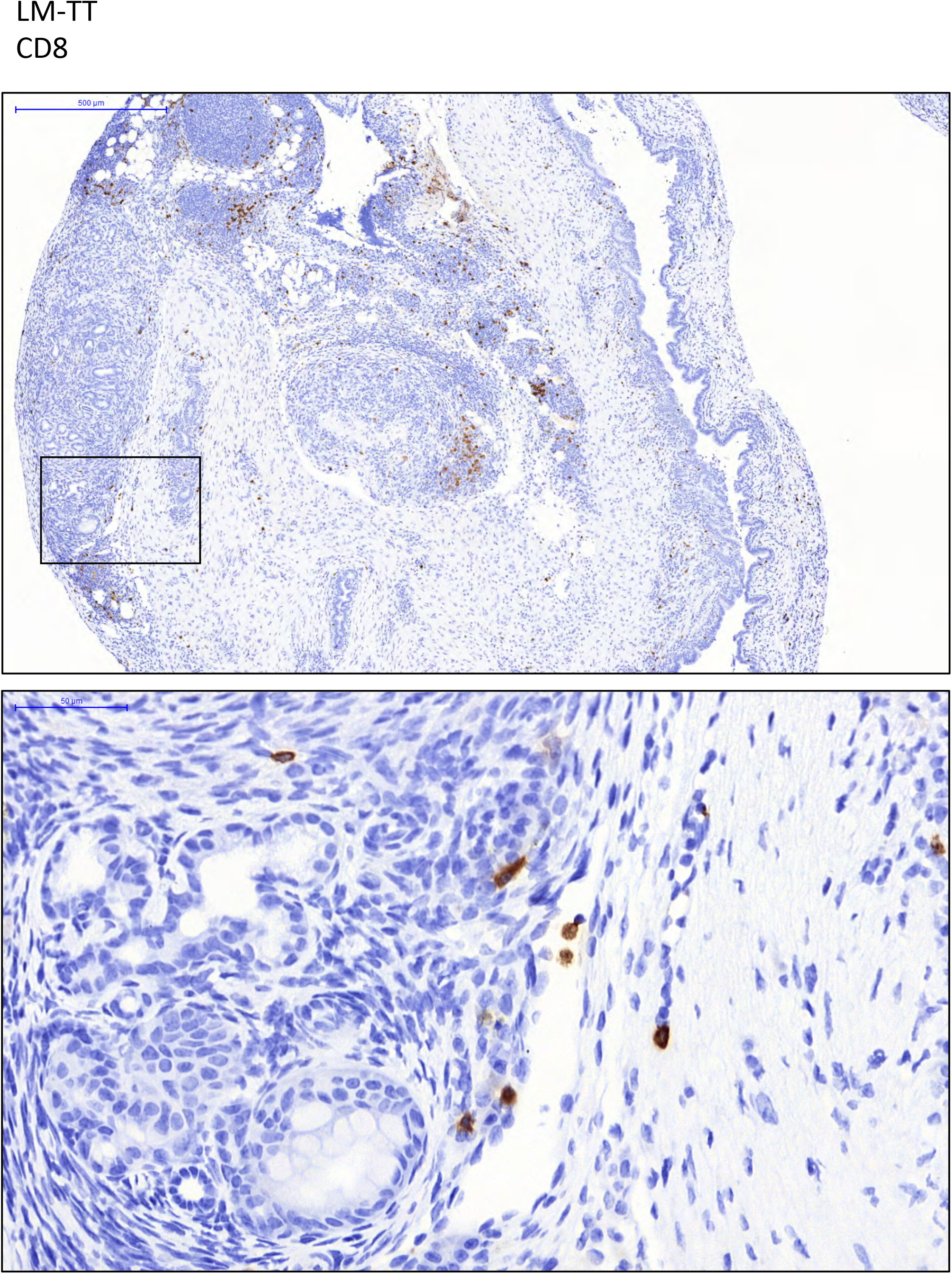
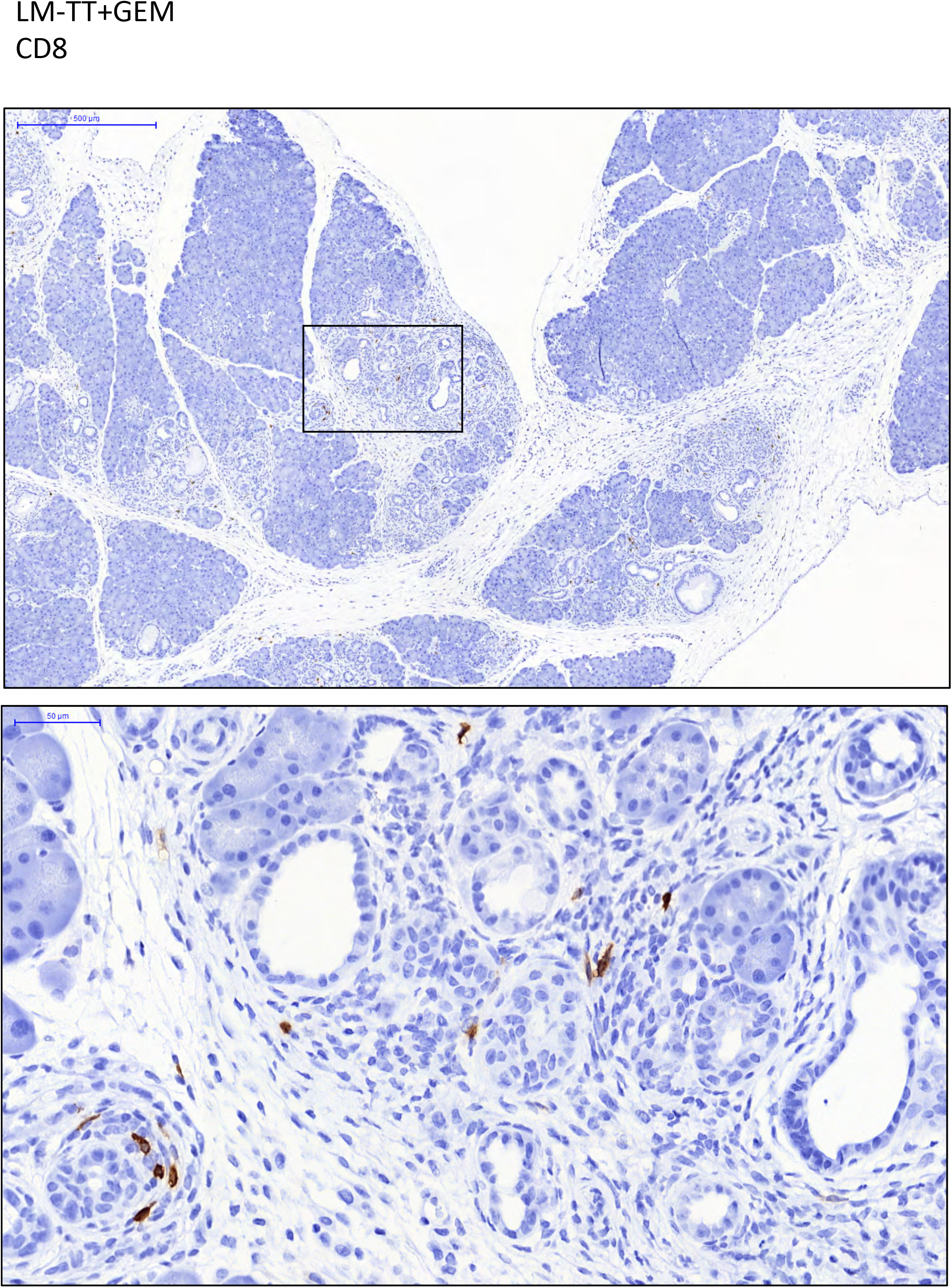
Analysis of CD4 and CD8 T cells in tumors of *Listeria*-TT+GEM-treated and control mice by IHC. KPC mice received treatment with *Listeria*-TT+GEM, or control treatments, as outlined in Fig 1B. Two days after the last treatment, tumor tissues were analyzed by IHC for the presence of CD4 **(A-D)** and CD8 **(E-H)** T cells. CD8, and particularly CD4 T cells, were predominantly observed in the tumor areas of KPC mice that received *Listeria*-TT or *Listeria*-TT+GEM. LM-TT = *Listeria*-TT.

**Fig. S6:**
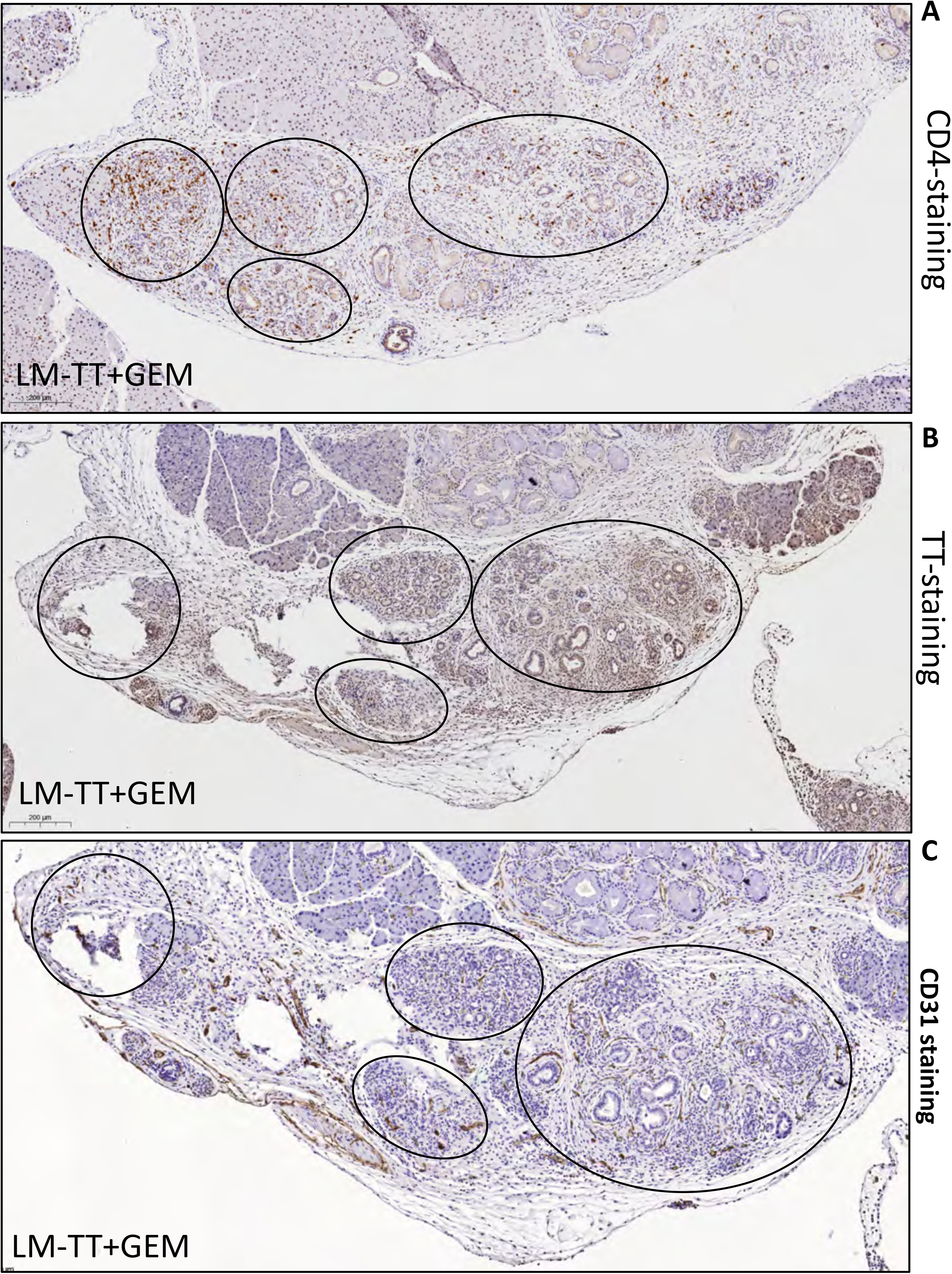
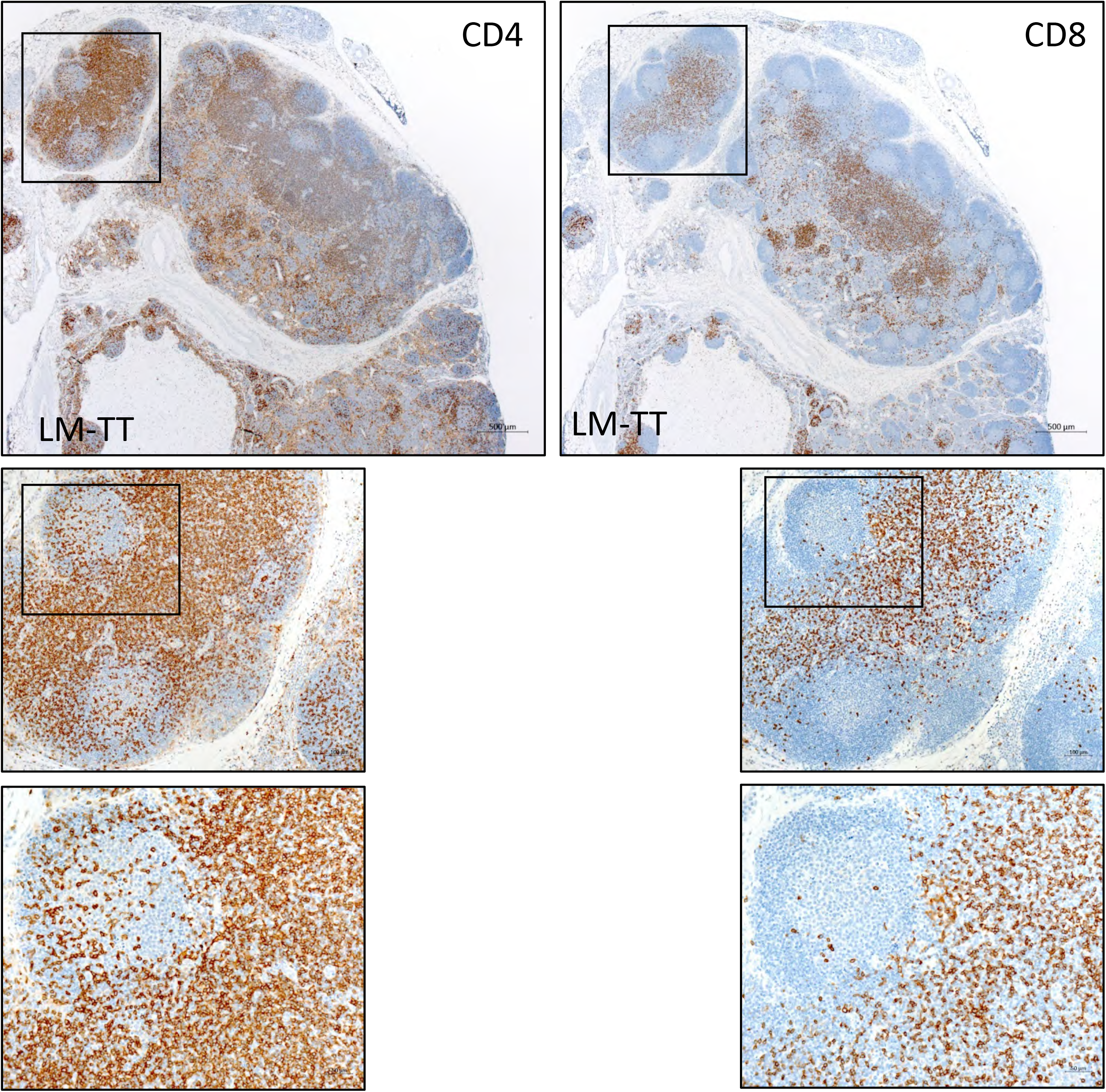
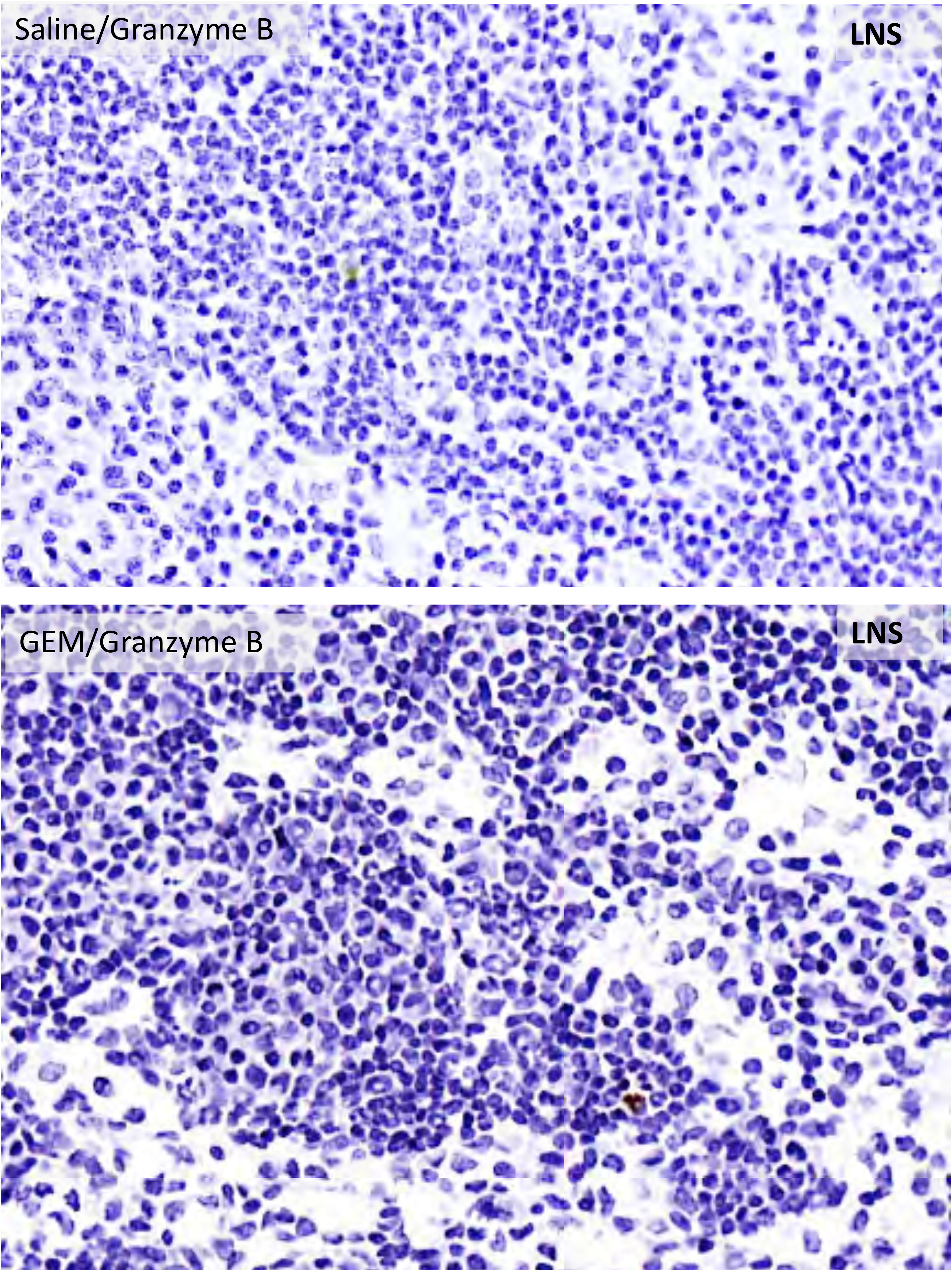
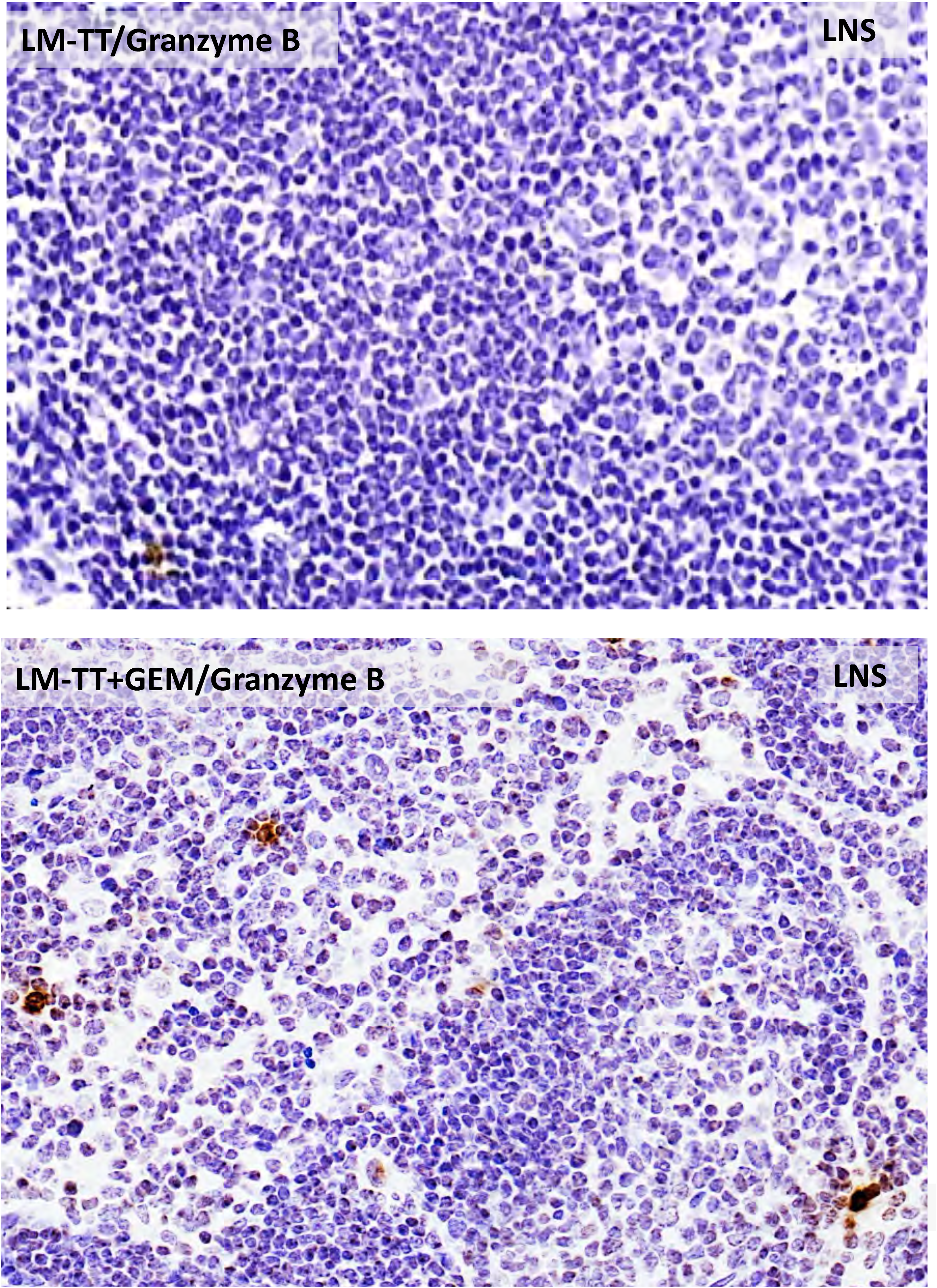
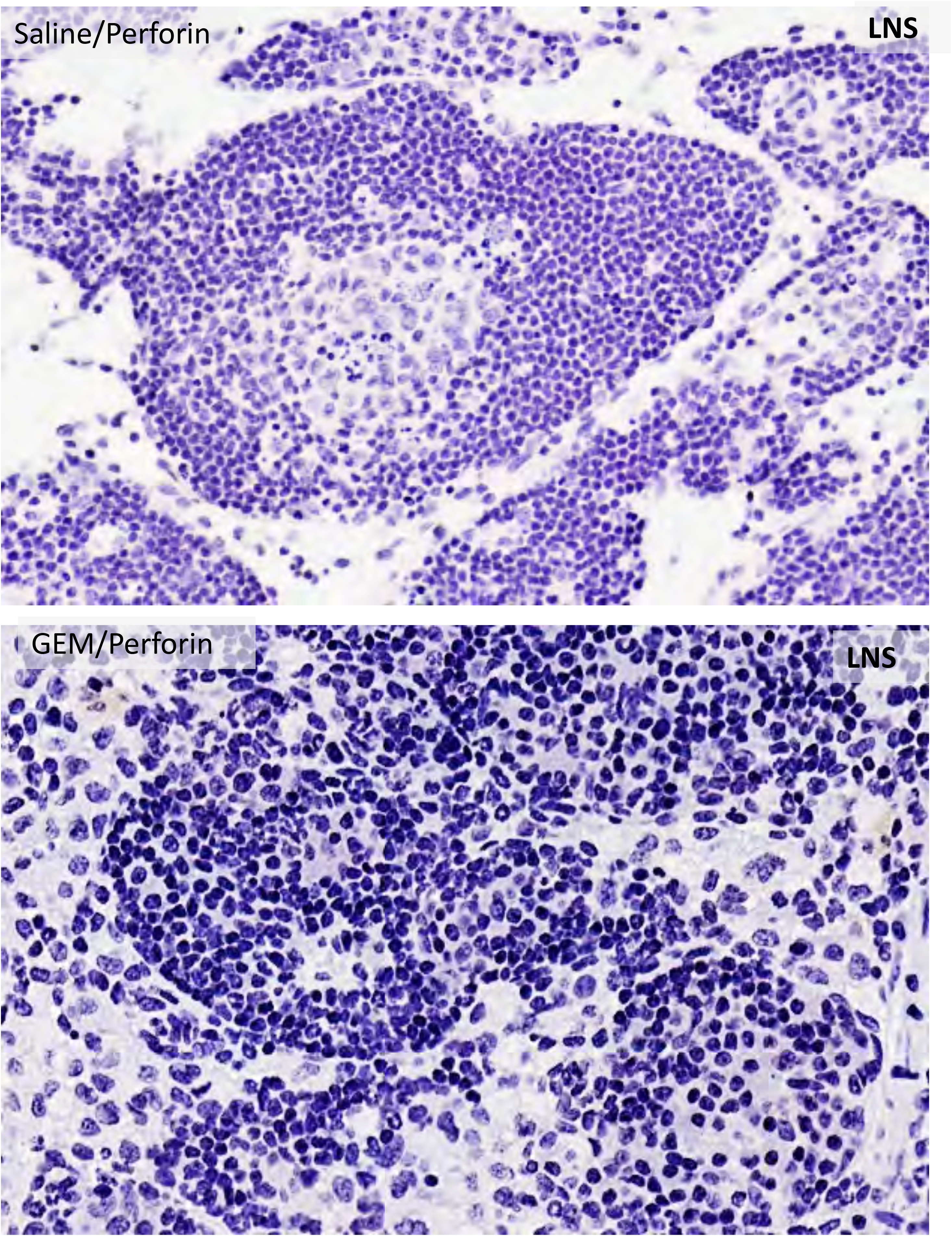
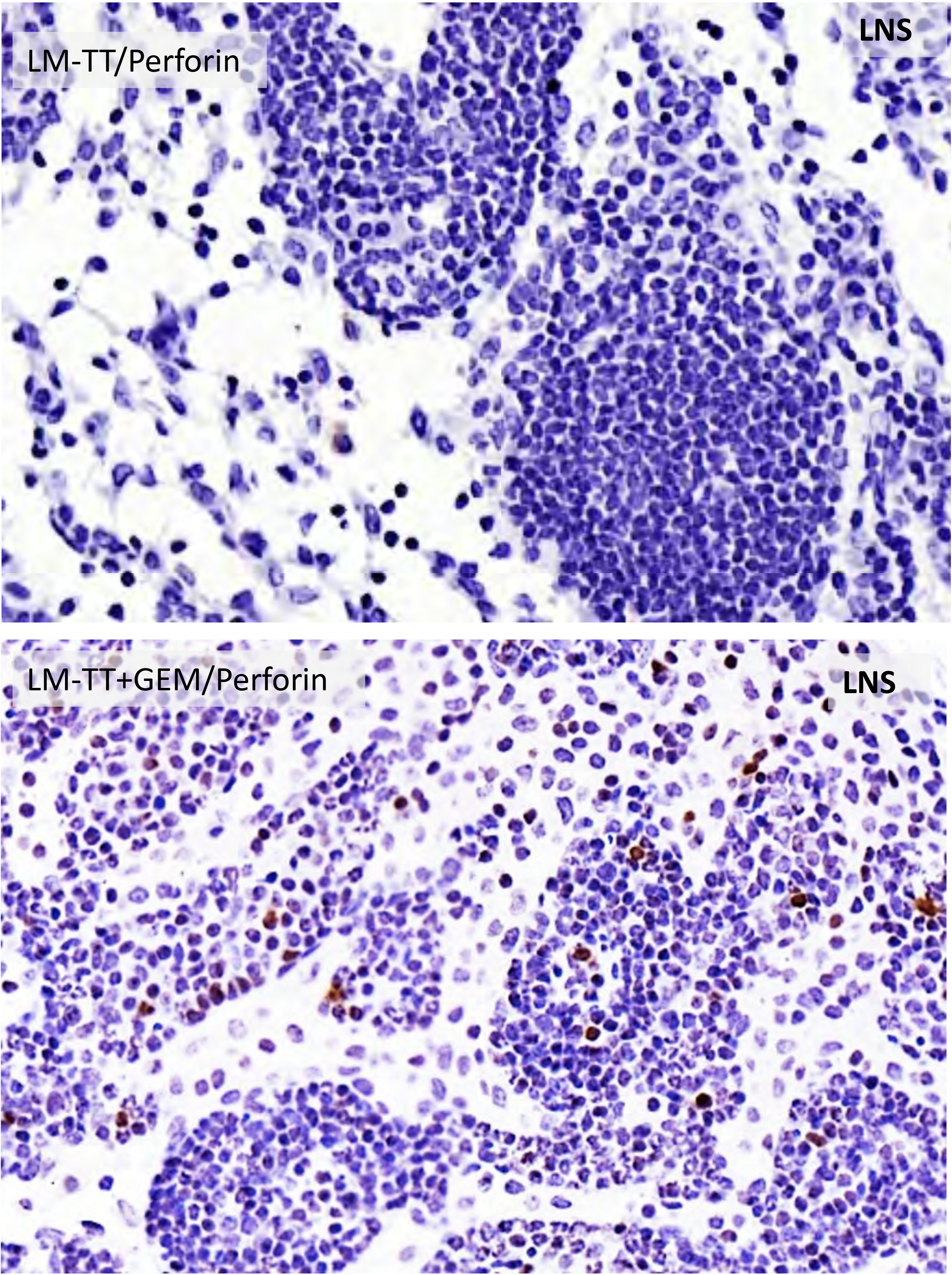
Analysis of KPC tumors and LNS by IHC. CD4 T cells **(A)** in tumor areas with TT (**B)** and CD31 **(C)** expression by IHC of KPC mice treated with *Listeria*-TT+GEM. Circles mark tumor areas. **(D)** Example of LNS in close proximity to a pancreatic tumor of a KPC mouse. In the overview of the lymphoid structure, there were well-developed follicular structures, many of which contained a pale germinal area at the center. CD4+ T cells were concentrated in the cortex and in the deep cortical zone/paracortex, where they were arranged in sheets rather than follicles. CD8+ T cells were concentrated within the deep cortical zone/paracortex. Below are views of the lymphoid structure at higher magnification, to visualize the density of CD4+ and CD8+ T cells. CD4+ cells were located mostly within the cortex, surrounding and within the follicles, and in the paracortex. CD8+ cells were primarily within the paracortex. **(E-H)** Production of Granzyme B and Perforin in LNS of KPC mice treated with saline, GEM, *Listeria*-TT, or *Listeria*-TT+GEM, detected by IHC. LM-TT = *Listeria*-TT.

**Fig. S7:**
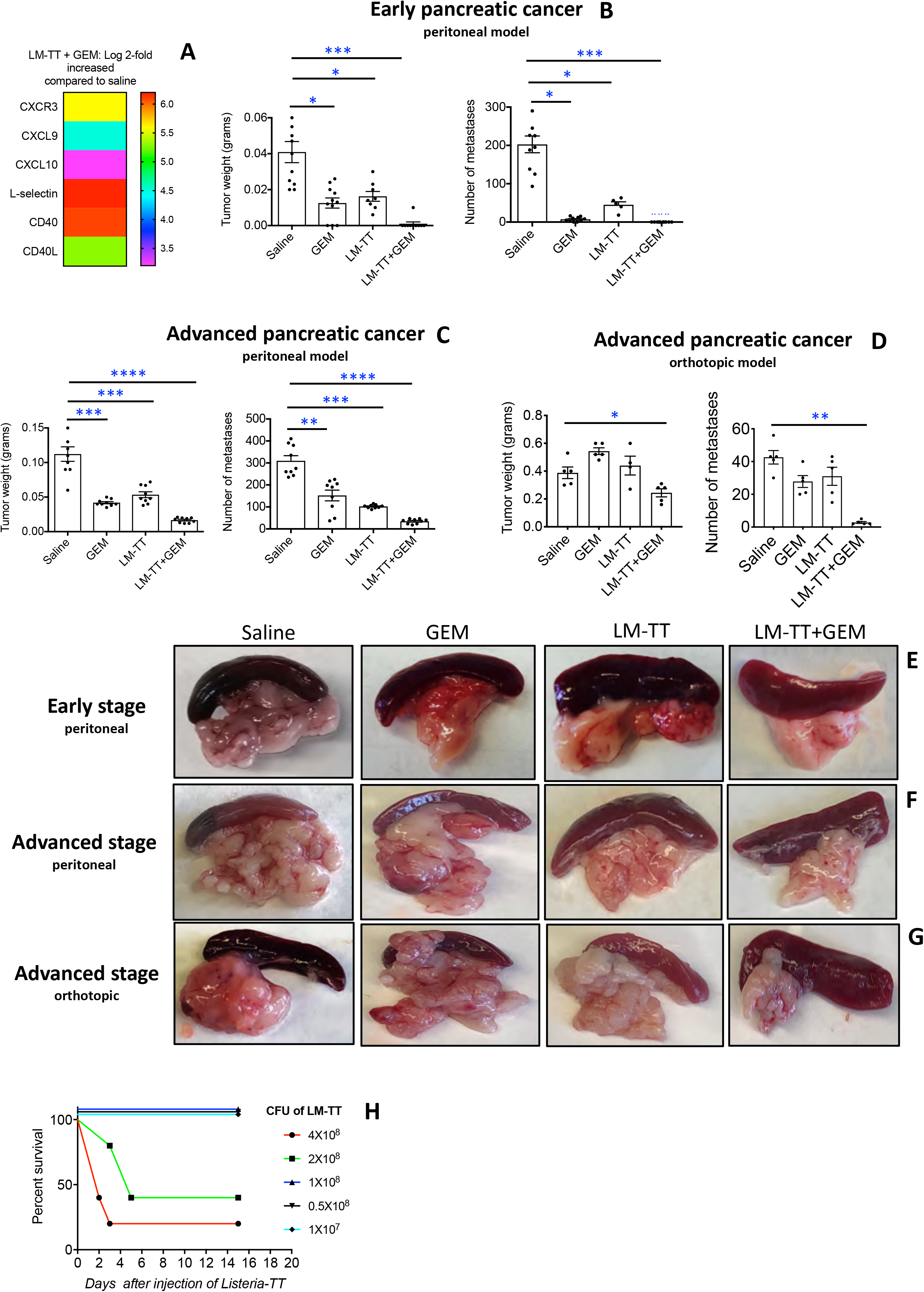
Effect of LM-TT+GEM treatment on expression of genes involved in T cell migration, efficacy and safety. **(A)** Detection of chemokines and their receptors in pancreatic tumors of *Listeria* TT+GEM-treated or untreated KPC mice by RNAseq. Results of 2 mice are shown in each group. Log2-fold increase in the group of *Listeria*-TT+GEM compared to the untreated mice. The heatmap was generated by unbiased hieratical clustering at the following statistical parameters. Statistical parameters for the paired-wise comparison: p<0.05. **(B)** LM-TT+GEM testing in the Panc-02 mouse peritoneal cavity model of early pancreatic cancer. Treatments began three days after tumor cell injection as outlined in Fig 1B. Mice were euthanized after treatment, and analyzed for tumor weight and the number of metastases. Two independent experiments with 5 mice per group were averaged. **(C)** LM-TT+GEM testing in the Panc-02 mouse peritoneal cavity model of advanced pancreatic cancer. Treatments began ten days after tumor cell injection as outlined in Fig 1B. Two independent experiments with 5 mice per group were averaged. **(D)** LM-TT+GEM testing in the orthotopic Panc-02 mouse model of advanced pancreatic cancer. Treatments were started 14 days after tumor cell injection as outlined in Fig 1B. Averages are shown for a single experiment with 5 mice in each group. Error bars represent standard error of the mean. Mann-Whitney *p<0.05, *** p<0.01. Examples of pancreatic metastases and tumors are shown of early pancreatic cancer in the peritoneal Panc-02 model **(E)**, advanced pancreatic cancer in the peritoneal Panc-02 model **(F)**, and advanced pancreatic cancer in the orthotopic Panc-02 model **(G). (H)** Dose-limiting toxicity study of *Listeria*-TT in tumor-naive C57Bl6 mice (n=5 mice per group). LM-TT = *Listeria*-TT. In B, C and D, all groups were compared to the saline group.

**Fig. S8:**
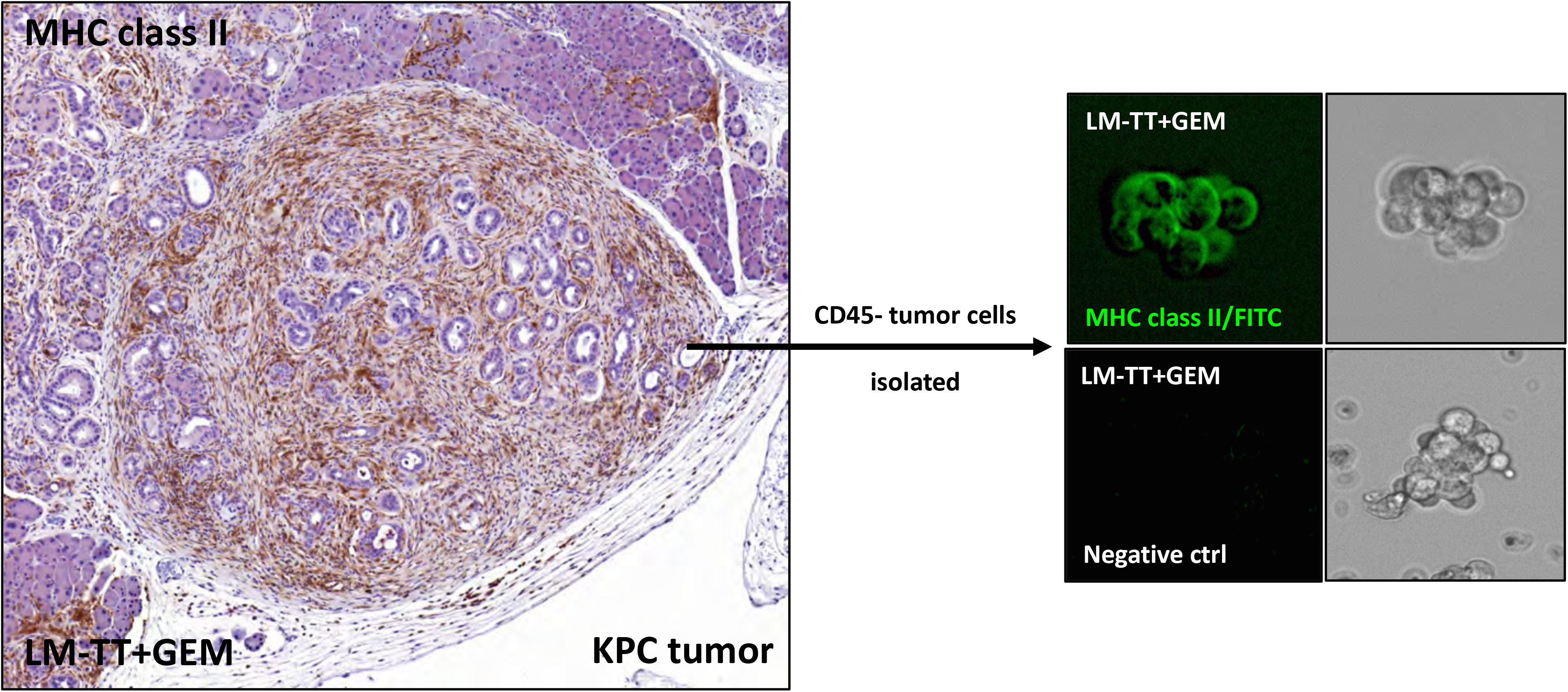
Detection of MHC class II in KPC tumors by IHC and IF. Tumors of KPC mice treated with *Listeria*-TT+GEM were analyzed for the presence of MHC class II expression by IHC. In addition, CD45-negative KPC cells were sorted from the tumors using magnetic beads and analyzed for MHC class II expression by immunofluorescence (IF). LM-TT = *Listeria*-TT.

**Fig. S9:**
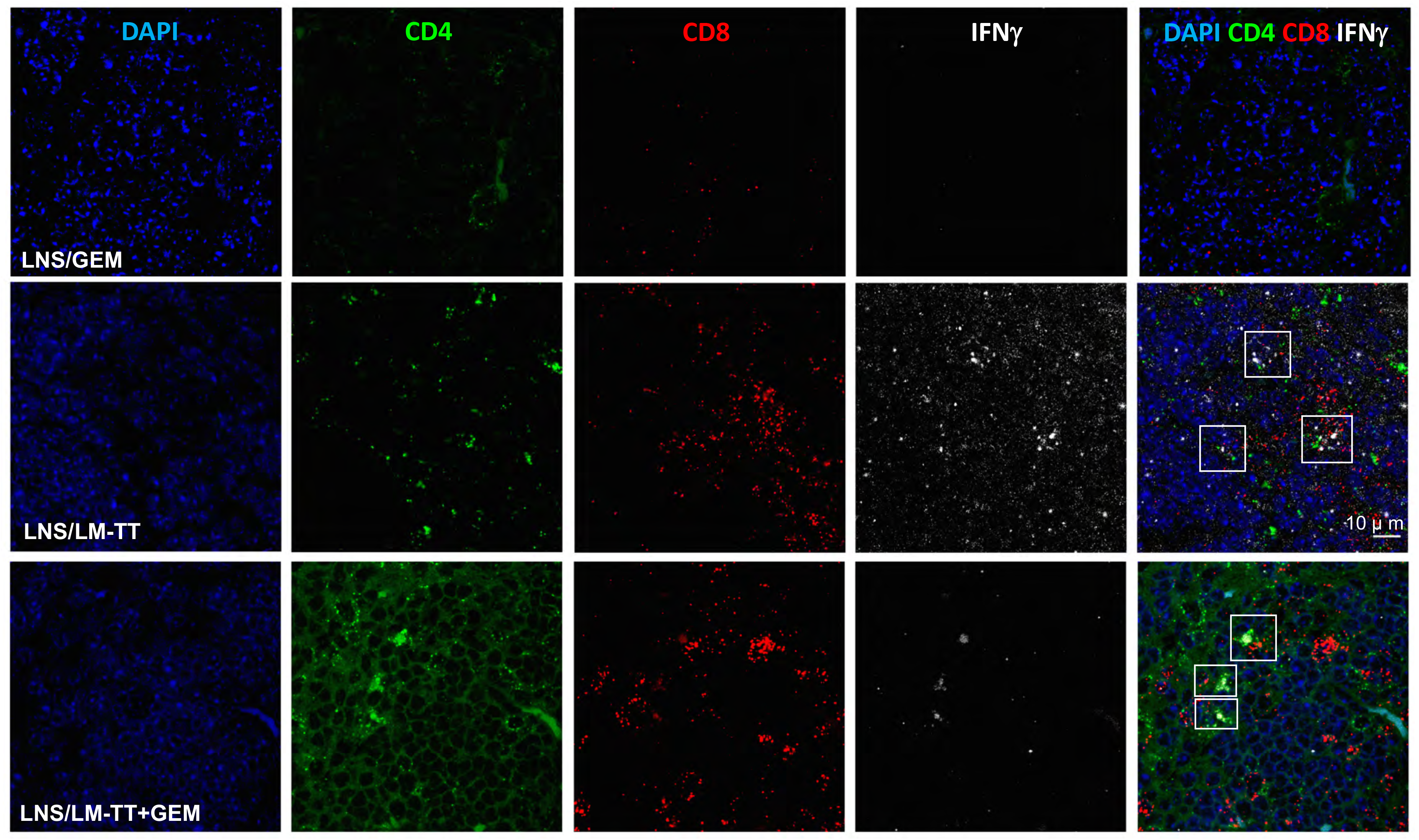
Analysis of IFNγ by CD4 and CD8 T cells in LNS of KPC mice by RNAscope. To confirm the flow cytometry data in Fig 4B, we analyzed the production of IFNγ by CD4 and CD8 T cells in LNS of KPC mice treated GEM, LM-TT, or LM-TT+GEM by RNAscope. The saline group did not exhibit LNS. Tissues were analyzed by HALO Image Analysis. CD4 T cells are green, CD8 T cells red, IFNγ white and the nucleus blue. LM-TT = *Listeria*-TT.

**Fig. S10:**
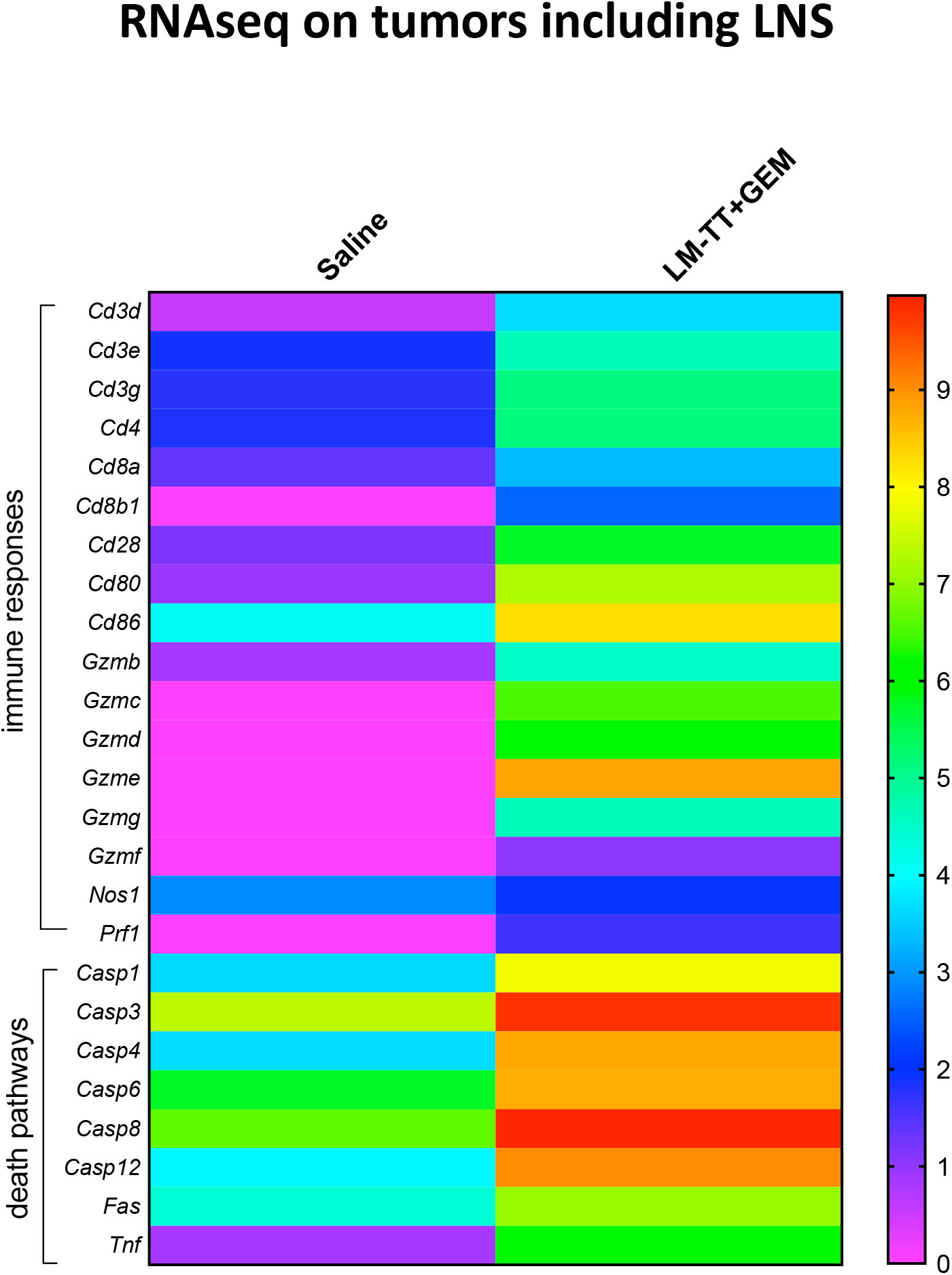
Analysis of KPC tumors including LNS by RNAseq. RNAseq analysis of immune responses and tumor cell death pathways in pancreatic tumors including LNS, of KPC mice in response to *Listeria*-TT+GEM treatment and saline control group. The results of 2 mice in each group was averaged. The heatmap was generated by unbiased hieratical clustering at the following statistical parameters. Statistical parameters for the paired-wise comparison: p<0.05. The signature contains 25 important genes.

**Fig. S11:**
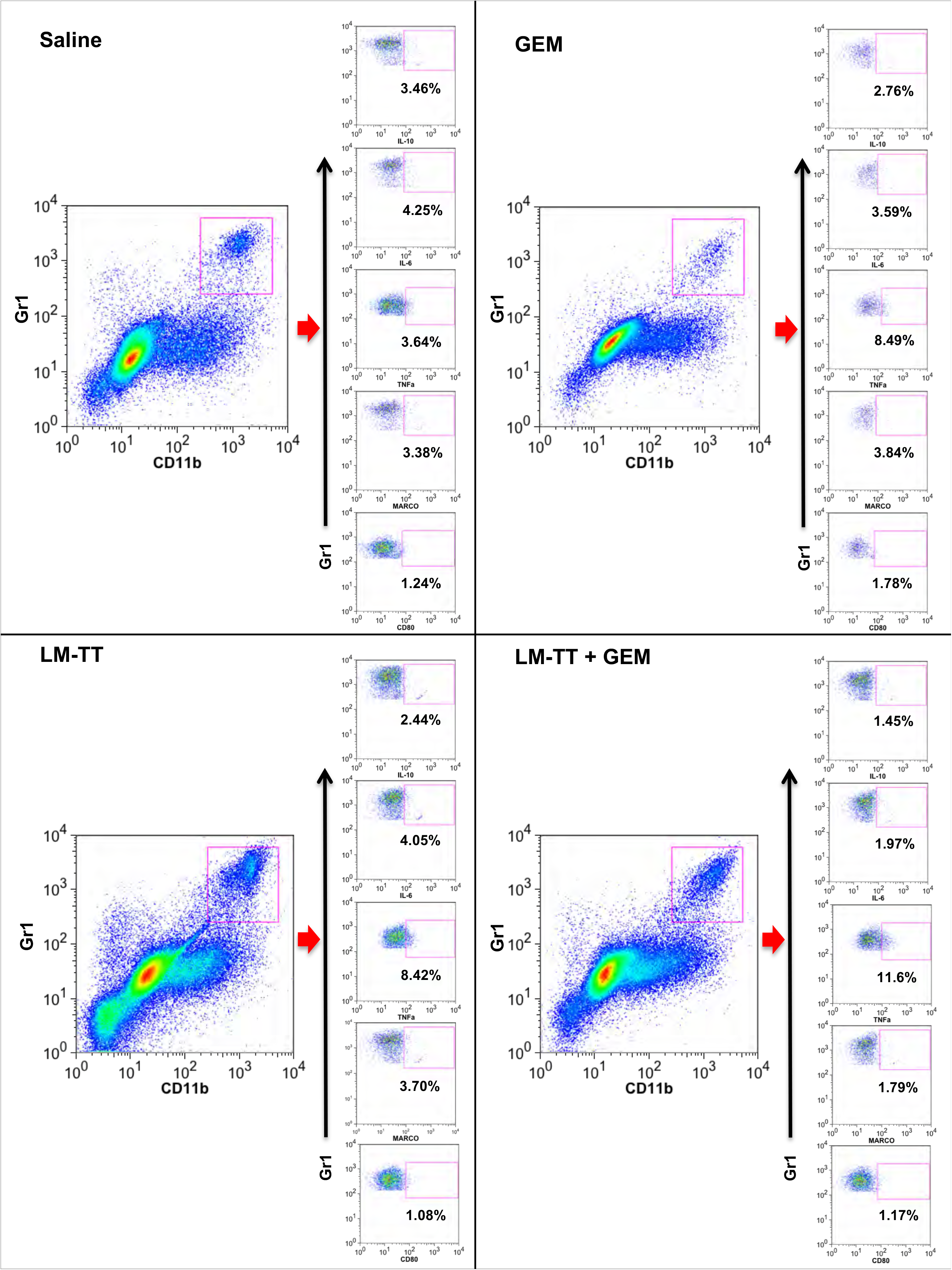
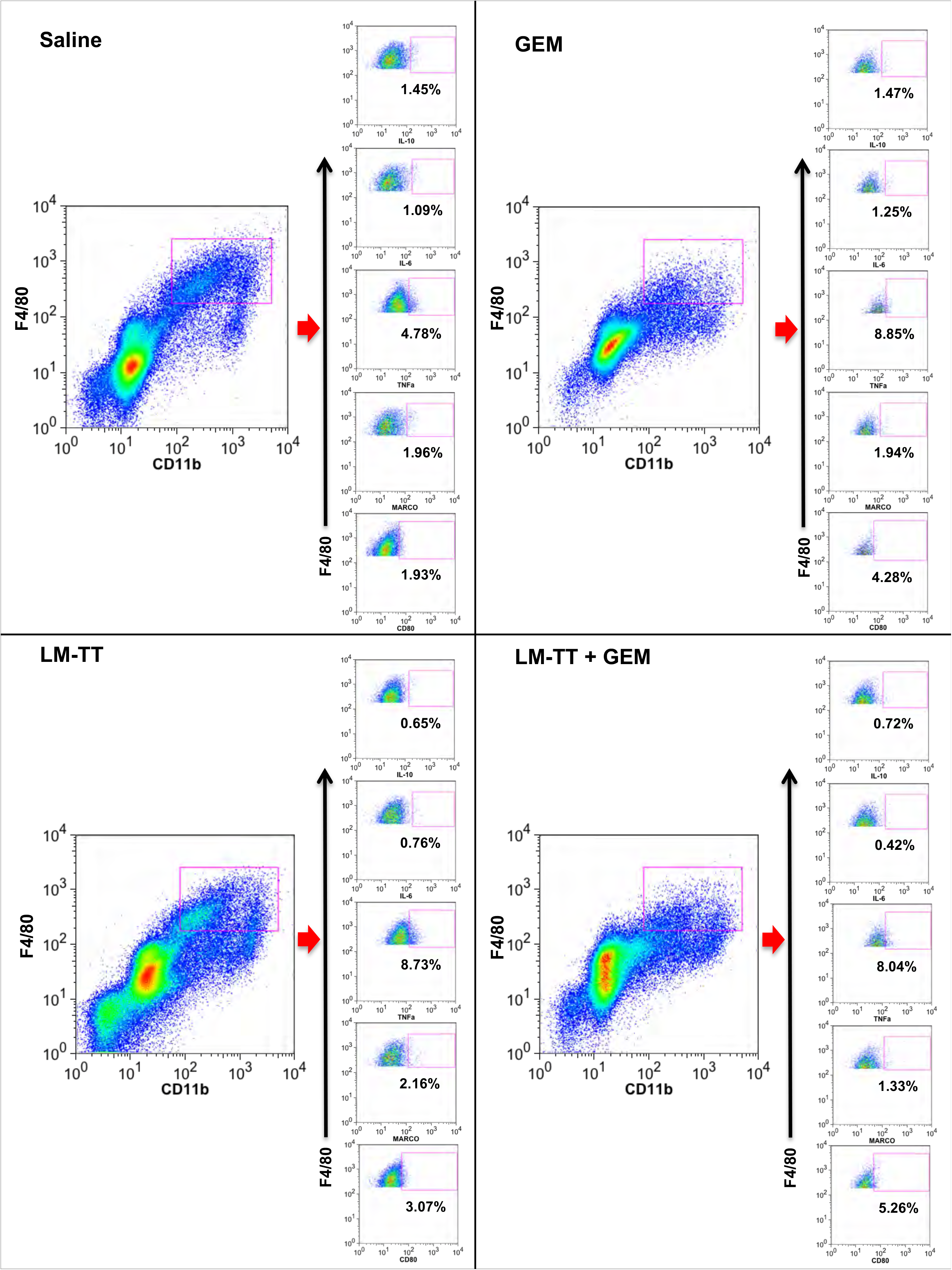
Gating of MDSC and TAM in metastases of tumor-bearing Panc-02 mice. **(A)** Example of gating of MDSC cells present in metastases of treated and control mice. First, live cells were gated followed by CD45+ leukocytes to exclude tumor cells. Subsequently, CD11bGr1-pos cells (MDSC) were gated in the live CD45+ cell population, followed by the gating of Gr1-pos cells producing IL-10, IL-6, TNFα, or expressing MARCO or CD80 as indicated in the figure. **(B)** Example of gating TAMs present in metastases of treated and control mice. First, live cells were gated followed by CD45+ leukocytes to exclude tumor cells. Subsequently, CD11bF4/80-pos cells were gated, followed by gating of F4/80-pos cells producing IL-10, IL-6, TNFα, or expressing MARCO or CD80.

---

**SI Table 1:** Number of CFU of Listeria in mouse Panc--02 tumors and normal tissues of C57Bl/6 mice (Average ± SEM)

| Tissue | Day 1 | Day 3 | Day 7 |
| --- | --- | --- | --- |
| Blood | 0.0 ± 0.0 | 0.0 ± 0.0 | 0.0 ± 0.0 |
| Spleen | 7541.6 ± 3049 | 6215.6 ± 1820 | 0.0 ± 0.0 |
| Kidney | 812.5 ± 235 | 0.0 ± 0.0 | 0.0 ± 0.0 |
| Lungs | 125.0 ± 81.8 | 0.0 ± 0.0 | 0.0 ± 0.0 |
| Liver | 298.6 ± 109.5 | 1585.7 ± 744.7 | 0.0 ± 0.0 |
| Heart | 0.0 ± 0.0 | 0.0 ± 0.0 | 0.0 ± 0.0 |
| GI | 0.0 ± 0.0 | 0.0 ± 0.0 | 0.0 ± 0.0 |
| LN | 102.6 ± 50.2 | 0.0 ± 0.0 | 0.0 ± 0.0 |
| Bone marrow | 166.6 ± 83.3 | 0.0 ± 0.0 | 0.0 ± 0.0 |
| Tumor | 80375.0 ± 4336.7 | 466.6 ± 79 | 0.0 ± 0.0 |
| Liver Mets | 14331.2 ± 5740 | 100.0 ± 56.6 | 1023.8 ± 517.4 |
| Panc Mets | 19562.5 ± 59581 | 46782.7 ± 7967.8 | 3875.0 ± 1485.9 |
| GI Mets | 10150.0 ± 1426.8 | 6619.9 ± 1829.5 | 0.0 ± 0.0 |

Panc-02 mice were injected ip with $1 \times 10^7$ colony-forming units (CFU) of *Listeria*-TT and euthanized on days 1, 3, and 7. The number of CFU of *Listeria* was determined per gram tissue, and the results of 3 mice were averaged. GI; Gastrointestinal; LN, lymph nodes, Mets; metastases.

**Table S2:** Number of CFU of Listeria in human MiaPaCa2 tumors and normal tissues in nude mice (Average ± SEM)

| **Tissue** | **Day 1** | **Day 3** | **Day 7** |
| --- | --- | --- | --- |
| **Blood** | 26.7 ± 17.6 | 30 ± 18.5 | 8.9 ± 8.9 |
| **Spleen** | 766.3 ± 290.1 | 5994.1 ± 2486.3 | 2388.9 ± 367.3 |
| **Kidney** | 75.6 ± 32.7 | 301.9 ± 116.9 | 167.6 ± 35.0 |
| **Lungs** | 229.6 ± 55.1 | 3906.3 ± 3445.1 | 586.1 ± 231.8 |
| **Liver** | 1,270.0 ± 550.4 | 1,577.5 ± 673.2 | 529.6 ± 177.7 |
| **Heart** | 19.4 ± 8.1 | 35.2 ± 18.1 | 827.8 ± 423.5 |
| **GI** | 0.0 ± 0.0 | 66.7 ± 47.1 | 88.9 ± 30.9 |
| **LN** | 6,446.7 ± 3242.7 | 114.8 ± 52.4 | 277.8 ± 60.6 |
| **Bone Marrow** | 0.0 ± 0.0 | 2.2 ± 2.2 | 0.0 ± 0.0 |
| **Tumor** | 8,311.1 ± 1785.1 | 123,240.4 ± 30429.9 | 166,666.7 ± 1825.7 |
| **Liver Mets** | 3,300.0 ± 911.1 | 333,333.4 ± 74535.6 | 502,33.3 ± 22256.4 |

Nude mice (NU/J) with human Mia-PaCa2 tumors and metastases (Mets) were injected with $5 \times 10^7$ colony forming units (CFU) of Listeria and euthanized on days 1, 3, and 7. The number of CFU of Listeria was determined per gram tissues, and the results of 3 mice were averaged. Gastrointestinal, GI; LN, lymph nodes.

**Table S3:** Pathological examination of tissues in C57Bl/6 mice 2 days after the last treatment with Listeria-TT and Gemcitabine

| **Treatment** | **Saline** |  |  | **Listeria-TT+Gem** |  |  |
| --- | --- | --- | --- | --- | --- | --- |
| **Mouse** | **1** | **2** | **3** | **1** | **2** | **3** |
| **Liver** |  |  |  |  |  |  |
| *Random foci of mixed leukocytes* | 1 | 1 | 1 | 0 | 1 | 1 |
| *Individual hepatocyte cell death* | 1 | 0 | 0 | 0 | 1 | 1 |
| *Extramedullary hematopoiesis* | 0 | 0 | 0 | 0 | 0 | 1 |
| *Hepatocellular cytoplasmic vacuolation* | 0 | 0 | 0 | 0 | 0 | 0 |
| *Foci of necrosis* | 0 | 0 | 0 | 0 | 0 | 0 |
| **Kidney** |  |  |  |  |  |  |
| *Tubular ectasia, cortex* | 0 | 0 | 0 | 0 | 0 | 0 |
| *Tubular ectasia, medulla* | 0 | 0 | 0 | 0 | 0 | 1 |
| *Tubular epithelial cell cytoplasmic vacuolation, cortex* | 0 | 0 | 0 | 0 | 1 | 0 |
| *Tubular epithelial cell cytoplasmic vacuolation, medulla* | 0 | 0 | 0 | 0 | 0 | 0 |
| *Regenerative tubules, cortex* | 0 | 0 | 0 | 1-2 | 1-2 | 0 |
| *Regenerative tubules, medulla* | 0 | 0 | 0 | 0 | 0 | 0 |
| *Tubular proteinosis, cortex* | 0 | 0 | 0 | 0 | 0 | 0 |
| *Tubular proteinosis, medulla* | 0 | 0 | 0 | 0 | 0 | 1 |
| **Spleen** |  |  |  |  |  |  |
| *Lymphoid hyperplasia, white pulp* | 2 | 2 | 2 | 0 | 0 | 0 |
| *Extramedullary hematopoiesis, red pulp* | 0 | 2 | 2 | 3 | 3 | 3 |
| **Lung** |  |  |  |  |  |  |
| *Increased mixed leukocytes, alveolar septae* | 2-3 | 2-3 | 2-3 | 2-3 | 2-3 | 2-3 |
| *Perivascular/peribronchiolar lymphocytic to lymphohistiocytic infiltrates* | 1 | 0 | 0 | 1-2 | 0 | 0 |

**Table S4:** The effect of Listeria-TT and Gemcitabine on liver functions in C57Bl/6 mice

| **Treatment** | **AST (U/L)** | **ALT (U/L)** |
| --- | --- | --- |
| Saline | 107 | 16 |
| Listeria-TT and GEM | 137 | 15 |
| Normal values | 50-100 | 25-60 |

## KEY RESOURCES TABLE

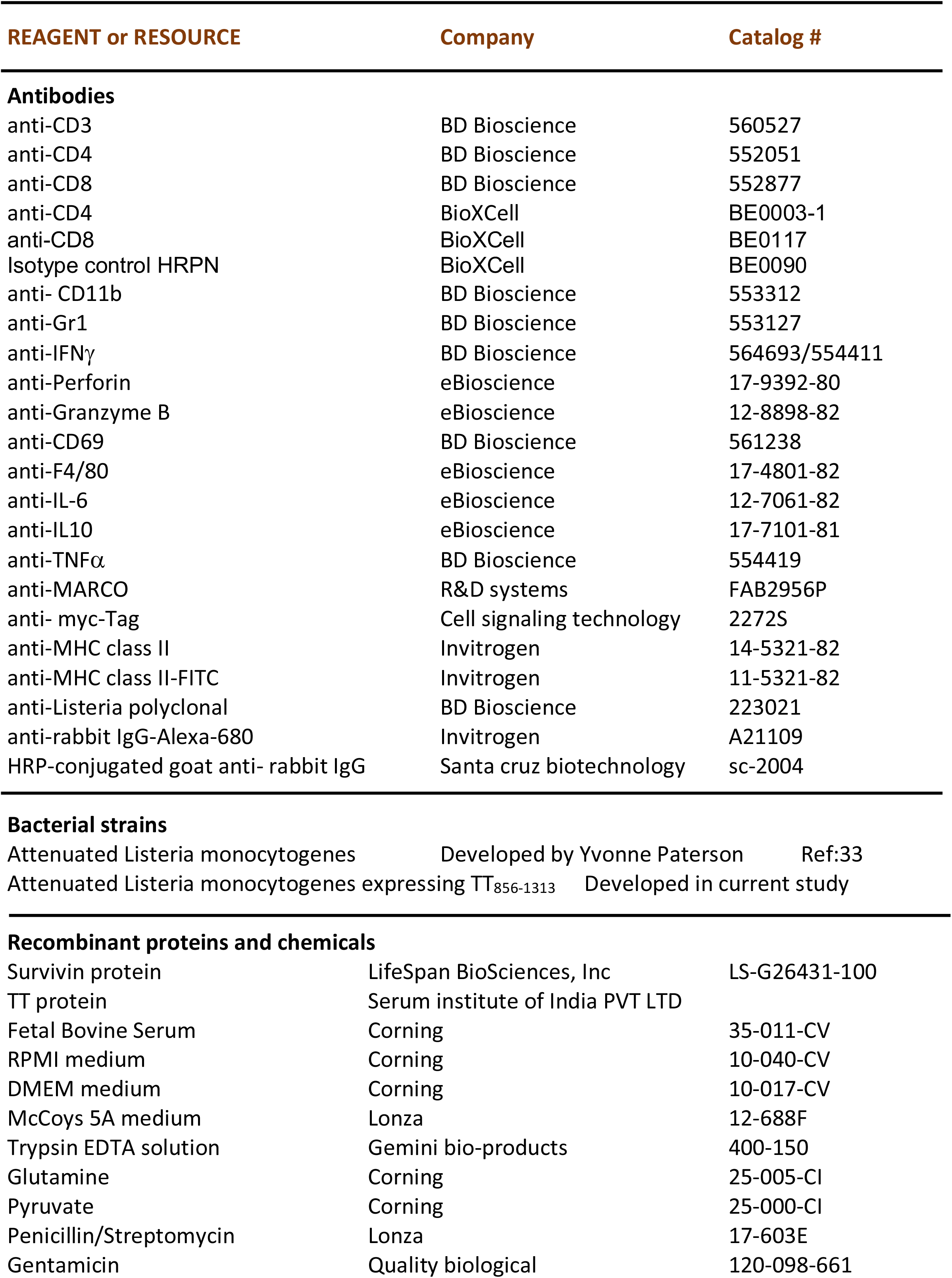
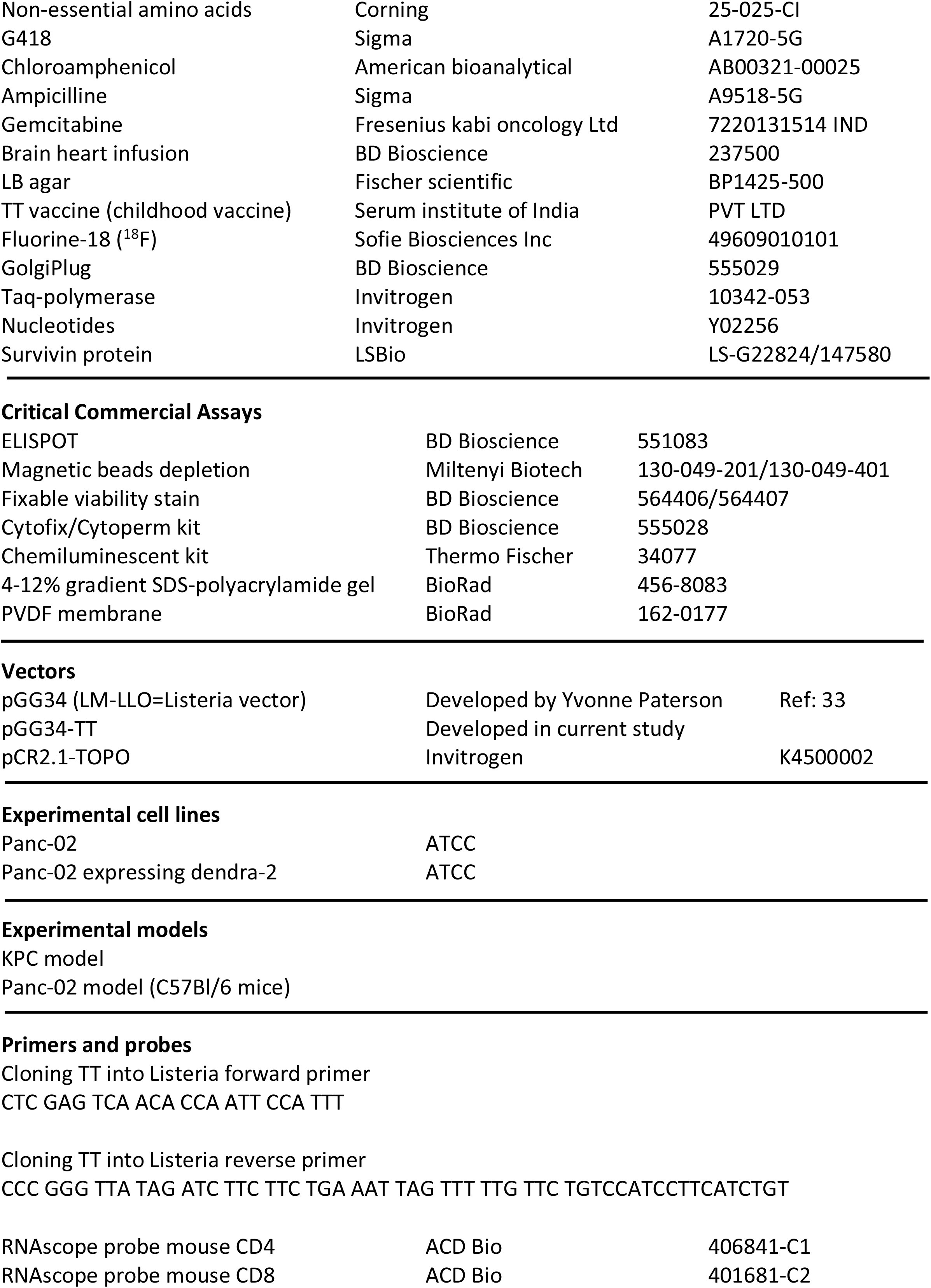
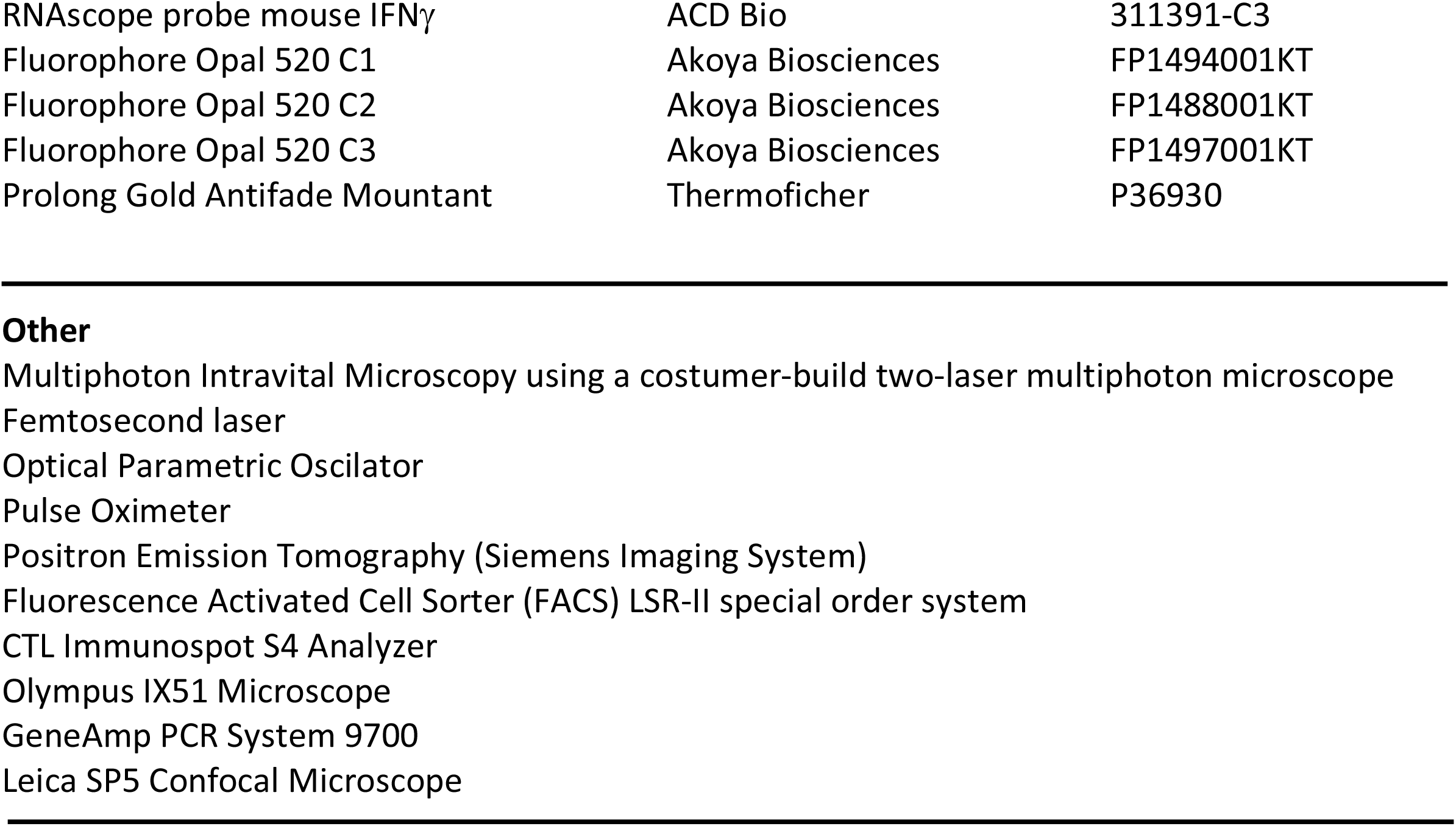

| REAGENT or RESOURCE | Company | Catalog # |
| --- | --- | --- |
| **Antibodies** |  |  |
| anti-CD3 | BD Bioscience | 560527 |
| anti-CD4 | BD Bioscience | 552051 |
| anti-CD8 | BD Bioscience | 552877 |
| anti-CD4 | BioXCell | BE0003-1 |
| anti-CD8 | BioXCell | BE0117 |
| Isotype control HRPN | BioXCell | BE0090 |
| anti- CD11b | BD Bioscience | 553312 |
| anti-Gr1 | BD Bioscience | 553127 |
| anti-IFNγ | BD Bioscience | 564693/554411 |
| anti-Perforin | eBioscience | 17-9392-80 |
| anti-Granzyme B | eBioscience | 12-8898-82 |
| anti-CD69 | BD Bioscience | 561238 |
| anti-F4/80 | eBioscience | 17-4801-82 |
| anti-IL-6 | eBioscience | 12-7061-82 |
| anti-IL10 | eBioscience | 17-7101-81 |
| anti-TNFα | BD Bioscience | 554419 |
| anti-MARCO | R&D systems | FAB2956P |
| anti- myc-Tag | Cell signaling technology | 2272S |
| anti-MHC class II | Invitrogen | 14-5321-82 |
| anti-MHC class II-FITC | Invitrogen | 11-5321-82 |
| anti-Listeria polyclonal | BD Bioscience | 223021 |
| anti-rabbit IgG-Alexa-680 | Invitrogen | A21109 |
| HRP-conjugated goat anti- rabbit IgG | Santa cruz biotechnology | sc-2004 |
| **Bacterial strains** |  |  |
| Attenuated Listeria monocytogenes | Developed by Yvonne Paterson | Ref:33 |
| Attenuated Listeria monocytogenes expressing TT<sub>856-1313</sub> | Developed in current study |  |
| **Recombinant proteins and chemicals** |  |  |
| Survivin protein | LifeSpan BioSciences, Inc | LS-G26431-100 |
| TT protein | Serum institute of India PVT LTD |  |
| Fetal Bovine Serum | Corning | 35-011-CV |
| RPMI medium | Corning | 10-040-CV |
| DMEM medium | Corning | 10-017-CV |
| McCoy's 5A medium | Lonza | 12-688F |
| Trypsin EDTA solution | Gemini bio-products | 400-150 |
| Glutamine | Corning | 25-005-CI |
| Pyruvate | Corning | 25-000-CI |
| Penicillin/Streptomycin | Lonza | 17-603E |
| Gentamicin | Quality biological | 120-098-661 |
| Non-essential amino acids | Corning | 25-025-CI |
| G418 | Sigma | A1720-5G |
| Chloroamphenicol | American bioanalytical | AB00321-00025 |
| Ampicilline | Sigma | A9518-5G |
| Gemcitabine | Fresenius kabi oncology Ltd | 7220131514 IND |
| Brain heart infusion | BD Bioscience | 237500 |
| LB agar | Fischer scientific | BP1425-500 |
| TT vaccine (childhood vaccine) | Serum institute of India | PVT LTD |
| Fluorine-18 (<sup>18</sup>F) | Sofie Biosciences Inc | 49609010101 |
| GolgiPlug | BD Bioscience | 555029 |
| Taq-polymerase | Invitrogen | 10342-053 |
| Nucleotides | Invitrogen | Y02256 |
| Survivin protein | LSBio | LS-G22824/147580 |
| Critical Commercial Assays |  |  |
| ELISPOT | BD Bioscience | 551083 |
| Magnetic beads depletion | Miltenyi Biotech | 130-049-201/130-049-401 |
| Fixable viability stain | BD Bioscience | 564406/564407 |
| Cytofix/Cytoperm kit | BD Bioscience | 555028 |
| Chemiluminescent kit | Thermo Fischer | 34077 |
| 4-12% gradient SDS-polyacrylamide gel | BioRad | 456-8083 |
| PVDF membrane | BioRad | 162-0177 |
| Vectors |  |  |
| pGG34 (LM-LLO=Listeria vector) | Developed by Yvonne Paterson | Ref: 33 |
| pGG34-TT | Developed in current study |  |
| pCR2.1-TOPO | Invitrogen | K4500002 |

Experimental cell lines
|  |  |
| --- | --- |
| Panc-02 | ATCC |
| Panc-02 expressing dendra-2 | ATCC |
| RNAscope probe mouse CD4 | ACD Bio | 406841-C1 |
| RNAscope probe mouse CD8 | ACD Bio | 401681-C2 |
| RNAscope probe mouse IFNγ | ACD Bio | 311391-C3 |
| Fluorophore Opal 520 C1 | Akoya Biosciences | FP1494001KT |
| Fluorophore Opal 520 C2 | Akoya Biosciences | FP1488001KT |
| Fluorophore Opal 520 C3 | Akoya Biosciences | FP1497001KT |
| Prolong Gold Antifade Mountant | Thermofisher | P36930 |

## Notes

### Competing Interest Statement

The authors have no competing interest, except Gravekamp. A patent of the Listeria-recall antigen concept is licensed to Loki Therapeutics

### Summary of Updates

We have added data survival, CD4 and CD8 T cell depletion, colonization of Listeria in human tumors, infecting and killing human tumors, expressing TT in tumors, RNAscope, RNAseq

